# Identification of SARS-CoV-2 Mpro inhibitors through deep reinforcement learning for de novo drug design and computational chemistry approaches

**DOI:** 10.1101/2024.02.12.579977

**Authors:** Julien Hazemann, Thierry Kimmerlin, Roland Lange, Aengus Mac Sweeney, Geoffroy Bourquin, Daniel Ritz, Paul Czodrowski

**Affiliations:** Physical Chemistry, Chemistry Department, Johannes Gutenberg University, Duesbergweg 10-14, 55128 Mainz, Germany; Drug Discovery Chemistry, Idorsia Pharmaceuticals Ltd., Hegenheimermattweg 91, 4123 Allschwil, Switzerland

**Author notes:** Footnotes relating to the title and/or authors should appear here.

## Abstract

Severe acute respiratory syndrome coronavirus 2 (SARS-CoV-2) has caused a global pandemic of coronavirus disease (COVID-19) since its emergence in December 2019. As of January 2024, there has been over 774 million reported cases and 7 million deaths worldwide.^[1]^ While vaccination efforts have been successful in reducing the severity of the disease and decreasing the transmission rate, the development of effective therapeutics against SARS-CoV-2 remains a critical need.[2] The main protease (Mpro) of SARS-CoV-2 is an essential enzyme required for viral replication and has been identified as a promising target for drug development. In this study, we report the identification of novel Mpro inhibitors, using a combination of deep reinforcement learning for de novo drug design with 3D pharmacophore/shape-based alignment and privileged fragment match count scoring components followed by hit expansions and molecular docking approaches. Our experimentally validated results show that 3 novel series exhibit potent inhibitory activity against SARS-CoV-2 Mpro, with IC50 values ranging from 1.3 uM to 2.3 uM and a high degree of selectivity. These findings represent promising starting points for the development of new antiviral therapies against COVID-19.

## Introduction

The COVID-19 pandemic has highlighted the urgent need for effective therapies to treat the disease caused by the SARS-CoV-2 virus. One promising approach is the development of small molecule inhibitors targeting the main protease of the virus, also known as Mpro. This enzyme plays a critical role in the replication of the virus, making it an attractive target for drug discovery.^[3],[4]^

In this context, the Covid Moonshot effort is a collaborative and open-science initiative^[5]^ aimed at discovering drugs to target the SARS-CoV-2 main protease. The campaign utilized crowdsourcing, high-throughput structural biology, machine learning (ML), and molecular simulations to identify new chemical series with potent nanomolar activity. Through this effort, a comprehensive understanding of the structural plasticity of the main protease was obtained, along with extensive structure-activity relationships for multiple chemotypes and a large dataset of biochemical activities. Notably, the initiative achieved a significant milestone by openly sharing all compound designs, crystallographic data, assay data, and synthesized molecules, creating a large knowledgebase for future anti-coronavirus drug discovery that is accessible and free from intellectual property restrictions. The Covid Moonshot efforts have been highly beneficial to our research. They discovered several potent chemical series and provided valuable insights into the structure of the SARS-CoV-2 main protease. The open sharing of data has created a valuable resource for our anti-coronavirus drug discovery and helped accelerating our progress.

Moreover, in recent years, ML methods have gained popularity in the field of drug discovery due to their ability to accelerate the identification of novel drug candidates.^[6],[7]^ One such approach is deep reinforcement learning for de novo drug design, which involves a trained neural network, scoring components and a reward function. The trained neural network or Deep Generative Model (DGM) generates novel chemical structures, the generated molecules are ranked by scoring components and a reward function combines all the scores. The generative model iteratively adapts its policy to maximize the reward and therefore generates, over time, molecules with desirable properties, such as high affinity, good pharmacokinetics, and low toxicity.

Researchers have applied a similar approach to the identification of epidermal growth factor (EGFR) inhibitors, using a generative model pre-trained with ChEMBL[8] molecules and an EGFR predictive ML model. The generated structures were experimentally validated and led to the identification of several new bioactive molecules.^[9]^ Another similar approach employing Reinforcement Learning has been applied retrospectively to demonstrate that a DGM could generate active molecules where these molecules were not included in the training dataset.^[10]^ Finally, de novo design approaches, without reinforcement learning were applied to discover new SARS-COV-2 Mpro inhibitors for which two approaches were experimentally validated^[11,12]^ while two other were not.^[13,14]^

In this study, we have used REINVENT 2.0 an AI tool for de novo drug design developed by Thomas Blaschke et al. The generative engine of the tool is directed by a Reinforcement Learning (RL) module. To generate compounds from a specific area of the chemical space, REINVENT 2.0 employs a composite scoring function consisting of different user-defined components.^[15]^ The tool has some readily available scoring components such as Quantitative Estimate of Drug-likeness QED^[16]^ score, Tanimoto similarity or Jaccard distance. It also offers two scenarios: exploration or exploitation. In the exploration scenario, the pre-trained DGM is used as such to explore, in a RL fashion, the chemical space for the generation of new desired molecules. In the exploitation scenario, the DGM is retrained with a set of reference inhibitors to exploit, in a RL fashion, a known chemical space for the generation of desired molecules.

Potential new inhibitors for the Mpro enzyme were generated using the exploration and exploitation scenarios. Two additional components were developed: a 3D pharmacophore/shape-alignment scoring component and a Substructure Match Count scoring component. Deep generative models (DGMs) were trained using reinforcement learning in both scenarios, and the trained DGMs were utilized to generate a large set of potential Mpro inhibitors. Various filters were applied to select promising candidates. The molecules surviving all filters were docked into Mpro, ranked and filtered. At this point, compounds commercially available molecules were ordered, while the remaining compounds underwent further docking and scoring, leading to the selection of additional compounds. Finally, another batch of compounds was ordered, synthesized, and delivered based on budget and synthesizability considerations. 16 out of 17 compounds were delivered, assessed for their potential to inhibit Mpro and 7 compounds demonstrated promising inhibitory activity. To further explore their potential and increase their novelty, additional investigations were conducted using hit expansion and molecular docking techniques with Schrödinger Glide^[17]^. However, due to a late delivery, it was not possible to conduct the hit expansion for one compound despite being bioactive.

As a result of these efforts, four new Mpro inhibitors were identified. One of them underwent optimization using a 3D-structure based approach, while another one was optimized using a classical Structure Activity Relation (SAR) approach. The affinities of the compounds were measured by a Mpro FRET IC50 assay, with values ranging from 1.3 uM to 2.3 uM. However, one of the four compounds exhibited close similarity to known Mpro inhibitors and was therefore excluded from further consideration.

Based on these findings, the three remaining compounds were selected as starting points for additional refinement in the Idorsia hit-to-lead pipeline.

## Results and Discussion

### AI generated molecules

Both the exploration and the exploitation scenarios were applied for the generation of potential new Mpro inhibitors. QED and Jaccard distance were employed, and two additional scoring components were developed, a component for a 3D pharmacophore/shape-alignment (PheSA) scoring and a component for Substructure Match Count (SMC) scoring. PheSA^[18]^ which stands for Pharmacophore Enhanced Shape Alignment is an internally developed tool at Idorsia.

In the first scenario, the original generative model, trained on large set of diverse molecules, is subjected to Transfer Learning. A small dataset of known Mpro inhibitors was used to retrain this generative model. This focused generative model allows then the exploitation mode and is capable to generate compounds from the known Mpro chemical space. The focused agent is then subjected to Reinforcement Learning with the four components described above.

In the second approach, the generative model is used in full exploration mode which means that the original generative model was applied without being retrained. This original generative model capable to generate diverse molecule is subjected to Reinforcement Learning with the same four components described above.

Once trained in the TL loop, the DGMs of both scenarios were used to equally sample a large set of potential new Mpro inhibitors and, to enrich the generated molecule dataset with compounds possessing a high likelihood of desired properties, the following selection workflow was applied:

- Initial sampling: 409,600 molecules
- Molecule sanitizer (valid smiles, duplicate filter, and SMARTS filters): 84,162 molecules
- QED > 0.5: 80,011 molecules
- SMC > 0.5: 44,377 molecules
- Jaccard distance > 0.5: 43,494 molecules
- PheSA score > 0.63: 3,119 molecules
- Bioactivity ML model predictions > 0.30: 1,736 molecules
- Docking: 188 molecules
- Synthesizability assessment: 60 molecules
- Commercially available (Enamine REAL space): 7 molecules
- Consensus scoring: 10 molecules

The post-filtering process led, after the Bioactivity ML model prediction filter, to two smaller datasets of 1,007 molecules for the exploration mode and 729 for the exploitation mode. The two datasets were compared to each other as follows: a Tanimoto Nearest Neighbor Analysis was performed on both datasets; exploitation and exploration. Each molecule of each dataset was compared with each molecule of a Mpro reference compound dataset (DGM-TL) dataset. Each comparison is based on the Tanimoto similarity coefficient (1-Jaccard distance) and allows the identification of the most similar molecule in the DGM-TL. The similarity values were then binned with a range of 0.05 and a distribution bar chart was plotted (Fig. 1) with the python library Seaborn.^[19]^ This analysis gives a solid understanding of how similar or dissimilar a dataset compared to a reference dataset looks like Figure 1 shows that the exploration dataset exhibits a greater abundance of dissimilar compounds when compared to the DGM-TL compounds, highlighting the capacity of the exploration mode to generate more novel compounds. This discrepancy can be attributed to the absence of transfer learning in the exploration approach, which encourages greater diversity in the generated molecules. Conversely, the exploitation mode involves retraining the generative model on the Mpro chemical space, resulting in a reduced potential for novelty. In this mode, the generative model predominantly produces molecules that align with its retraining, limiting the exploration of new chemical space.

**Fig. 1.**
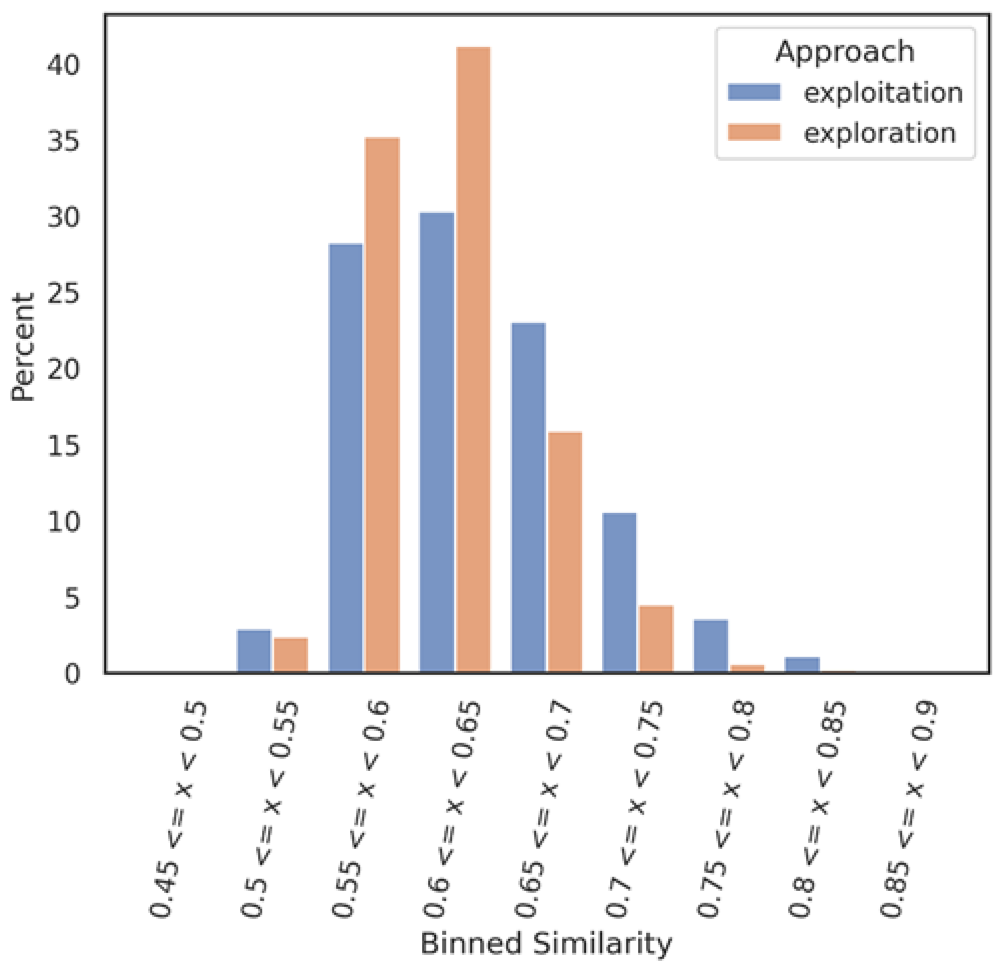
Nearest Neighbour similarity of the exploration and exploitation datasets against the DGM-TL dataset. Each subset (exploration and exploitation) was normalized independently.

Then, a Tanimoto Self-Nearest Neighbor Analysis was performed on both datasets, exploitation, and exploration respectively. Each molecule of each dataset was compared against each other molecules of the same dataset to evaluate the self-diversity of the dataset. Each comparison is based on the Tanimoto similarity coefficient (1-Jaccard distance) and allows to identify the most similar molecule within the same training set. The similarity values were then binned with a range of 0.05 and a distribution bar chart was plotted with the python library Seaborn. This analysis allows to assess the self-diversity of a dataset.

Figure 2 illustrates a distribution shift between the two datasets, revealing an interesting finding. The exploitation datasets demonstrate a reasonable level of self-diversity, while surprisingly, the exploration mode appears to have achieved a lower level of self-diversity. One potential explanation for this unexpected result is that the exploration generative model tends to generate compounds that are dissimilar to the Mpro DGM-TL dataset but exhibit self-similarity. However, further investigation is required to confirm this hypothesis and gain a better understanding of the underlying factors at play.

**Fig. 2.**
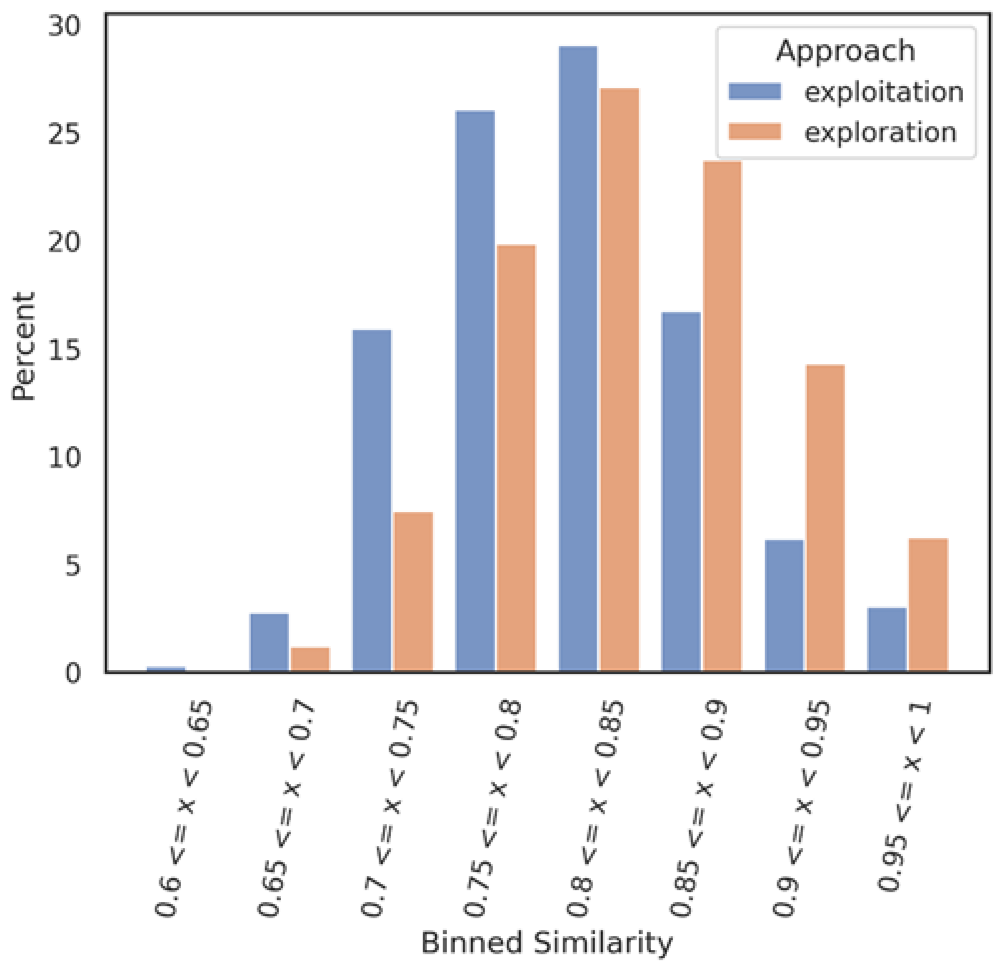
In orange, Self-Nearest Neighbor similarity of the exploration dataset against itself. In blue, Self-Nearest Neighbor similarity of the exploitation dataset against itself. Each subset (exploration and exploitation) was normalized independently.

At this stage, 1,007 molecules from the exploration approach and 729 molecules from the exploitation approach survived all the filters mentioned above. The molecules were docked into Mpro with BiosolveIT SeeSAR^[20]^. After inspection of the docking poses, 188 molecules were considered interesting.

The remaining molecules were docked again into Mpro with the open-source software Gold^[21]^, and Aizynthfinder was then employed to assess their synthesizability. 7 compounds which were found to be commercially available were directly selected and ordered. Finally, a consensus scoring, budget considerations and synthesizability constraints led to a selection of 10 additional compounds. In total 17 compounds were ordered and 16 could be synthesized, delivered, and biologically assessed.

### Biological results

The 16 AI generated, synthesized, and delivered molecules were measured in Mpro FRET (1uM DTT), Cathepsin L FRET, Mpro MCA and for some cases in SPR (Biacore) assays. As depicted in Fig. 3, 6 hits belonging to 3 clusters were identified. Cluster 1 (piperazine) showed activities between 3.3 to 63.5 uM. For cluster 2 (cyclized urea) the Mpro FRET IC50 could not be determined but a Kd of 185 and 368 uM could be measured by SPR demonstrating a weak binding affinity. Finally, the single compound in cluster 3 (N-benzoimidazol-1-yl-acetamide) showed an IC50 of 22.6 uM. Compound 21 depicted in the SI table 2 and displaying a Mpro FRET IC50 of 66.9 uM was not further considered due to a late delivery from our provider. All compounds (the 6 hits described above, compound 21 and the inactive compounds) and their biological data are shared in the supplementary info table 1.

**Table 1.** Delivered AI molecules and their biological data

| Structure | Compound # | Mpro_FRET_DTT_IC50 [umol/l] | Mpro_Biacore_KD [uM] | CathL_FRET_IC50 [umol/l] | Mpro_MCA_IC50[umol/l] |
| --- | --- | --- | --- | --- | --- |
|  | 1 | 3.27 |  | >125 | >125 |
|  | 2 | 63.0 |  | >125 | >125 |
|  | 3 | 63.5 |  | >125 | >125 |
|  | 4 | NA | 185 | >125 | >125 |
|  | 5 | NA | 368 | >125 | >125 |
|  | 6 | 22.6 | 32.8 | 103 | >125 |
|  | 13 | >125 |  | >125 | >125 |
|  | 14 | >125 | >400 | >125 |  |
|  | 15 | >125 | >400 | >125 |  |
|  | 16 | >125 |  | >125 | >125 |
|  | 17 | >125 |  | >125 | >125 |
|  | 18 | >125 |  | >125 | >125 |
|  | 19 | >125 |  | >125 | >125 |
|  | 20 | >125 |  | >125 | >125 |
|  | 21 | 66.9 |  | >125 | >125 |
|  | 22 | >125 |  | >125 | >125 |

**Table 2.**
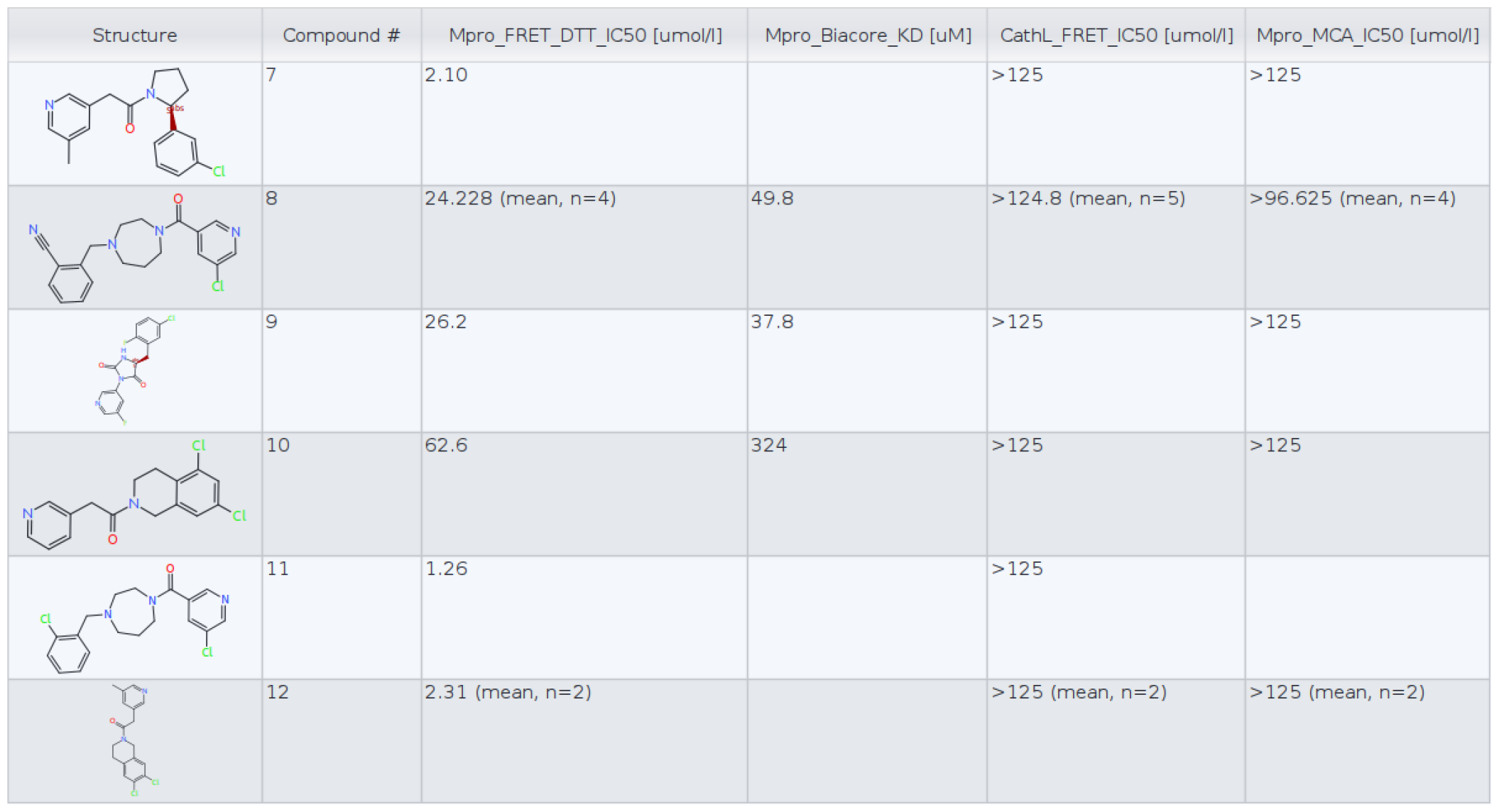
Compounds identified in the frame of the hit expansion and follow-up optimized compounds

| Structure | Compound # | Mpro_FRET_DTT_IC50 [umol/l] | Mpro_Biacore_KD [uM] | CathL_FRET_IC50 [umol/l] | Mpro_MCA_IC50 [umol/l] |
| --- | --- | --- | --- | --- | --- |
|  | 7 | 2.10 |  | >125 | >125 |
|  | 8 | 24.228 (mean, n=4) | 49.8 | >124.8 (mean, n=5) | >96.625 (mean, n=4) |
|  | 9 | 26.2 | 37.8 | >125 | >125 |
|  | 10 | 62.6 | 324 | >125 | >125 |
|  | 11 | 1.26 |  | >125 |  |
|  | 12 | 2.31 (mean, n=2) |  | >125 (mean, n=2) | >125 (mean, n=2) |

**Table 3.** X-ray data collection and processing for compounds 7, 8, 11 and 12

| Compound | Cpd 7 | Cpd 8 | Cpd 11 | Cpd 12 |
| --- | --- | --- | --- | --- |
| Diffraction source | Swiss Light Source, X06DA (PXIII) | Swiss Light Source, X06SA (PXI) | Swiss Light Source, X06DA (PXII) | Swiss Light Source, X06DA (PXIII) |
| Wavelength (Å) | 1.00 | 1.00 | 1.00 | 1.00 |
| Temperature (K) | 100 | 100 | 100 | 100 |
| Detector | Eiger 16M | Eiger 16M | Eiger 16M | Eiger 16M |
| Crystal to detector distance (mm) | 135 | 135 | 150 | 135 |
| Total rotation range (°) | 180 | 360 | 180 | 360 |
| Rotation per image (°) | 0.25 | 0.20 | 0.25 | 0.20 |
| Exposure time per image (s) | 0.120 | 0.015 | 0.250 | 0.125 |
| Space group | P 21 21 21 | P 21 21 21 | P 21 21 21 | P 21 21 21 |
| a, b, c (Å) | 68.1 102.0 104.7 | 68.0 100.3 104.6 | 67.8 101.2 103.8 | 68.0 101.8 105.0 |
| $\alpha$ $\beta$, $\gamma$ (°) | 90 90 90 | 90 90 90 | 90 90 90 | 90 90 90 |
| Mosaicity (°) | 0.19 | 0.16 | 0.12 | 0.08 |
| Resolution range (Å)<br>(highest shell) | 51.0-1.31<br>(1.40-1.31) | 56.76-1.59<br>(1.76-1.59) | 72.47-1.34<br>(1.42-1.34) | 49.80-1.30<br>(1.33-1.30) |
| Total no. of reflections | 777335 (34443) | 538998 (26606) | 897063 (36301) | 1190708 (84732) |
| No. of unique reflections | 142995 (7149) | 81101 (4055) | 136333 (6816) | 179022 (13124) |
| Completeness (%)<br>(spherical) | 81.8 (21.9) | 73.2 (15.9) | 84.3 (25.4) | 99.8 (99.7) |
| Completeness (%)<br>(ellipsoidal) | 93.7 (52.1) | 94.8 (61.4) | 96.1 (66.4) | Not calculated |
| Multiplicity | 5.4 (4.8) | 6.6 (6.6) | 6.6 (5.3) | 3.4 (3.3) |
| $\langle I/\sigma(I) \rangle$ from merged data | 7.6 (1.6) | 13.9 (1.8) | 15.4 (1.7) | 10.3 (1.1) |
| CC<sub>1/2</sub> (highest shell) | (50.2) | (54.4) | (69.0) | (36.5) |

**Fig. 3.**
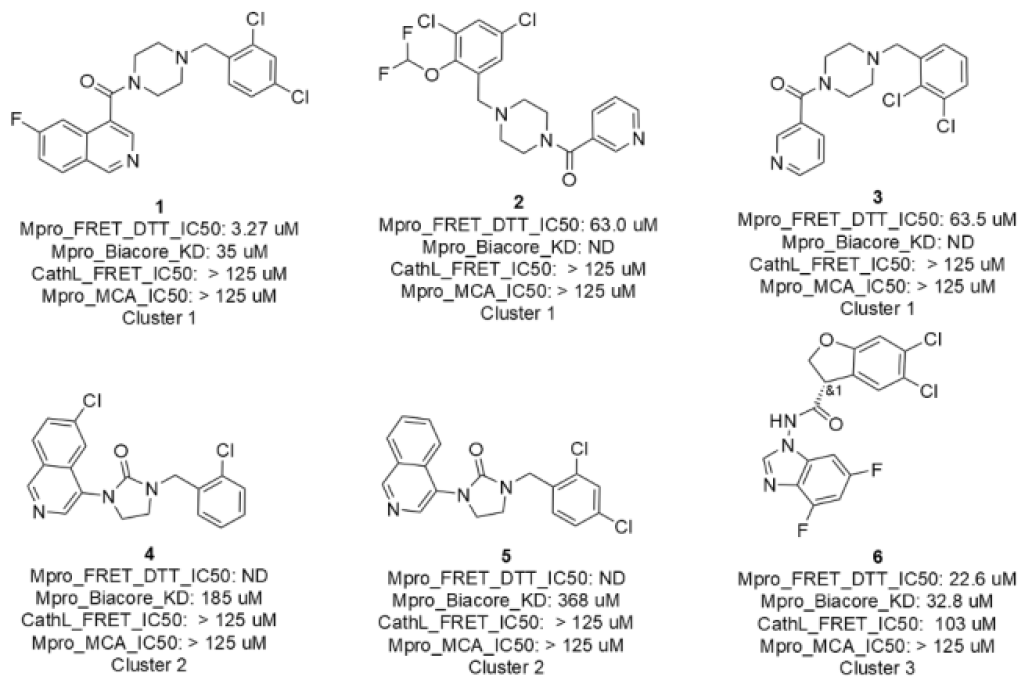
Biological results of the selected, synthesized and delivered AI generated molecules

While the 6 hits showed good to low affinity, they all, except for compounds 4 and 5 (not conclusive at this point), demonstrated a good selectivity profile as indicated by the Cathepsin L IC50 values. All hits turned up not being auto-fluorescent as demonstrated by the Mpro MCA assay.

As mentioned above, these 6 hits were categorized in 3 clusters which we defined as 3 hit series: a piperazine, a cyclized urea and a N-benzoimidazol-1-yl-acetamide series. The 2D similarity was evaluated between the 6 hits and known Mpro inhibitors from ChEMBL and Covid Moonshot. A limited novelty was observed as compared to known Mpro inhibitors, therefore, a hit expansion was performed for the 3 hit series to bring additional novelty. Analogues, based on 2D similarity, were identified in either the Chemspace^[22]^ screening compound stock or in the Idorsia corporate collection. The analogues were docked, and the best docking poses were selected. In total, 173 compounds were considered for a biological assessment. This hit expansion led to the identification of 4 interesting and diverse inhibitors (compounds 7, 8, 9 and 10) as depicted in Fig. 4. Compounds 8 and 10 were originated from the hit expansion of cluster 1, compound 7 was originated from the hit expansion of cluster 2 and compound 9 was originated from the hit expansion of cluster 3. Additional Mpro inhibitors were identified with the approach but cannot be disclosed due to corporate restrictions.

**Fig. 4.**
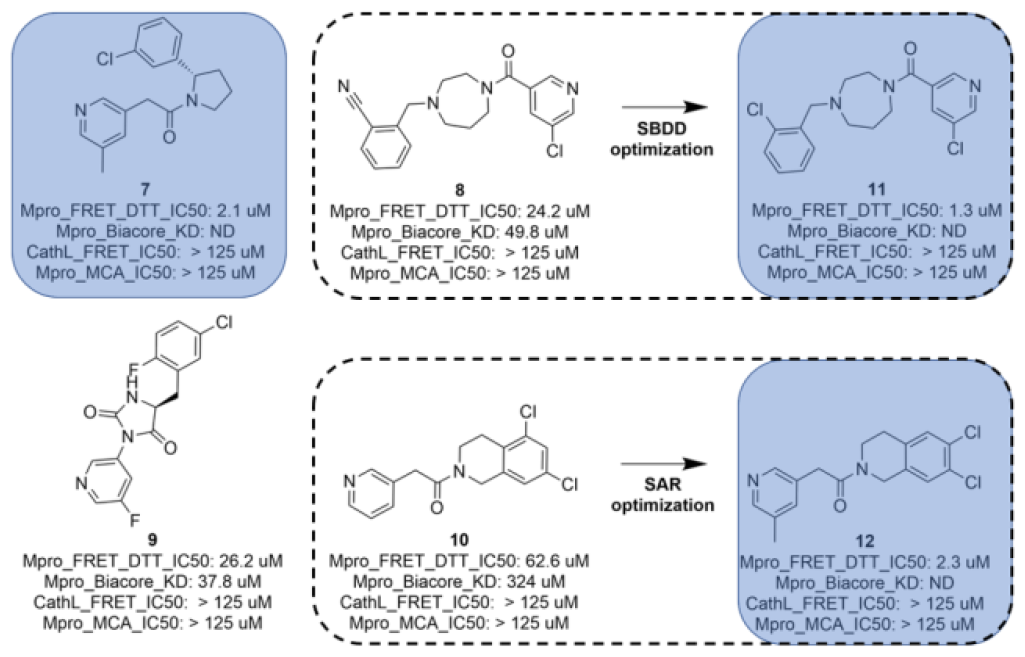
Biological results of the hit expansion and first optimization molecules

Two (compound 8 and 10) of the four inhibitors identified in the frame of the hit expansion were further optimized. The diazepane molecule (compound 8) was optimized following a structure-based approach (SBDD) while the tetrahydroisoquinoline molecule (compound 10) was improved by a standard SAR approach. This first round of optimization allowed a 28-fold improvement for the diazepane series to 1.3uM and a 27-fold improvement for the tetrahydroisoquinoline series to 2.3uM.

Eventually, the AI generated hits followed-up by hit expansion, docking and a first round of optimization for compound 8 and 10 resulted in the identification of 3 novel Mpro inhibitors, a diazepane (compound 11), a pyrrolidine (compound 7) and a tetrahydroisoquinoline (compound 12) with IC50s ranging from 1.3 to 2.3 uM. Compound 9, while better than its queries (compound 4 and 5) was considered too similar to the compounds identified in the frame of the COVID Moonshot.

### Crystal structure determination and structure-based design

The X-ray structures of SARS-CoV-2 Mpro in complex with compounds 7 and 8 were solved.

The binding mode of compound 7 is depicted in Figure 5. The following observation can be made with this figure, the m-methylpyridine sits in the S1 pocket where the pyridine nitrogen makes a key H-bond interaction with H163. The amide carbonyl makes a key interaction with the E166 backbone-NH and a water-mediated interaction with the E166 backbone-carbonyl. The m-chlorophenyl sits in the hydrophobic S2 pocket making face-to-edge aryl interaction with H41.

**Fig. 5.**
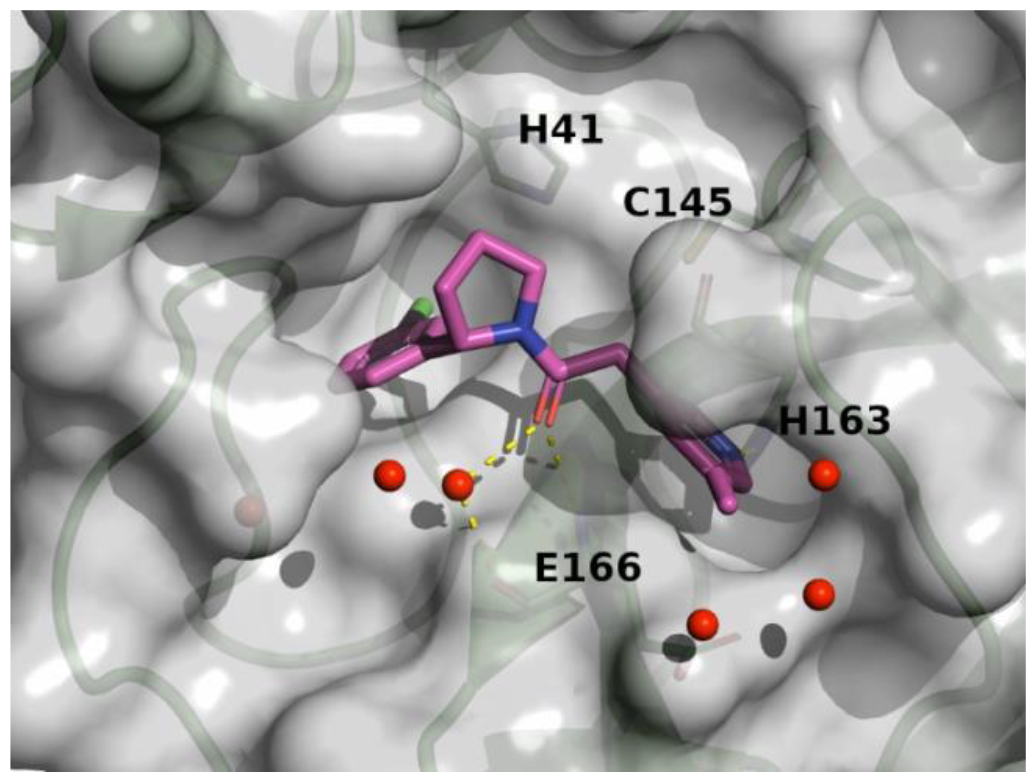
X-ray crystal structure of compound 7 in the SARS-CoV-2 Mpro active site

**Fig. 6.**
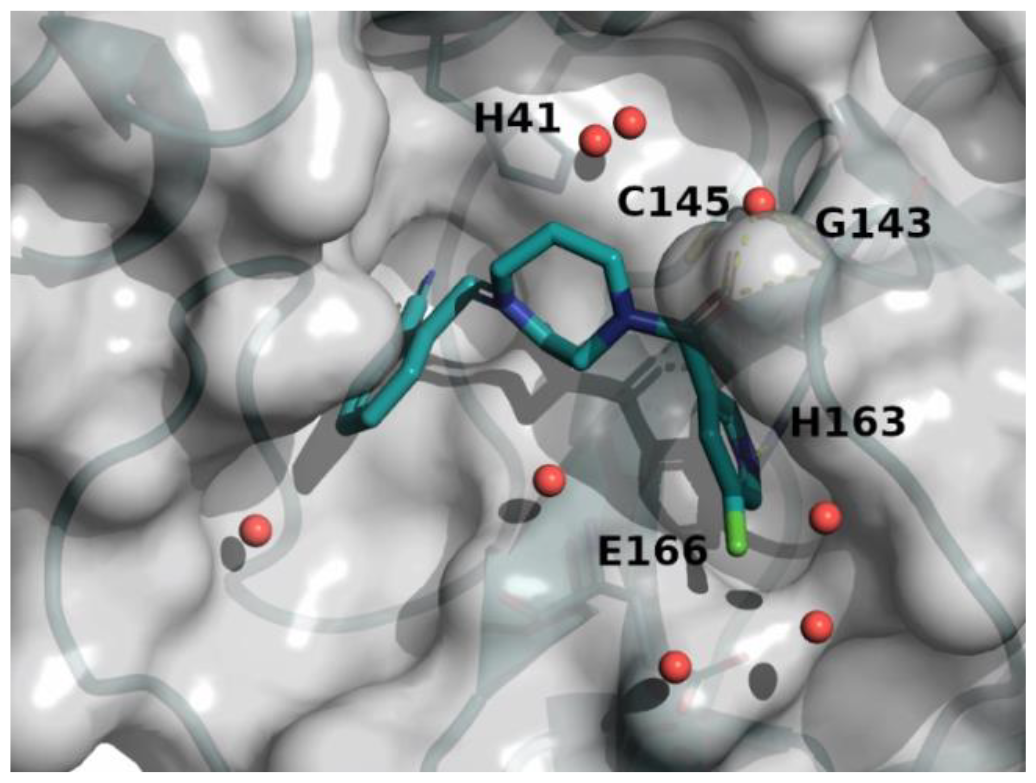
X-ray crystal structure of compound 8 in the SARS-CoV-2 Mpro active site

The binding mode of compound 8 is depicted in Figure 6. The following observation can be made with this figure: the m-chloropyridine sits in the S1 pocket where the pyridine nitrogen makes a key H-bond interaction with H163. The amide carbonyl makes a key interaction with the G143. The m-chlorophenyl sits in the hydrophobic S2 pocket making face-to-edge aryl interaction with H41.

As depicted in Figure 7, the 3D structure observations allowed to understand the importance of a chlorine atom in the S2 pocket. Indeed, for example, compound 7 (magenta) with an Mpro FRET IC50 of 5.6uM or the COVID Moonshot compound EDJ-MED-92e193ae-1 (yellow) with an Mpro FRET IC50 of 0.230uM for which available crystal structures were aligned with the crystal structure of compound 8 (cyan). This picture shows the possibility of replacing the cyano moiety of compound 8 by a chlorine in ortho position. This SBDD suggestion was then synthesized and led to compound 11 with an Mpro FRET IC50 of 1.3uM (28-fold improvement)

**Fig. 7.**
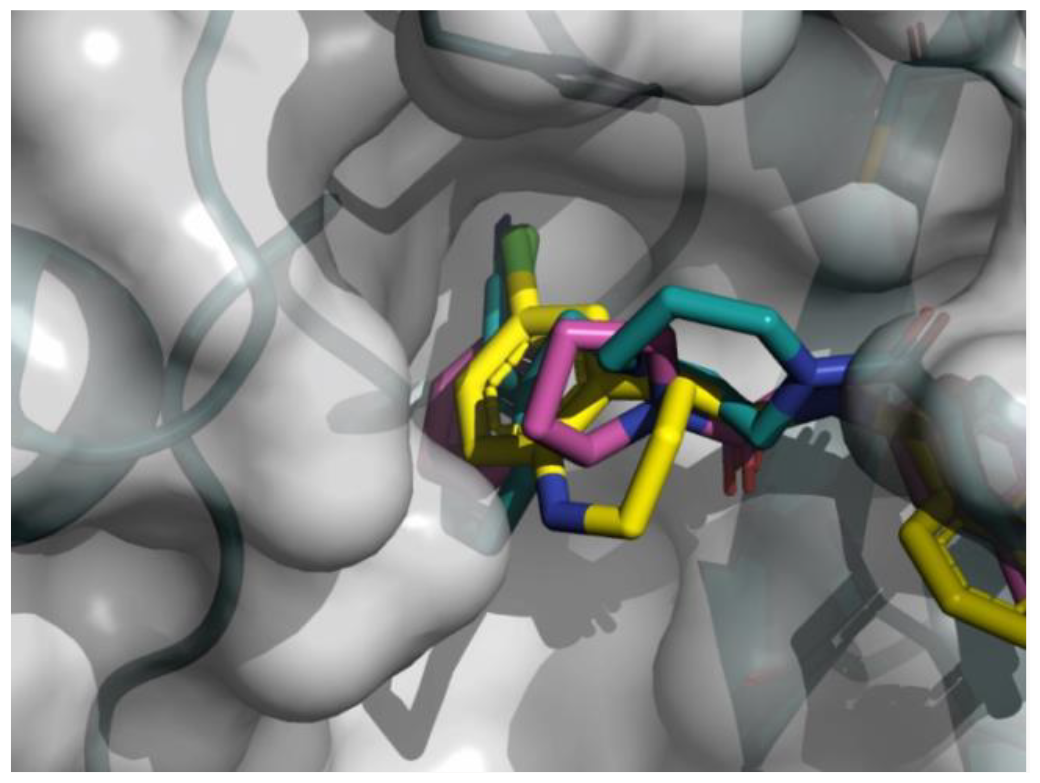
X-ray crystal structure of compound 8, 7 and COVID Moonshot EDJ-MED-92e193ae-1 superimposed in the SARS-CoV-2 Mpro active site

The X-ray structures of SARS-CoV-2 Mpro in complex with compounds 11 and 12 were solved.

The binding mode of compound 11 is depicted in Figure 8. The following observation can be made with this figure: the m-chloropyridine sits in the S1 pocket where the pyridine nitrogen makes a key H-bond interaction with H163. The amide carbonyl makes a key interaction with the G143. The m-cyanophenyl sits in the hydrophobic S2 pocket making face-to-edge aryl interaction with H41.

**Fig. 8.**
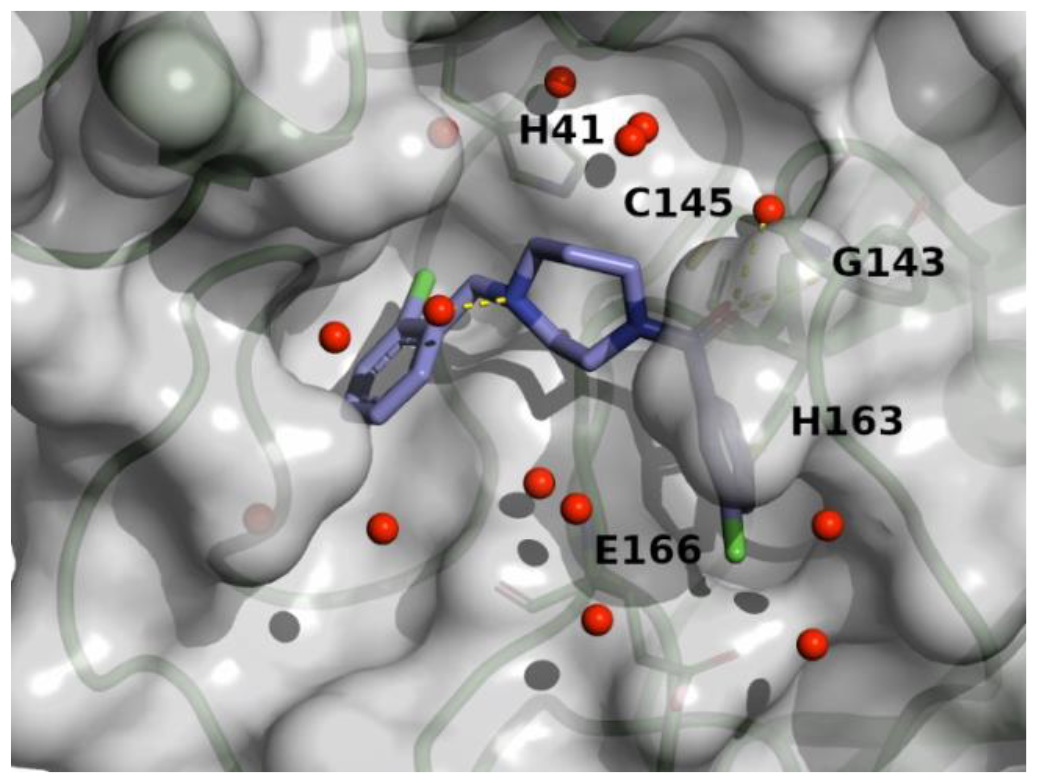
X-ray crystal structure of compound 11 in the SARS-CoV-2 Mpro active site

**Fig. 9.**
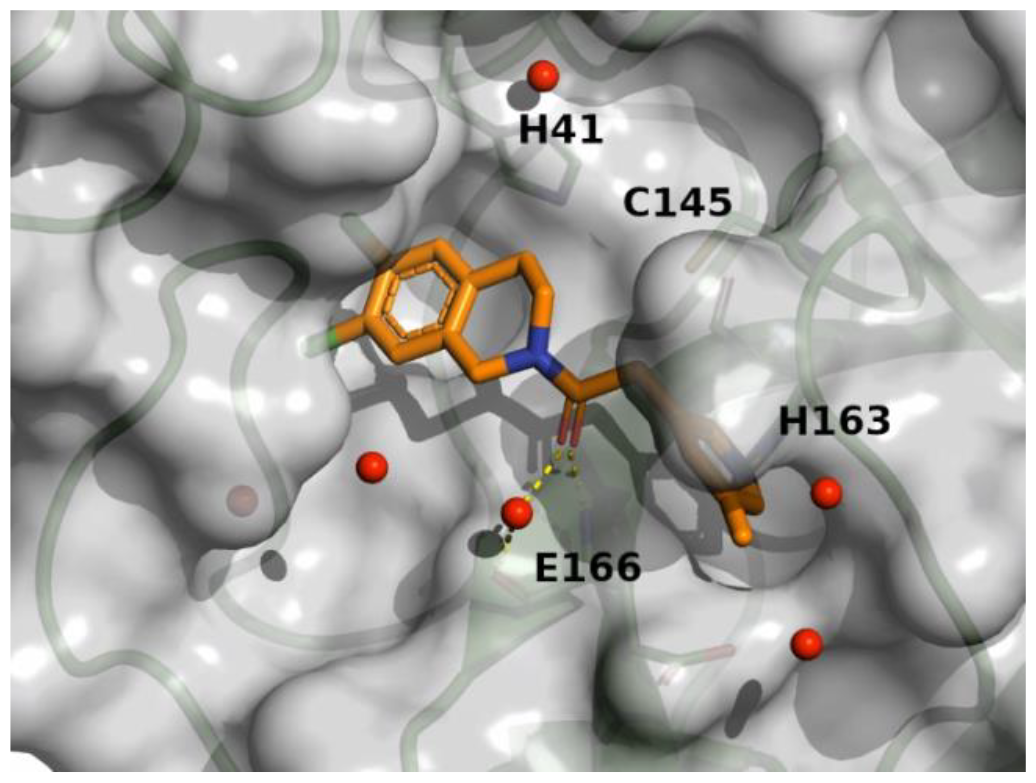
X-ray crystal structure of compound 12 in the SARS-CoV-2 Mpro active site

**Figure 10.**
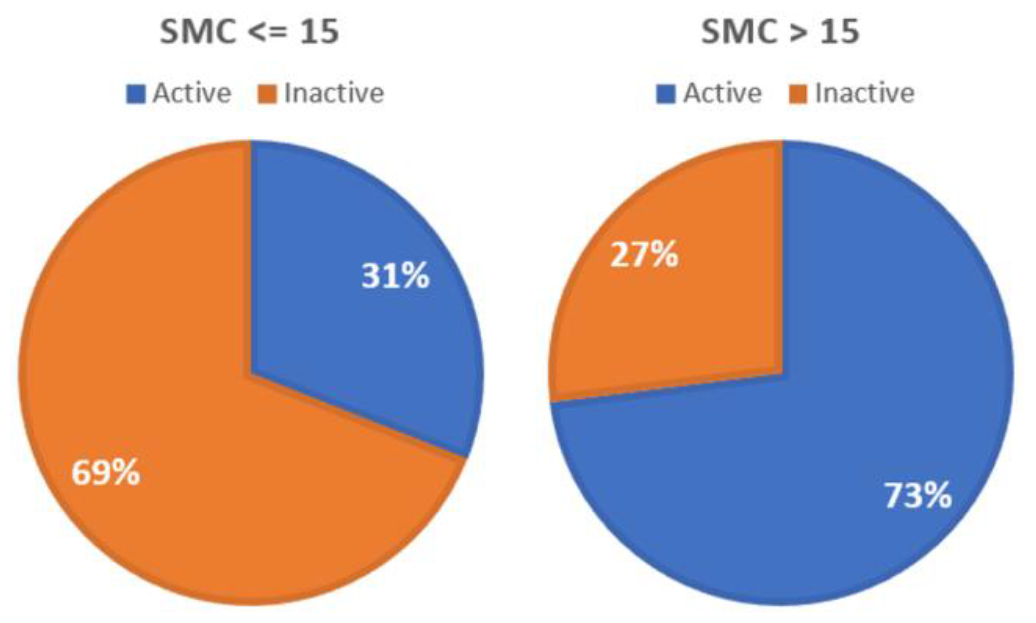
The pie charts represent the proportion of active versus inactive molecule for molecules with either a SMC <= 15 (n=815) or a SMC > 15 (n=125)

## Discussion

In this study, we have used a generative model associated with scoring components for reinforcement learning. The prior (generative model) was originally trained on smiles from the ChEMBL database. The following scoring components were employed: a 3D pharmacophore/shape alignment component, a Jaccard distance component, a Substructure Match Count scoring component and QED scoring component. These scoring components are crucial parts of the reinforcement learning, as they allow the generative model to create molecules with desired properties.

We will discuss in the following some aspects of the aforementioned workflow. On the one hand, we screened for compounds similar to known Mpro inhibitors by making use of a pharmacophore search and a substructure match count. Subsequently, potential new Mpro inhibitors very similar compounds to known Mpro inhibitors were filtered out. One might argue a pharmacophore screen yields far too many too similar molecules, but it would be very risky to not take such information into account. We intentionally decided not to include docking or any computationally expensive processes in the reinforcement learning loop to avoid lengthy training of the system. We, however, opted for using expensive techniques in a post-filter fashion.

Two approaches were considered. In the first approach, the agent is subjected to Transfer Learning. A small dataset of known Mpro inhibitors is used to retrain the prior. This focused agent allows then the exploitation mode and is capable to generate compounds from the Mpro chemical space immediately. The agent was trained in Reinforcement Learning with the four components described above. In the second approach, the generative model was used in exploration mode which means that the prior was used without being retrained and the agent was then trained in Reinforcement Learning with the same four components described above.

Both trained systems were used to generate twice 200,000 molecules. The 400,000 molecules were filtered in order to decrease the search space. The filtered compounds of both approaches were then compared in a nearest neighbor analysis against a set of known Mpro inhibitors. It demonstrates that compounds from the exploration model are more dissimilar to the Mpro known inhibitors as compared to the ones generated during the exploitation model. However, the molecules generated during the exploitation model were also fairly dissimilar to the known Mpro molecules. In a second analysis each compound of a dataset (exploration or exploitation) was compared against each other compound of the same set in a self-nearest neighbor analysis. This second analysis shows a shift as molecules generated by the exploration model are less diverse than the ones from the exploitation model. This finding is counter intuitive as we would expect the molecules generated by the exploration model more diverse, but this phenomenon can perhaps be explained by an over representation of similar molecules that are indeed dissimilar to the Mpro reference molecules. Overall, these two analyses illustrate the model and filter’s abilities to reach an interesting dissimilarity as compared to the known Mpro chemical space and to reach a fair self-diversity in both approaches.

At this stage, 1700 molecules were considered. To further reduce the number and to focus on the ones with a higher likelihood of being active, these molecules were docked against 3 Mpro complexes obtained from the COVID Moonshot research. We selected 3 different complexes to help overcome the high plasticity of Mpro^[23]^ and hypothesized that using them would help to address different shapes of the protein and therefore improve the outcome of the docking. We are aware that such a selection by far does not cover the full MPro conformation space, but it represents a good compromise between speed and accuracy.

Eventually, 7 commercially available molecules were directly ordered and delivered. 10 additional compounds were selected by employing a complementary docking procedure and taking a consensus score into account. This should further lower the bias of crystal structure selection and the choice of the docking methodology.

In total, 17 compounds were considered and 16 of them were successfully delivered. However, it was discovered that some compounds looked challenging to synthesize, despite being filtered with aizynthfinder^[24]^. This highlighted the need for additional considerations for compound synthesizability before their purchase. It is essential to ensure that the selected compounds can be synthesized efficiently, as unsuccessful synthesis can lead to waste of time, resources, and funding.

The compounds were evaluated in biological assays and 7 of them demonstrated a weak to fair affinity and a 5 of them exhibited a good selectivity for Mpro. These results were very encouraging as 44% of the delivered compounds showed some bioactivity. Except for compound 21, the molecules were categorized in three clusters/series: piperazine (compounds 1, 2 and 3), N-benzoimidazol-1-yl-acetamide (compound 6) and cyclized ureas (compounds 4 and 5). The compounds for the three series were compared with the compounds identified in the frame of the COVID Moonshot work. The piperazine core was already known from Moonshot but some novelty was brought by the amidopyridine or the dichloro-benzyl moiety (compound 1 and 2). The cyclized urea core was also known from the Moonshot but never combined with an isoquinoline and a chlorobenzyl moiety (compound 4). The N-benzoimidazol-1-yl-acetamide was known as well and the novelty was brought by a dichloro-dihydrobenzofuran (compound 6). The similarities observed with some Moonshot molecules can certainly be explained by the SMC scoring components used during the reinforcement learning and the filtering cascade.

To make these findings even more interesting and to bring more novelty, a hit expansion was performed. Molecules displaying 2D similarities with the 3 series mentioned above were identified from stock collections (Chemspace or corporate collection). These compounds were docked with hydrogen-bond constraints, the docking poses were visually inspected and the ones possessing the necessary key H-bond interactions and adding some added value in terms of novelty were selected. The compounds were then ordered and assessed biologically and led to the identification of 4 molecules: A diazepane (compound 8), a pyrrolidine (compound 7), tetrahydroisoquinoline (compound 10) and an imidazolidine-2,4-dione (compound 9). Compound 21 that was delivered at a later stage and found to be active could regrettably not be selected for hit expansion. This compound is however depicted in Supplementary Information Table 1.

Three of them, the diazepane (compound 8), the pyrrolidine (compound 7) and tetrahydroisoquinoline (compound 10) remained in our focus as they were considered novel. The diazepane (compound 8) was further optimized with a 3D structure-based approach. Indeed, the replacement of an ortho-cyanobenzyl to an ortho-chlorobenzyl in the S2 pocket taken from observations made with our solved crystal structures combined with the ones from the COVID Moonshot enabled the first optimization of compound 8 to compound 11. The tetrahydroisoquinoline (compound 10) was optimized following a classical Structure Activity Relation (SAR) approach. Both approaches led to highly interesting starting points with IC50s ranging from 1.3 to 2.3 uM.

Our work showcases the potential of combining deep reinforcement learning for de novo drug design with additional computational chemistry techniques, as we successfully identified three novel Mpro inhibitors that display high potential for further development. We believe that our stepwise approach is key for slowly but surely identify new inhibitors.

There are, however, several limitations and challenges that need to be addressed to improve the efficiency and effectiveness of RL-based de novo molecule design.

One of the main limitations is the similarity of the generated molecules with reference Mpro molecules. Since the re-trained agent of our exploitation approach learns from existing molecules, it tends to generate molecules that resemble the training set. Additionally, both for the exploitation or exploration, the SMC scoring component certainly influenced the training of the agent towards the known Mpro chemical space. As a result, the generated molecules may not be sufficiently diverse or novel, leading to limited success in identifying new lead compounds.

To overcome this issue, training the generative model without the SMC scoring component in full exploration mode might constitute an interesting perspective or considering a docking scoring component instead of the SMC scoring components might also have the potential to lead to additional novelty.

Another significant challenge is the synthetic accessibility of the generated molecules. The models may generate molecules that are structurally complex or contain building blocks that are difficult to synthesize, making them impractical for further experimental evaluation. Despite the aid of Aizynthfinder in eliminating challenging or infeasible molecules, we failed to fully consider the true availability and cost of the building blocks for the interesting molecules we considered. Indeed, Aizynthfinder allows one to make its own collection of building blocks. This stock collection is then used by the tool when it recursively breaks down the molecule into available precursors. In our situation, we used the building block collection of aggregators (regrouping many vendors) which made the approach not fully effective. Indeed, the molecules were successfully considered synthesizable by Aizynthfinder but as the building blocks were available from a plethora of vendors it made very difficult one Contract Research Organization (CRO) to make a reasonable offer. Lesson learned: make an Aizynthfinder stock from a preferred vendor/CRO and not from an aggregator. Consequently, some of the compounds that were considered turned out to be excessively costly.

## Conclusion

In summary, we have used a de novo method for the generation of molecules with the goal of identifying new SARS-COV-2 Mpro inhibitors. The method uses a reinforcement learning approach that guides the generative model in creating desired molecules. It is based on the open-source code REINVENT 2.0 an AI tool for de novo drug design. The presented method is designed to generate molecules which contain key features of known Mpro inhibitors such as 3D pharmacophore/shape and privileged fragments. Additional computational chemistry techniques, used as post filters, such as molecule sanitization, QED, SMC, Jaccard distance, PheSA, ML affinity, AiZynthfinder, molecular diversity, molecular docking, and final consensus scoring were employed to further enrich the generated molecules toward higher likelihood of being active against SARS-COV-2 Mpro.

We believe that the identification of SARS-CoV-2 Mpro inhibitors through reinforcement learning de novo design combined with 3D pharmacophore modeling, shape-based alignment, hit expansion, and molecular docking approaches represents a promising avenue for the identification of new starting points that could lead to new therapies for COVID-19. We are convinced that combining modern ML techniques together with state-of-the-art computational chemistry techniques have the potential to accelerate the identification of effective treatments for this severe disease.

The success of our workflow speaks for itself: Six AI generated primary hits demonstrated weak to decent affinity. These molecules could be easily and quickly improved by hit expansion and showed novel Mpro chemotypes.

In our case study, we show that analogues picked out in the frame of the hit expansion and further prioritized by molecular docking led to the identification of three novel inhibitors. Two of them underwent a first round of optimization in the lab.

We then additionally report the identification of three novel SARS-COV-2 inhibitors displaying inhibitory activity ranging from 1.3 uM to 2.3 uM fulfilling our initial goal.

## Acknowledgements

We are grateful to Frederick Götz, Joël Wahl, Naomi Tidten, Alexander Metz, Céline Potot, Nadia Artico, Sylvia Richard, Sylvain Regeon, Eser Ihan, Siefke Siefken, Victor Ribic, Clément Pompinau, Anastasiia Netrebchuk, Alain Chambovey, Solange Meyer, Marina Dos Santos and Laksmei Goglia for their contributions.

## Supporting Information

The authors have cited additional references within the Supporting Information.[25, 41]

## Experimental section

### Datasets

Reference Mpro inhibitors: A set of 225 Mpro inhibitors (FRET or FRET and RapidFire IC50 < 5uM) was collected from various sources (ChEMBL, COVID Moonshot, and literature). The dataset contains compounds inhibiting SARS, SARS.CoV-2 and MERS Mpro. The dataset contains covalent and noncovalent inhibitors.

ML dataset: ChEMBL and COVID Moonshot molecules with IC50 data (FRET and RapidFire assay data) were pooled together. For molecules with several measurements, a standard deviation was calculated together with the difference between the max and min value. Measurements showing a difference greater than 2 times standard deviation were discarded. The ML dataset was made up of 739 datapoints.

DGM-TL dataset: A set of 225 Mpro inhibitors (FRET or FRET and RapidFire IC50 < 5uM) previously collected was combined with 113 additional inhibitors (FRET or FRET and RapidFire 5uM < IC50 < 15uM) from various sources (ChEMBL, Covid Moonshot, and literature).

COVID Moonshot FRET activity dataset: The dataset was downloaded from https://covid.postera.ai/covid/activity_data and contains all measured compounds (24.03.2021) by FRET assay for which an IC50 value is available. The dataset contains 940 molecules.

Privileged fragments: A list of privileged fragments was assembled as described in the following SMC scoring component validation section.

Docking validation set: 25 “active” (nanomolar range) non-covalent inhibitors and 25 “inactive” compounds were randomly picked from the COVID Moonshot dataset and combined.

### RL scoring component validation

A 3D pharmacophore and shape alignments scoring component from single active conformer queries was adapted as REINVENT scoring component. This component was derived from PheSA (Pharmacophore-Enhanced Shape Alignment) a tool developed at Idorsia. The reference Mpro inhibitor dataset was used to query the PDB files and extract the active conformers of the exact same ligand of the 225 reference inhibitors. 69 active ligands were identified and extracted from PDB files. PheSA descriptors were generated for these query active conformers. The 69 inhibitors were clustered, and non-covalent representatives of each cluster were selected. In total, 23 query compounds were selected. The 23 PheSA queries (single active conformers) were used to screen the test set of 69 active and 6000 inactive compounds. Molecules with a PheSA score >= 0.63 were considered predicted “active”Results:

- Precision: 0.168
- Sensitivity: 0.532
- Kappa: 0.238
- ROC AUC: 0.88

The results above show with the precision value that for all compounds predicted active 17% were correctly predicted. The sensitivity shows that 53% of all active compounds were identified. Cohen’s kappa is a quantitative measure of reliability for two raters that are rating the same thing, correcting for how often the raters may agree by chance. A kappa of 0.24 would judge the model as fair^[25]^. Finally, the ROC (receiver operating characteristic) curve gives the performance of a classification model at all classification thresholds. An AUC of 0.5 means random prediction and the closer to 1 the AUC the better. An AUC of 0.88 is considered as excellent.^[26]^

The single conformer PheSA scoring was implemented as a new RL scoring component.

### SMC scoring component validation

The SMC component is a privileged fragment substructure match scoring component. A Matched Molecular Pair Analysis was performed on the reference Mpro inhibitor dataset. Fragments that allowed a 2-fold gain in potency were selected. Fragments containing covalent warheads were discarded. The selected fragments were then combined with the noncovalent fragment hits discovered in the frame of the Covid Moonshot crystallographic screen.^[27]^ Only fragments with Mw <= 350 g/mol were retained and duplicates were removed. In total a collection of 265 privileged fragments were obtained.

The following analysis was performed with the Knime analytical platform^[28]^ combined with python:

The COVID Moonshot FRET activity dataset was split in two groups:

- 354 active molecules <= 10uM (FRET IC50)
- 586 inactive molecules > 10uM (FRET IC50)

A substructure match was performed using these compounds against a set of selected privileged fragments. The python script counts how many fragments would match a COVID Moonshot compound.

Again, the reference compounds were split in two groups.

- 815 matching counts <= 15 fragments
- 125 matching counts > 15 fragments

As we can see from the figure below, we clearly see an enrichment of “active” molecules for compounds with a matching count > 15 fragments.

A Python script was written to allow generated molecules to be evaluated against a list of privileged fragment substructures. The script counts how many privileged fragments match a molecule. Then a membership trapezoidal scoring function is applied as depicted in figure 11.

**Figure 11.**
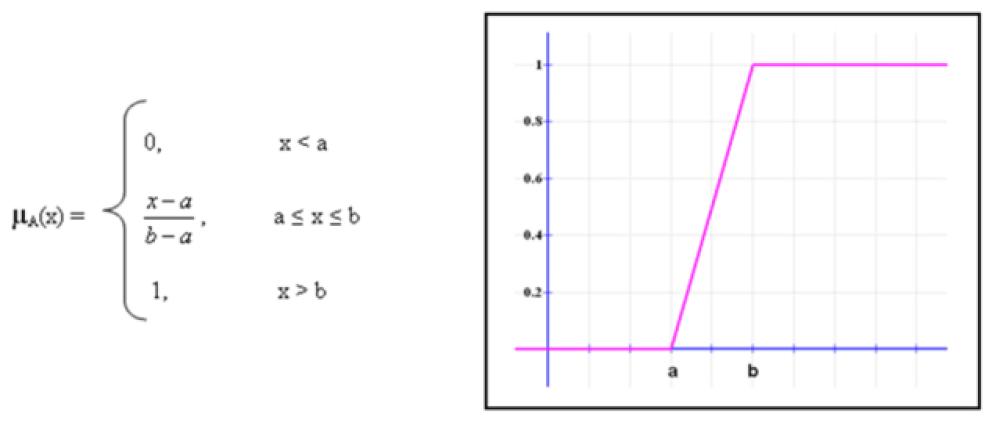
Membership trapezoidal scoring function, Substructure match count on the x axis and score on the y axis, a = 0 and b = 15

The script can score molecules based on their substructure match count against privileged fragments. The script was implemented as a scoring component in the DGM RL allowing the generation of molecules containing many privileged fragments.

### Bioactivity ML model classification

The Knime analytical platform was employed for the analysis below. The ML dataset was labeled as follow:

- Label 1 (active) <= 10000nM
- Label 0 (inactive) > 10000nM

Validation method: A random molecule (compound number 1) was picked, and its closest analogue (highest Tanimoto/Morgan FP) is identified. The identified closest analogue is assigned to compound number 2 and, again, its closest analogue (excluding its predecessor) is identified as the following compound. The same logic is applied until all molecules got a compound number. Since there is no time stamp on this dataset, a virtual chronology was simulated by sorting the compounds by similarity. Molecules were sorted from 1 to 739 and a 10-fold linear sampling was applied. Random Forest and XGBoost were evaluated with either all possible RDKit^[29]^ fingerprints (FPs) or calculated physico-chemical properties. DeepChem^[30]^ Graph Convolutional Neural Network (GCNN) was also evaluated with 25, 50, 75 and 100 epochs.

List of the best performers:

- XGBoost-Global_chiral_ECFP6
- GCNN-global_70_epochs
- XGBoost-Global_ECFP6
- RF-Global_Torsion

Then the following split was applied 80/10/10 for training/validation/test to a model ensemble (see list above)

#### ML performance on the validation set

ROC AUC: 0.881

Kappa: 0.517 (adjusted threshold)

ML performance with the test set:

- ROC AUC: 0.506
- Kappa: -0.027

The above results suggest that a ML (RF-FPs, XGBoost-FPs and GCNN) approach can predict compounds being similar with compounds in the training set (Fig. 12) but is not able to make useful prediction for dissimilar compounds (Fig. 13). In these conditions, an ML classification applied to RL or even post-processed generated molecules would be suboptimal.

**Figure 12.**
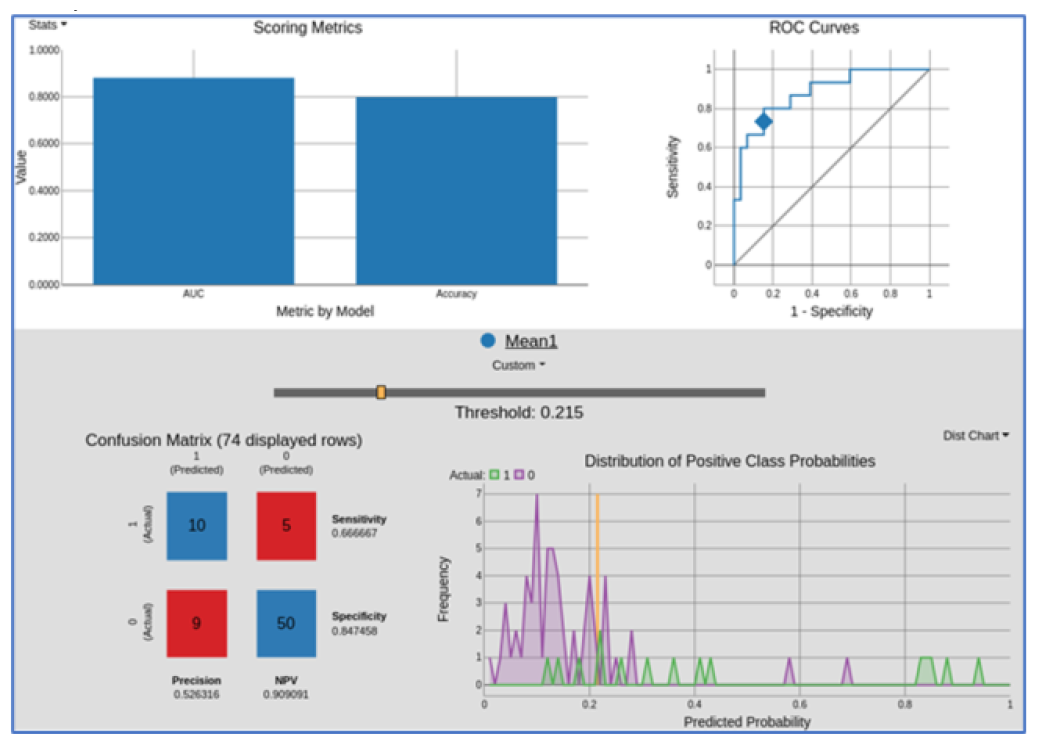
The figure was obtained from the Knime Binary Classification Inspector node. The view display performance of the RF-FPs, XGBoost-FPs and GCNN model ensemble on the validation set. On the top left the model’s statistic bar chart: AUC and overall accuracy. On the top right the ROC curve. The confusion matrix on the bottom left and the distribution of positive class probabilities on the bottom right. The statistical metrics adapt according to the threshold value.

**Figure 13.**
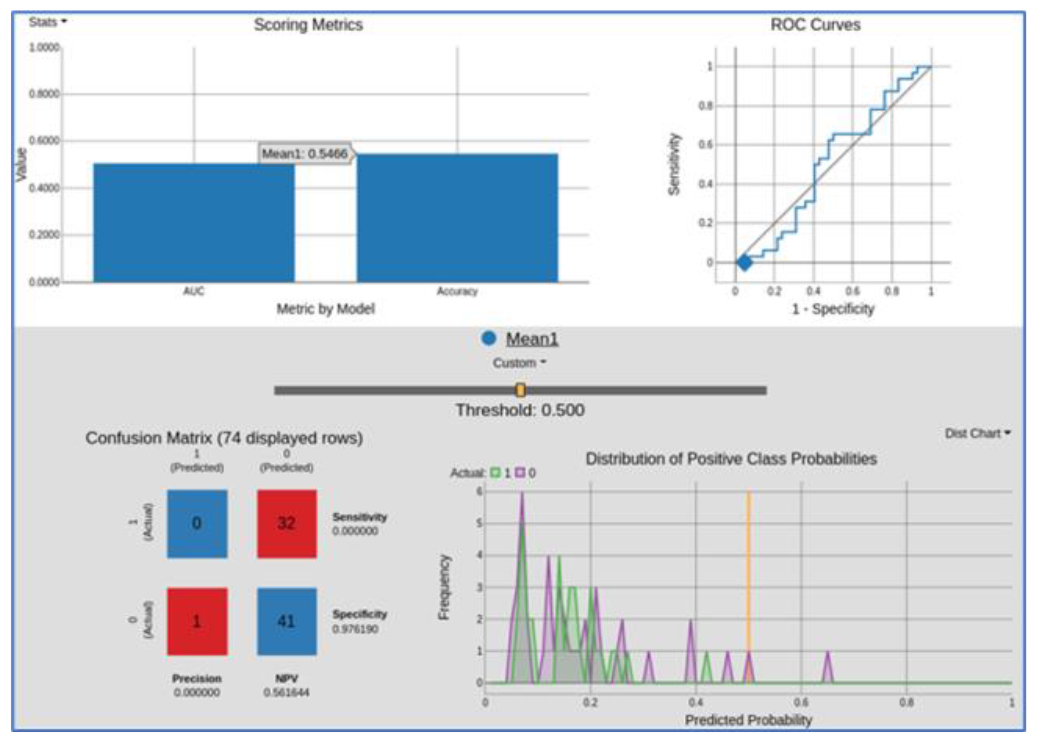
The figure was obtained from the Knime Binary Classification Inspector node. The view display performance of the RF-FPs, XGBoost-FPs and GCNN model ensemble on the test set. The view display performance of ML model predictions. On the top left the model’s statistic bar chart: AUC and overall accuracy. On the top right the ROC curve. The confusion matrix on the bottom left and the distribution of positive class probabilities on the bottom right. The statistical metrics adapt according to the threshold value.

**Figure 14.**
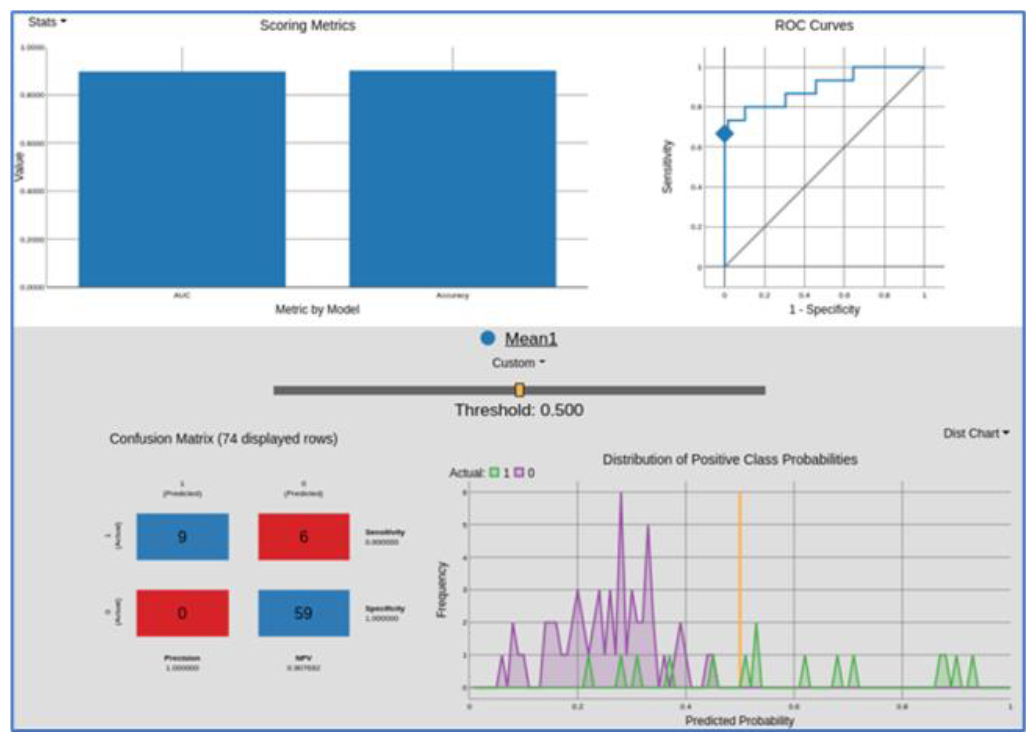
The view display performance of the RF-E3FP, XGBoost-E3FP model ensemble on the validation set. The view display performance of ML model predictions. On the top left the model’s statistic bar chart: AUC and overall accuracy. On the top right the ROC curve. The confusion matrix on the bottom left and the distribution of positive class probabilities on the bottom right. The statistical metrics adapt according to the threshold value.

**Figure 15.**
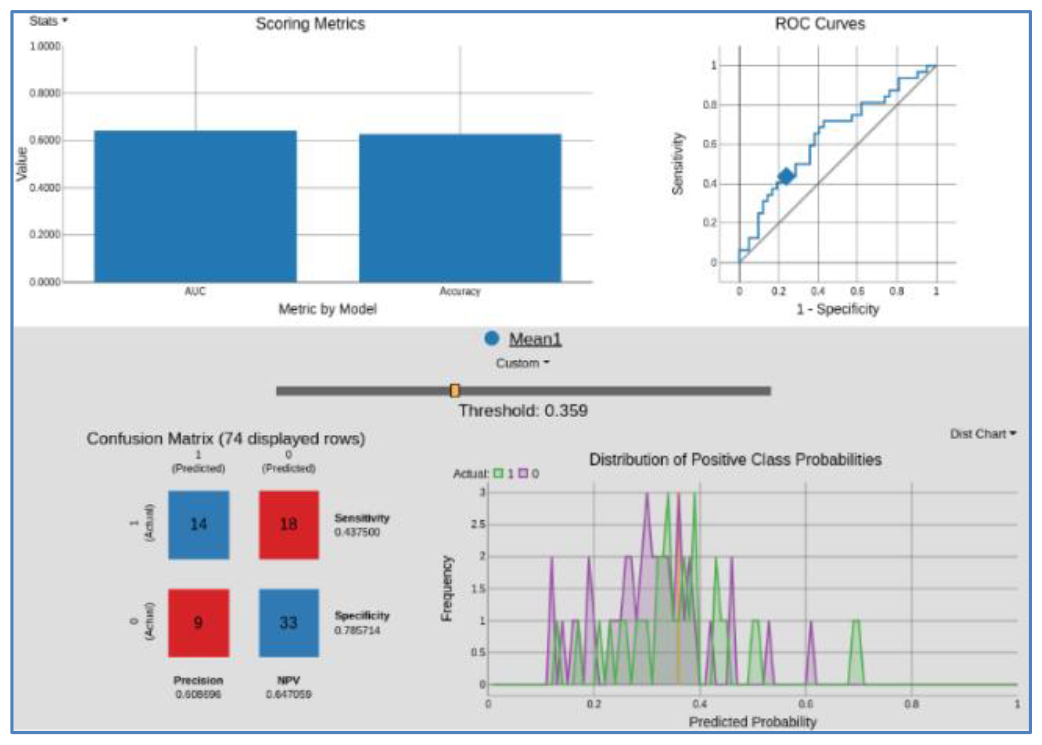
The view display performance of the RF-E3FP, XGBoost-E3FP model ensemble on the test set. The view display performance of ML model predictions. On the top left the model’s statistic bar chart: AUC and overall accuracy. On the top right the ROC curve. The confusion matrix on the bottom left and the distribution of positive class probabilities on the bottom right. The statistical metrics adapt according to the threshold value.

In the order to identify better ML models, the Extended 3-Dimensional FingerPrint^[31]^ (E3FP default parameters) was evaluated on the previously assembled ML dataset. A similarity sorting split 80/10/10 was performed with an ensemble of models (XGBoost and RF).

#### ML performance on the validation set

ROC AUC: 0.898

Kappa: 0.761

#### ML performance with the test set

ROC AUC: 0.642

Kappa: 0.231 (adjusted threshold)

The model ensemble XGBoost-E3FP and RF-E3FP shows better predictive performance compared to the previous ML model ensemble (RF-FPs, XGBoost-FPs and GCNN). While the performance, still weak, would not speak in favor of using this model in RL, it has been considered that a post-filtering of DGM generated molecules could be useful with a class probability threshold < 0.359 as it would help to exclude compounds with a higher probability of being inactive.

### Reinforcement Learning

Four scoring components were employed to direct the de novo design among which two were readily available in Reinvent 2.0 and two were, as described above, developed for this study. The quantitative estimate of drug-likeness QED scoring component was used to help the DGM generating “drug-like” molecules. The Jaccard distance component was employed to assist the model in generating molecule dissimilar to already known Mpro inhibitors. The 3D pharmacophore/ shape alignment component and SMC scoring component were applied to help the DGM to generate Mpro bioactive molecules.

Two DGM scenarios were considered for generating molecules, for the first scenario, “exploration”, the pre-trained DGM was used as such to explore the chemical space for the generation of new Mpro inhibitors. For the second scenario, “exploitation”, the DGM was retrained with a set of known Mpro inhibitors collected from the Open Science consortium COVID Moonshot and the ChEMBL database to exploit the known Mpro chemical space.

The two scenarios were then used in RL fashion employing the scoring components described above. Both systems were trained for 500 epochs for the exploitation mode and 1000 epochs for the exploration mode. In both cases, until the scores of each individual scores plateaued out. The two trained DGMs were utilized to generate molecules that were afterwards further filtered. Filters such as molecule sanitizer, QED, SMC, Jaccard distance, SMARTS, PheSA, ML affinity, AiZynthfinder, molecular diversity, molecular docking, and a final consensus score were employed to enrich the dataset with molecules virtually possessing the desired properties. The SMARTS filters used in the filtering workflow were assembled in a previous work.^[32]^

During the exploitation approach, the originally trained prior with ChEMBL molecules was retrained for 40 epochs with the DGM-TL dataset containing 338 known Mpro inhibitors. The purpose of this “Transfer Learning” approach is to bias the original DGM with compounds from the Mpro chemical space with the expectation that the generated molecules will be similar to the ones it was trained for.

Once the prior was retrained, the system was used in a RL context for 500 epochs where the PheSA, Jaccard distance, SMC, QED scoring components were utilized to score the generated molecules and assist the system in understanding the desired properties.

The best RL agent was obtained with the following parameters:

- Epochs: 500
- Scoring components:
- QED weight: 1.0
- PheSA weight: 4.0
- SMC weight: 1.0
- Jaccard distance (to ref. Cpds) weight: 1.0

The individual scores of the exploitation mode were monitored with Tensorboard as depicted in Fig. 16. The graphs show a score increase across all scoring components. However, a plateau is reached for PheSA after 350 epochs and for QED, while unstable, after 150 epochs.

**Figure 16.**
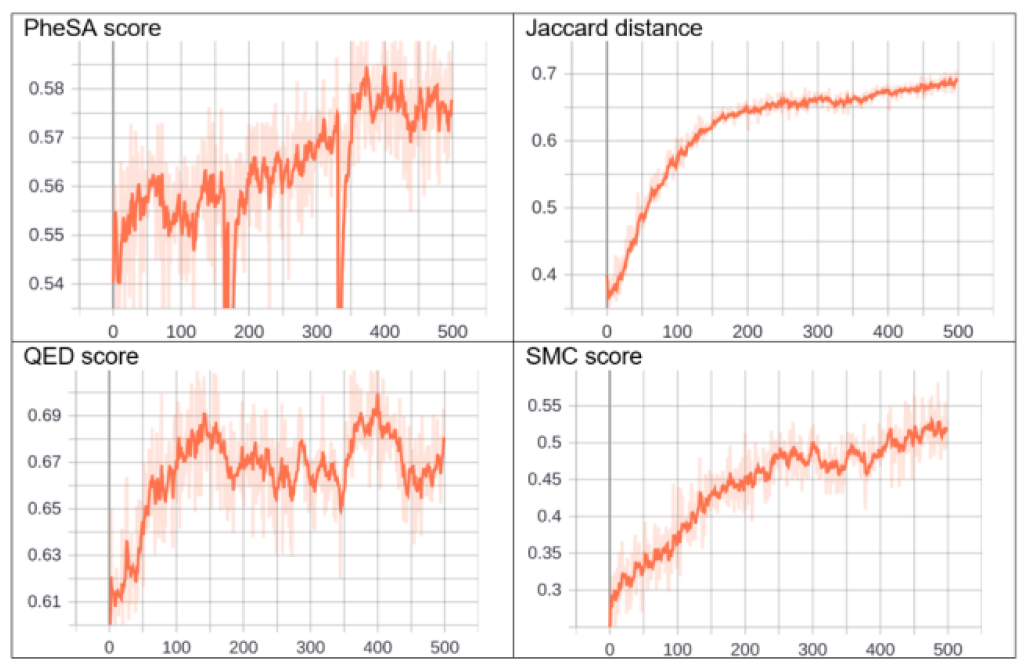
Tensorboard shows how the scores change over every epoch for the exploitation mode

During the exploration apprach, we utilized the original pre-trained prior based on ChEMBL molecules without any retraining. This model was directly applied in a Reinforcement Learning (RL) setting, incorporating scoring components like PheSA, Jaccard distance, SMC, and QED. We ran this setup for 1000 epochs to help the system grasp the targeted molecular properties. The aim of this second approach was to ascertain if the system could produce the desired molecules with reduced bias.

The individual scores of the exploration mode were monitored with Tensorboard as depicted in Fig. 17. The plots show an increase in learning for PheSA and SMC scores. A plateau is reached after 600 epochs for PheSA and 400 epochs for SMC. For the Jaccard distance, a steep increase was observed then a decline and again an increase. For QED, the score improved until 400 epochs and then deteriorate.

**Figure 17.**
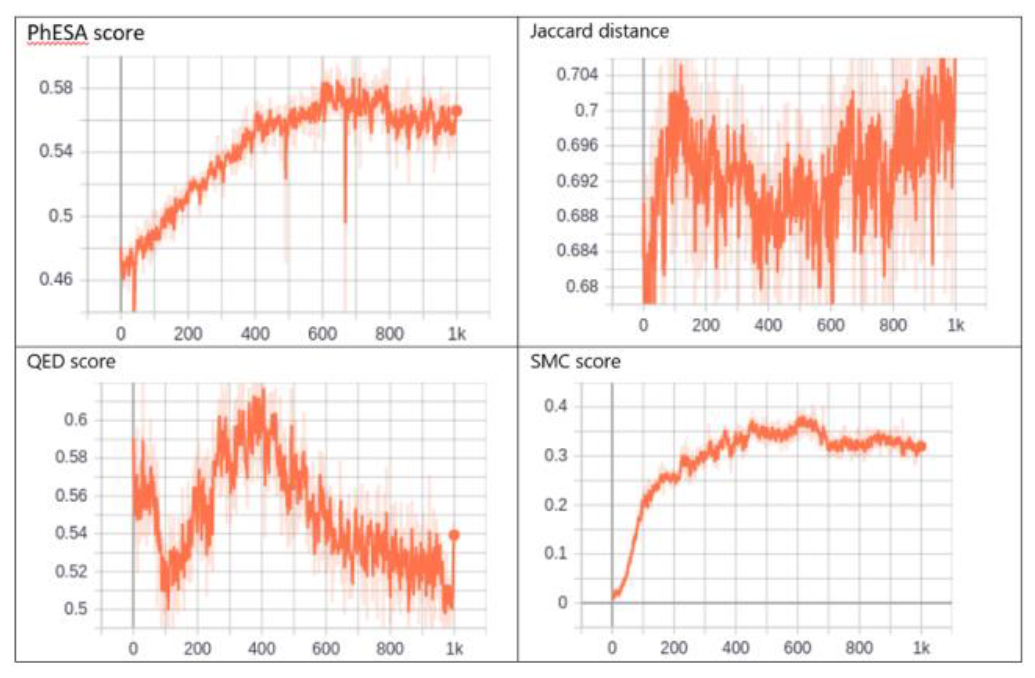
Tensorboard shows how the scores change over every epoch for the exploration mode

By comparing the 2 approaches, focused and exploration, it is noticeable that the focused approach allowed an overall better and quicker learning process. It is also interesting to see that for PheSA and Jaccard distance similar score can be obtained for both approaches. For QED, it was difficult to improve or even maintain the original scores while using the exploration approach. Finally, the transfer learning significantly helped the system to generate structure containing privileged fragments as we can see by comparing both SMC score figures.

The above trained agents were used to generate 400,000 molecules. The generated molecules were filtered with the following filter cascade:

- Duplicate removal
- Valid smiles only
- SMARTS filters
- QED > 0.5
- SMC > 0.5
- Distance > 0.5
- PheSA > 0.63
- Synthesizabilty > 0.85 (AIzynthfinder with Enamine and MolPort building blocks)
- ML classification > 0.3 (Class 1 probability)

1,736 molecules survived all filters. 1,007 molecules from the exploration approach and 729 from the exploitation approach.

### Docking with SeeSAR

Three crystal structures corresponding to three non-covalent chemical series were selected:

- MPro-x11294 (quinolone)
- MPro-x12692 (3-amino-pyridine)
- MPro-P0009 (Ugi-non-covalent)

The docking validation dataset was docked into MPro-x12692 and MPro-P0009. Two pharmacophore constraints (acceptor spheres) were added to guide the docking for two key H-bond interactions with the conserved H163 and E166 residues.

A template docking (10 poses) using the co-crystallized ligand was performed. After docking, a visual inspection allowed us to filter out poses for which one or two of the key interactions were missing.

The validation set was then also docked against MPro-x11294. Two pharmacophore constraints (acceptor spheres) were added to guide the docking for two key H-bond interactions with the H163 and either N142 and/or G143 residues (as observed from the crystal structure)

A template docking (10 poses) using the co-crystallized ligand was performed. After docking, a visual inspection allowed us to filter out poses for which one or two of the key interactions were missing.

The compounds with acceptable poses were predicted to be active, and the other compounds (either non-acceptable binding poses or when SeeSAR could not generate poses) were predicted as negative.

The evaluation of the predictive power led to the following results:

- Precision: 0.64 (for 22 predicted active 14 were actually active)
- Kappa: 0.24
- ROC AUC: 0.62

As we can see from the results, a docking approach does not represent an ideal classifier nevertheless, it allows, among other filter (PheSA, ML, SMC) to enrich the final dataset with possible new inhibitors.

Filtering: The combined 1736 compounds were docked following the above description (validation). Approximately 188 compounds were selected according to their binding poses. A hard cut-off was applied on the SeeSAR estimated affinity and only compounds below 3500uM were retained. 118 were then selected after docking.

### Docking with GOLD

To get the input molecules ready for docking, their three-dimensional (3D) structure needs to be determined. This step was accomplished using the oechem.package^[33]^. Similarly, the proteins were prepared with the same package. This process involved removing any atoms not part of the protein, reconstructing missing side chains and loops, and adding hydrogen atoms to the protein structure. The ligands, which are smaller molecules that bind to proteins, were retrieved from Protein Data Bank (PDB) files using the biopython package. These ligands were then processed using the rdkit module, a process that involved removing any salts and neutralizing charges in the ligands.

The idea behind the following workflow is that compounds with similar 3D structures would perturb the binding pocket in the same way. To identify a reference compound that closely resembles the ligands, the PheSA program was used. PheSA was employed to calculate the similarity between all the ligands to evaluate and the active conformations of reference compounds.

For the docking process, hydrogen atoms were also added to the input molecules. The docking itself was conducted using the GOLD docking program, which allows for automatic non-covalent docking. This procedure was integrated into a script for efficiency. The docking utilized both the prepared PDB files of the ligands most similar to the reference compounds and the prepared input molecules.

### Consensus scoring (CS)

It has been reported that CS, which combines multiple scoring functions in binding affinity estimation, leads to higher hit rates in virtual library screening studies.^[34]^ It has also been demonstrated that consensus scoring outperforms, for statistical reason, any single scoring.^[35]^

A consensus scoring was developed taking PheSA, docking SeeSAR & Gold, SMC and binding free energy score into account.

- The SMC and PheSA scores were already normalised from 0 to 1 so the scores were kept unchanged
- ChemScore was normalized from 0 to 1
- SeeSAR Hyde_IC50 was converted to pIC50 = - log(Hyde_IC50) then the score was normalized from 0 to 1
- The CS was calculated with the following equation:

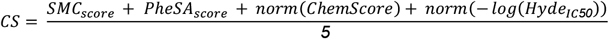

The CS score was used to make the final selection of compounds to be purchased.

### Hit expansion & docking with Glide

Analogues, based on 2D similarity, were identified in either the Chemspace^[22]^ screening compound stock or in the Idorsia corporate collection. The analogues were docked with H-bond constraints using grids generated from publicly available crystal structures.

The hit expansion (2D similarity analogue search) was performed for the 3 hit series (piperazine, cyclized urea and N-benzoimidazol-1-yl-acetamide). The hit expansion was done by searching the Idorsia corporate collection for the piperazine (cluster 1) and the N-benzoimidazol-1-yl-acetamide compound (cluster 3). For the cyclized ureas (cluster 2) a hit expansion was performed in the Chemspace in stock screening compounds as no analogues could be identified from the corporate collection.

- 1400 analogues of cluster 1 (positive control included) were identified
- 4,402 analogues of cluster 2 were identified
- 128 analogues of cluster 3 were identified

Positive control: 3 reference MLDD compounds + 2 Moonshot active compounds synthesized internally.

For cluster 1, the 1400 analogues were docked with Schrodinger Glide.

The fragalysis complex Mpro-x11812 (co-crystalized ligand) was selected for its similarity with the cluster 1 hits and used to generate a docking grid.

The analogues were docked with grid-based with the following constraints (at least two constraints):

- H-bond with H163
- H-bond with G143
- H-bond with C145
- The docking poses were inspected, the best poses were starred, and 115 compounds were selected.

The positive control molecules were present in the top docking score list and provided some in-silico evidence on the validity of the approach.

110 compounds were selected from the Idorsia corporate collection and submitted to the biological measurements.

For cluster 2, 4402 Chemspace analogues were docked with Schrodinger Glide.

The fragalysis complex Mpro-x11513 (co-crystalized ligand) was used for its similarity with the cluster 2 hits.

The docking (SP followed by XP) was performed on the analogues with grid-based constraints:

- H-bond with H163
- H-bond with E166

The docking poses were inspected, and the best poses were starred, and finally 43 compounds were selected, and a quote was requested at ChemSpace.

38 compounds were delivered and submitted to biological measurements.

For cluster 3, the 128 compounds were docked with Schrodinger Glide.

The fragalysis complex Mpro-x11612 (co-crystalized ligand) was used for its relative similarity with the cluster 3 hit.

The analogues were docked (SP followed by XP) with grid-based constraints:

- H-bond with H163
- H-bond with E166

The docking poses were inspected, and the best poses were starred, and finally 25 compounds were selected, and ordered from the Idorsia store for a biological assessment.

After visual inspection of the docking poses, the best compounds were selected, ordered, and measured in a FRET assay.

### Synthetic details for the preparation of compounds

Commercially available starting materials were used as received without further purification. Flash column chromatography was performed using Biotage SNAP cartridges (10−340 g) and elution was with a Biotage Isolera system. Merck pre-coated thin layer chromatography (TLC) plates were used for TLC analysis. Final compounds were purified to >95% purity (UV and NMR) by reverse phase preparative HPLC using a Waters XBridge column (10 μm, 75 × 30 mm). Conditions: MeCN [eluent A]; water + 0.5% NH4OH (25% aq.) [eluent B]; Gradient: 90% B → 5% B over 6.5 min (flow: 75 mL/min). Detection: UV/Vis + MS. Racemates can be separated into their enantiomers by chiral HPLC using a ChiralPaK IC column (5 μm, 250 × 4.6 mm). Conditions: Heptane + 0.05% DEA [eluent A]; EtOH + 0.05% DEA [eluent B]; Isocratic elution with 50% eluent B (flow: 1 mL/min), or a CHIRALCEL OZ-H column (5 mm, 250 × 4.6 mm). Conditions: CO2 [eluent A]; EtOH + 0.1% DEA [eluent B]; Isocratic elution with 50% eluent B (flow: 4 mL/min).

Mass spectrometry data were recorded by one of the following methods:

#### LC-MS with acidic conditions

##### Method A

Agilent 1100 series with mass spectrometry detection (MS: Finnigan single quadrupole). Column: Zorbax SB-aq (3.5 μm, 4.6 × 50 mm). Conditions: MeCN [eluent A]; water + 0.04% TFA [eluent B]. Gradient: 95% B → 5% B over 1.5 min (flow: 4.5 mL/min). Detection: UV/Vis + MS

##### Method B

Agilent 1100 series with mass spectrometry detection (MS: Finnigan single quadrupole). Column: Waters XBridge C18 (2.5 μm, 4.6 × 30 mm). Conditions: MeCN [eluent A]; water + 0.04% TFA [eluent B]. Gradient: 95% B → 5% B over 1.5 min (flow: 4.5 mL/min). Detection: UV/Vis + MS.

#### LC-MS with basic conditions

##### Method C

Agilent 1100 series with mass spectrometry detection (MS: Finnigan single quadrupole). Column: Zorbax Extend C18 (5 μm, 4.6 × 50 mm). Conditions: MeCN [eluent A]; 13 mmol/L NH3 in water [eluent B]. Gradient: 95% B → 5% B over 1.5 min (flow: 4.5 mL/min). Detection: UV/Vis + MS.

##### Method D

Agilent 1100 series with mass spectrometry detection (MS: Finnigan single quadrupole). Column: Waters XBridge C18 (5 μm, 4.6 × 50 mm). Conditions: MeCN [eluent A]; 13 mmol/L NH3 in water [eluent B]. Gradient: 95% B → 5% B over 1.5 min (flow: 4.5 mL/min). Detection: UV/Vis + MS.

#### LC−HRMS parameters were the following

analytical pump Waters Acquity binary, Solvent Manager, MS, SYNAPT G2 MS, source temperature of 150 °C, desolvation temperature of 400 °C, desolvation gas flow of 400 L/h; cone gas flow of 10 L/h, extraction cone of 4 RF; lens 0.1 V; sampling cone 30; capillary 1.5 kV; high resolution mode; gain of 1.0, MS function of 0.2 s per scan, 120−1000 amu in full scan, centroid mode. Lock spray: keucine enkephalin, 2 ng/mL (556.2771 Da), scan time of 0.2 s with interval of 10 s and average of 5 scans; DAD: Acquity UPLC PDA detector. Column was an Acquity UPLC BEH C18 1.7 μm, 2.1 mm × 50 mm from Waters, thermostated in the Acquity UPLC column manager at 60 °C. Eluents were the following: water + 0.05% formic acid; B, acetonitrile + 0.05% formic acid. Gradient was 2−98% B over 3.0 min. Flow was 0.6 mL/min. Detection was at UV 214 nm.

1 H NMR spectra were recorded on a Bruker (400 or 500 MHz) spectrometer in the indicated deuterated solvent. Chemical shifts are reported in ppm relative to solvent peaks as the internal reference.

**Compound 1; (4-(2,4-dichlorobenzyl)piperazin-1-yl)(6-fluoroisoquinolin-4-yl)methanone**

Commercially available material. LC-HRMS: t_R_ = 0.875 min; [M+H]/z = 418.0889 found = 418.2800.

**Compound 2; (4-(3,5-dichloro-2-(difluoromethoxy)benzyl) piperazin-1-yl)(pyridin-3-yl) methanone**

Commercially available material. LC-HRMS: t_R_ = 0.916 min; [M+H]/z = 416.0744 found = 416.2700.

**Compound 3; (4-(2,3-dichlorobenzyl)piperazin-1-yl)(pyridin-3-yl)methanone**

Commercially available material. LC-HRMS: t_R_ = 0.631 min; [M+H]/z = 350.0827 found = 350.2200.

**Compound 4; 1-(2-chlorobenzyl)-3-(6-chloroisoquinolin-4-yl)imidazolidin-2-one**

Commercially available material. LC-HRMS: t_R_ = 1.102 min; [M+H]/z = 372.0670 found = 372.2600.

**Compound 5; 1-(2,4-dichlorobenzyl)-3-(isoquinolin-4-yl)imidazolidin-2-one**

Commercially available material. LC-HRMS: t_R_ = 1.037 min; [M+H]/z = 372.0670 found = 372.2300.

**Compound 6; rac-(R)-5,6-dichloro-N-(4,6-difluoro-1H-benzo[d]imidazol-1-yl)-2,3-dihydrobenzofuran-3-carboxamide**

Commercially available material. LC-HRMS: Not available

**Compound 13; rac-(R)-4,5-dichloro-N-(1-(cyclopropylmethyl)-1H-imidazol-5-yl)-2,3-dihydrobenzofuran-3-carboxamide**

Commercially available material. LC-HRMS: t_R_ = 0.616 min; [M+H]/z = 352.0620 found = 352.2000.

**Compound 14; rac-(R)-N-(1-(3-chloro-4-fluorophenyl)-3-hydroxypropyl)-N-methyl-2-(2,4,5-trifluorophenyl)acetamide**

Commercially available material. LC-HRMS: t_R_ = 1.104 min; [M+H]/z = 390.0884 found = 390.1900.

**Compound 15; N-(4,5-dichloro-2-(1H-1,2,4-triazol-1-yl)benzyl)-2-(3,5-difluorophenyl)acetamide**

Commercially available material. LC-HRMS: t_R_ = 1.043 min; [M+H]/z = 397.0434 found = 397.1400.

**Compound 16; 1-(3,4-dichlorophenyl)-N-(4,5-difluoro-2-(pyridin-4-yl)benzyl)cyclopropane-1-carboxamide**

Commercially available material. LC-HRMS: t_R_ = 1.126 min; [M+H]/z = 433.0686 found = 433.2900.

**Compound 17; (4-(3,5-dichloro-2-fluorobenzyl)piperazin-1-yl)(furan-3-yl)methanone**

Commercially available material. LC-HRMS: t_R_ = 0.854 min; [M+H]/z = 357.0573 found = 357.2000.

**Compound 18; 1-(4-(3,4-dichlorobenzyl)piperazin-1-yl)-2-(1H-1,2,4-triazol-1-yl)ethan-1-one**

Commercially available material. LC-HRMS: t_R_ = 0.526 min; [M+H]/z = 354.0888 found = 354.2300.

**Compound 19; N-(3,4-dichlorobenzyl)-2-(1H-tetrazol-1-yl) acetamide**

Commercially available material. LC-HRMS: t_R_ = 0.854 min; [M+H]/z =286.0262 found = 286.1700.

**Compound 20; 2-(4-cyclopropyl-1H-1,2,3-triazol-1-yl)-N-(3,4-dichlorobenzyl)acetamide**

Commercially available material. LC-HRMS: t_R_ = 0.966 min; [M+H]/z =325.0623 found = 325.2100.

**Compound 21; 6,8-dichloro-N-(1-ethyl-1H-imidazol-5-yl)-3-methylchromane-4-carboxamide**

Commercially available material. LC-HRMS: t_R_ = 0.675 min; [M+H]/z =354.0776 found = 354.4000.

**Compound 22; rac-(R)-4-chloro-N-(2H-pyrazolo[3,4-c]pyridin-4-yl)bicyclo[4.2.0]octa-1(6),2,4-triene-7-carboxamide** Commercially available material. LC-HRMS: t_R_ = 0.656 min; [M+H]/z =299.0700 found = 299.2500.

**Compound 7; (S)-1-(2-(3-chlorophenyl)pyrrolidin-1-yl)-2-(5-methylpyridin-3-yl)ethan-1-one**

DIPEA (0.0524 mL, 0.3 mmol, 3 eq) was added to a solution of 2-(5-methylpyridin-3-yl)acetic acid hydrochloride (0.1 mmol, 1 eq) in DMF p.a (0.20 mL), followed by a solution of (2S)-2-(3-Chlorophenyl)pyrrolidine (20.7 mg, 0.105 mmol, 1.05 eq) and DIPEA (0.0349 mL, 0.2 mmol, 2 eq) in DMF p.a (0.4 mL). Finally, a 0.5 M HATU stock solution was added to DMF (220 µL, 0.11 mmol, 1.1 eq) and stirred overnight at RT. The product was isolated by basic preparative HPLC (29 mg, 90% yield). LC-HRMS: t_R_ = 0.633 min; [M+H]/z = 315.1264 found = 315.4000; 1H NMR: Presence of 2 stable conformational isomers in DMSO at RT, ratio 2:1, δH(500 MHz, DMSO): 8.28 (0.67H, dd, J=2.0, 0.9 Hz), 8.24 (0.67H, d, J=2.1 Hz), 8.19 (0.33H, dd, J=2.1, 0.9 Hz), 7.95 (0.33H, d, J=2.0 Hz), 7.45 – 7.18 (3.34H, m), 7.17 – 7.08 (1.66H, m), 5.28 (0.33H, dd, J=8.1, 2.3 Hz), 5.02 (0.67H, dd, J=8.1, 3.1 Hz), 3.88 – 3.77 (1.33H, m), 3.74 – 3.62 (1.67H, m), 3.59 – 3.48 (0.67H, m), 3.18 – 3.14 (0.33H, m), 2.42 – 2.36 (0.33H, m), 2.28 (2H, d, J=0.7 Hz), 2.26 – 2.23 (0.33H, m), 2.20 (1H, d, J=0.8 Hz), 1.97 – 1.67 (3.34H, m).

**Compound 8; 2-((4-(5-chloronicotinoyl)-1,4-diazepan-1-yl)methyl)benzonitrile**

### Step 1: tert-butyl 4-(5-chloronicotinoyl)-1,4-diazepane-1-carboxylate

N-(t-Butyloxycarbonyl)-homopiperazine hydrochloride (947 mg, 4 mmol, 1 eq), 5-Chloronicotinic acid (715 mg, 4.4 mmol, eq) and DIPEA (2.05 mL, 12 mmol, 3 eq) were dissolved in DCM (40 mL). HATU (1725 mg, 4.4 mmol, 1.1 eq) was added and the mixture was stirred at RT for 30 min. Aq. NaHCO3 was added, and the product was extracted 2x with DCM. The combined organic phases were dried over MgSO4 and concentrated under reduced pressure. The product was isolated by flash chromatography (1.565 g, 115% yield). LC-MS D: t_R_ = 0.82 min; [M+H]+ = 340.17.

### Step 2: (5-chloropyridin-3-yl)(1,4-diazepan-1-yl)methanone hydrochloride

tert-butyl 4-(5-chloronicotinoyl)-1,4-diazepane-1-carboxylate (1565 mg, 3.9 mmol, 1 eq) was dissolved in MeOH (30 mL) and HCl 4M in dioxane (5 mL, 20 mmol, 5.134 eq) was added. The mixture was stirred at RT for 4h. Some extra HCl (2 eq.) was added, and the mixture was stirred at RT for 1h to reach full completion of the reaction. The solvents were evaporated under reduced pressure. A white solid was obtained. LC-MS D: t_R_ = 0.48 min; [M+H]+ = 240.13.

### Step 3: 2-((4-(5-chloronicotinoyl)-1,4-diazepan-1-yl)methyl) benzonitrile

(5-chloropyridin-3-yl)(1,4-diazepan-1-yl)methanone hydrochloride (102 mg, 0.3 mmol, 1 eq), 2-Cyanobenzaldehyde (48.2 mg, 0.36 mmol, 1.2 eq) and DIPEA (0.154 mL, 0.9 mmol, 3 eq) were dissolved in DCM (3 mL). Sodium triacetoxyborohydride 97% (167 mg, 0.75 mmol, 2.5 eq) was added and the mixture was stirred overnight at RT. Saturated aqueous NaHCO3 (0.5 mL) was added, the organic phase was filtered through a phase separator and evaporated in a Genevac overnight. The product was isolated by RP prep HPLC, basic condition, polar gradient. LC-HRMS: t_R_ = 0.527 min; [M+H]/z = 355.1264 found = 355.4000.

**Compound 9; (S)-5-(5-chloro-2-fluorobenzyl)-3-(5-fluoro pyridin-3-yl)imidazolidine-2,4-dione**

Commercially available material. LC-HRMS: t_R_ = 0.926 min; [M+H]/z = 338.0508 found = 338.4000.

**Compound 10; 1-(5,7-dichloro-3,4-dihydroisoquinolin-2(1H)-yl)-2-(pyridin-3-yl)ethan-1-one**

Commercially available material. LC-HRMS: t_R_ = 0.746 min; [M+H]/z = 321.0561 found = 321.3000.

**Compound 11; (4-(2-chlorobenzyl)-1,4-diazepan-1-yl)(5-chloropyridin-3-yl)methanone**

### Step 1: tert-butyl 4-(5-chloronicotinoyl)-1,4-diazepane-1-carboxylate

N-(t-Butyloxycarbonyl)-homopiperazine hydrochloride (947 mg, 4 mmol, 1 eq), 5-Chloronicotinic acid (715 mg, 4.4 mmol, eq) and DIPEA (2.05 mL, 12 mmol, 3 eq) were dissolved in DCM (40 mL). HATU (1725 mg, 4.4 mmol, 1.1 eq) was added and the mixture was stirred at RT for 30 min. Aq. NaHCO3 was added, and the product was extracted 2x with DCM. The combined organic phases were dried over MgSO4 and concentrated under reduced pressure. The product was isolated by flash chromatography (1.565 g, 115% yield). LC-MS D: t_R_ = 0.82 min; [M+H]+ = 340.17.

### Step 2: (5-chloropyridin-3-yl)(1,4-diazepan-1-yl)methanone hydrochloride

tert-butyl 4-(5-chloronicotinoyl)-1,4-diazepane-1-carboxylate (1565 mg, 3.9 mmol, 1 eq) was dissolved in MeOH (30 mL) and HCl 4M in dioxane (5 mL, 20 mmol, 5.134 eq) was added. The mixture was stirred at RT for 4h. Some extra HCl (2 eq.) was added, and the mixture was stirred at RT for 1h to reach full completion of the reaction. The solvents were evaporated under reduced pressure. A white solid was obtained. LC-MS D: t_R_ = 0.48 min; [M+H]+ = 240.13.

### Step 3: (4-(2-chlorobenzyl)-1,4-diazepan-1-yl)(5-chloropyridin-3-yl)methanone

(5-chloropyridin-3-yl)(1,4-diazepan-1-yl)methanone (34.0 mg, 0.10 mmol) was dissolved in a mixture of DCM (9.5 mL) and DIPEA (0.50 mL). 2-Chlorobenzaldehyde (0.12 mmol, 1.2 eq) was added, followed by Sodium triacetoxyborohydride 97% (55.8 mg, 0.25 mmol, 2.5 eq). The mixtures were stirred at RT overnight. Aqueous NaHCO3 and DCM were added, the products was extracted 3x with DCM. Finally, the product was isolated by reverse phase preparative HPLC, polar gradient. LC-HRMS: tR = 0.516 min; [M+H]/z = 364.0983 found = 364.2700 ; 1H NMR: δH(500 MHz, DMSO): 8.70 (1H, dd, J=7.0, 2.4 Hz), 8.57 (1H, dd, J=9.3, 1.8 Hz), 8.02 (1H, dt, J=18.1, 2.1 Hz), 7.57 – 7.39 (2H, m), 7.38 – 7.23 (2H, m), 3.73 (1H, s), 3.69 (1H, s), 3.69 – 3.64 (2H, m), 3.41 (2H, td, J=5.7, 2.8 Hz), 2.82 – 2.76 (1H, m), 2.74 – 2.66 (2H, m), 2.66 – 2.60 (1H, m), 1.89 – 1.81 (1H, m), 1.78 – 1.69 (1H, m).

**Compound 12; 1-(6,7-dichloro-3,4-dihydroisoquinolin-2(1H)-yl)-2-(5-methylpyridin-3-yl)ethan-1-one**

DIPEA (0.0524 mL, 0.3 mmol, 3 eq) was added to a solution of 6,7-dichloro-1,2,3,4-tetrahydroisoquinoline hydrochloride (0.1 mmol, 1 eq) in DMF p.a (0.20 mL), followed by a solution of 2-(5-methylpyridin-3-yl)acetic acid hydrochloride (20.7 mg, 0.105 mmol, 1.05 eq) and DIPEA (0.0349 mL, 0.2 mmol, 2 eq) in DMF p.a (0.4 mL). Finally, a 0.5 M HATU stock solution was added to DMF (220 µL, 0.11 mmol, 1.1 eq) and stirred overnight at RT. The product was isolated by basic preparative HPLC (30 mg, 90% yield). LC-HRMS: t_R_ = 0.738 min; [M+H]/z = 335.0718, found = 335.2100; 1H NMR: Presence of 2 stable conformational isomers in DMSO at RT, ratio 6:4, δH(500 MHz, DMSO): 8.30 – 8.22 (2H, m), 7.56 (0.6H, s), 7.50 (0.6H, s), 7.49 (0.4H, s), 7.48 (0.4H, s), 7.45 (0.6H, s), 7.42 (0.4H, s), 4.75 (0.8H, s), 4.62 (1.2H, s), 3.81 (1.2H, s), 3.80 (0.8H, s), 3.76 (1.2H, t, J=5.9 Hz), 3.67 (0.8H, t, J=6.0 Hz), 2.83 (1.2H, t, J=5.9 Hz), 2.77 (0.8H, t, J=6.0 Hz), 2.27 (1.8H, s), 2.25 (1.2H, s).

### Fluorescence resonance energy transfer (FRET)-based Mpro proteolytic activity assay

The enzymatic activity of the recombinant SARS-CoV-2 main protease Mpro was determined by a fluorescence resonance energy transfer (FRET) assay using a custom synthesized peptide substrate with (7-Methoxycoumarin-4-yl)acetyl [MCA] as fluorophore and 2,4-Dinitrophenyl [DNP] as fluorescence quencher: MCA-Ala-Val-Leu-Gln-Ser-Gly-Phe-Arg-Lys(Dnp)-Lsy-NH2-trifluoroacetate salt (Bachem AG, Bubendorf CH). This peptide substrate amino acid sequence corresponds to the nsp4/nsp5 (Mpro) cleavage site. A substrate stock solution (10 mM) was prepared in 100 % DMSO. 40 µL of a 4 µM substrate solution prepared in H2O/TWEEN-20 0.01%) is added to a solution (40 µL) containing Mpro to start the enzymatic reaction. The final concentrations of the assay reaction ingredients (80 µl) are 5 nM [E] Mpro, 2 µM [S] peptide substrate (Km 3.17 µM), 1 mM DTT, 1.2 % DMSO, 0.01 % TWEEN-20 25 mM TRIS pH 7.4. (?0.5 mM EDTA?). Mpro was diluted (10 nM) from aliquotes stored as stock solution (512 µM, -80 °C, storage buffer) in Mpro assay buffer (50 mM TRIS pH 7.4, 1 mM EDTA, 2 mM DTT, 0.01 % TWEEN-20). The rate of Mpro enzymatic activity (v) was determined by monitoring the increase in fluorescence intensity of reactions at room temperature in black microplates (NUNc 384-well F-bottom) with an Infinite M-1000 plate reader (Tecan) using 325 nm and 400 nm as wavelengths for excitation and emission, respectively. Test compounds were dissolved in DMSO and screened first at a 25 µM. 3-fold serial dilutions (125 µM – 6.35 nM) of small molecule test compounds are added to determine inhibitory potency. IC50 is determined by an in-house evaluation tool (IC50 Studio with 4-parametric fitting, Hill-equation).

Small molecule compounds showing putative inhibitory activity were tested in a separate assay for quenching potential of fluorescence emitted by the MCA fluorophore (Bioquest) to identify possible false positives.

### Fluorimetric human liver Cathepsin L (hCatL) activity assay

To determine the effect of small molecule test compounds on the enzymatic activity of human Cathepsin L a fluorescence-based assay has been implemented according to a published protocol.^[36]^

Cathepsin L from human liver and the fluorogenic peptidomimetic substrate Z-Phe-Arg7-amido-4-methylcoumarin hydrochloride (Z-FR-AMC) were purchased from SIGMA (#219402, #C9521). The Cathepsin L enzyme buffer consisted of 50 mM Tris pH 6.5, 5 mM EDTA, 200 mM NaCl, 2 mM DTT. The hCatL enzymatic reaction was initiated by adding 40 µL of a solution containing the substrate at 4 µM (50 mM Tris pH 6.5, 5 mM EDTA, 200 mM NaCl, 0.005 % triton-X-100) to 40 µL solution consisting of enzyme in assay buffer (50 mM Tris pH 6.5, 5 mM EDTA, 200 mM NaCl, 2 mM DTT, 1 nM Ca). Test compounds dissolved in DMSO were dispensed as 2-fold serial dilutions (125 µM – 244 nM) to black well assay plates (384 well, NUNC).

Final assay reaction mixtures consisted of Cathepsin L 0.5 nM [E], Z-Phe-Arg7-amido-4-methylcoumarin [S], [cpds] 125 – 0.214 µM or 50 – 0.0977 µM, 50 mM Tris-HCL pH 6.5, 200 mM NaCl, DTT 1 mM, 2.5 mM EDTA, Triton-X100 0.0005%, DMS0 1.27 %.

Fluorescence emitted by AMC fluorophore liberated by hCatL cleavage of the substrate was measured with a Tecan infinite M-1000 plate reader with filters for excitation at 360/40 nm and emission at 460/40 nm at RT immediately after initiating the reaction t0 and 35 min incubation at RT.

Leupeptin (SIGMA L5793) a validated natural protease inhibitor shows the expected biochemical potency in this assay with an IC50 value of around 2 nM.

### Cloning, protein expression and purification of SARS-CoV-2 Mpro

DNA encoding a recombinant fusion protein (supplementary information) composed of SUMO with a N-terminal hexa-histidine tag and Mpro (NC_045512.2, Nsp5, YP_009742612, Wuhan-Hu-1) was codon optimized for expression in E. coli and synthesized at Genscript. The synthetic DNA was cloned into pET29a(+)::[NdeI, BamHI] (Genscript) and transformed into BL21(DE3) cells. The fusion protein was expressed overnight (Luria broth medium, 25 µg/ml Kanamycin) at 18 ºC after induction with 0.5 mM isopropyl-b-d-thiogalactoside IPTG at OD600∼0.7. Overnight cultures were spun down and recovered cell paste was stored at -70 ºC. 12 g cell paste was resuspended in buffer (20 mM Tris-HCl, pH 7.8; 150 mM NaCl, 5 mM imidazole) and treated with lysozyme (0.1mg/ml; 30 min) and benzonase (2500 Units, 10 mM MgCl2; 15 min, RT). Bacterial cells were lysed by high pressure homogenization (29008 psi, Microfluidics MP110P, DIXC H10Z) and centrifuged 30 min at 16000 rpm (Fiberlite F21-8×50y). The hexa-histidine SUMO-Mpro fusion protein was purified by immobilized metal affinity chromatography (IMAC) with a HisTrap column (5 ml, Cytiva) connected to a FPLC Äkta purifier 100 system. His-tagged SUMO_Mpro fusion protein was eluted with a linear gradient of increasing imidazole concentration (elution buffer, 0-100 %, 20 column volumes, 20 mM Tris-HCl pH 7.8, 150 mM NaCl, 500 mM imidazole). Eluate fractions containing the target protein were combined and concentrated (Amicon, 10 kDa). The fusion protein was treated with SUMO protease (Sigma SAE0067, 5 U/ mg target protein) to liberate Mpro with authentic N-(Ser1) and C-termini (Q306)-C. The mixture of SUMO protease cleavage products was dialyzed overnight (4 ºC, 4 L, 20 mM Tris-HCl, 150 mM NaCl, Slide-A-Lyzer→ cassette, 10 kDa, Thermo Scientific). The hexa-histidine tagged SUMO protein was separated from non-tagged authentic Mpro present in the dialysate by IMAC and collecting Mpro in the flow through. Mpro was further purified by size exclusion chromatography (SEC, Hiload 26/60 Superdex 200) with storage buffer (20 mM Tris-HCl, 150 mM NaCl, 1 mM TCEP, 1 mM EDTA). The SEC elution volume for Mpro indicated a dimer as the oligomeric state. Pure (97 %, LC MS) Mpro was concentrated (Amicon, 10 kDa) to a concentration of 17 mg/ml (512 µM) Mpro and stored at -70 ºC.

### Surface plasmon resonance (SPR) binding analysis

The SPR experiments were performed using a Biacore T200 equipped with a Series S Sensor Chip SA (GE Healthcare #BR-1005-31). Biotinylated MproQ306E.AVI (35752 Da, >85% pure, only 21% biotinylated based on MS) was immobilized to the streptavidin covalently attached to a carboxymethyldextran matrix. The initial conditioning of the surfaces on flow cell 1 and 2 was performed by three 1-minute pulses of 1 M NaCl, 50 mM NaOH solution. The ligand at a concentration of 0.27mg/ml in immobilization buffer (10 mM HEPES, 150 mM NaCl, 1mM TCEP, 0.05% P20, pH 7.4) was immobilized at a density of 4300 RU on flow cell 2 at a flow rate of 5 μl/min and flow cell 1 was left blank to serve as a reference surface. Surfaces were stabilized with 3 hours injection at a flow rate of 40 µL/min of running buffer (10 mM HEPES, 150 mM NaCl, 1mM TCEP, 0.05% P20, 5% DMSO, pH 7.4).

To collect kinetic binding data, sample in 10 mM HEPES, 150 mM NaCl, 0.05% P20, 5%DMSO, pH 7.4, was injected over the two flow cells at ascending concentrations (0, 0.27, 0.82, 2.5, 7.5, 22.2, 66.6 and 200 µM) at a flow rate of 40 μl/min and at a temperature of 25°C. The complex was allowed to associate and dissociate for 40 and 100 s, respectively for each sample concentration.

A DMSO correction curve was performed before/after every 104 cycles.

Data were collected at a rate of 10 Hz and fitted to a simple 1:1 interaction model using the global data analysis option available within Biacore T200 Evaluation software.

### Crystallization and crystallography

Sitting drop co-crystallization was carried out by adding 0.3 µl each of Mpro (17 mg/ml, buffer as described above) and crystallization solution in 96-well plates (Intelliplate, Art Robbins). The inhibitors were added to the protein approximately one hour before crystallization. IDOR-1142-4622 and IDOR-1142-4624 were added to Mpro at a final concentration of 1 mM from a 10 mM stock solution in DMSO, while IDOR-1142-0506 was added at a final concentration of 10 mM from a 100 mM stock solution in DMSO.

The inhibitors IDOR-1142-0506 and IDOR-1142-4624 were co-crystallized using 10% (w/v) polyethylene glycol (PEG) 4000, 20% (w/v) glycerol, 0.03 M each of sodium fluoride, sodium bromide and sodium iodide, 0.1 M MES/imidazole pH 6.5 (condition B3 of “Morpheus I” screen, Molecular Dimensions Ltd.). IDOR-1142-4622 was co-crystallized using 0.1 M MES/imidazole pH 6.5, 10% (w/v) polyethylene glycol 4000, 20% (w/v) glycerol, 0.03 M each of magnesium chloride and calcium chloride (condition A3 of Morpheus I screen, Molecular Dimensions Ltd).

X-ray diffraction experiments were carried out at 100 K at beamline X06SA-PXI of the Swiss Light Source (SLS), Villigen, Switzerland. Measurements were made at the SLS with crystal rotation steps of 0.2 º using an EIGER 16M detector (Dectris). The data were processed and scaled using AutoProc^[37]^ and XSCALE^[38]^, respectively. Automated molecular replacement was carried out using Dimple^[39]^ (Collaborative Computational Project 1994) with the 3CLpro structure as template. COOT^[40]^ was used for model building. Phenix.refine^[41]^, Buster^[41]^ and Refmac^[42]^ were used for refinement of the structures. Data collection and refinement statistics are reported in Table 1 and Figures of molecular structures were generated with PyMOL(Schrodinger 2015

**Scheme 1.**
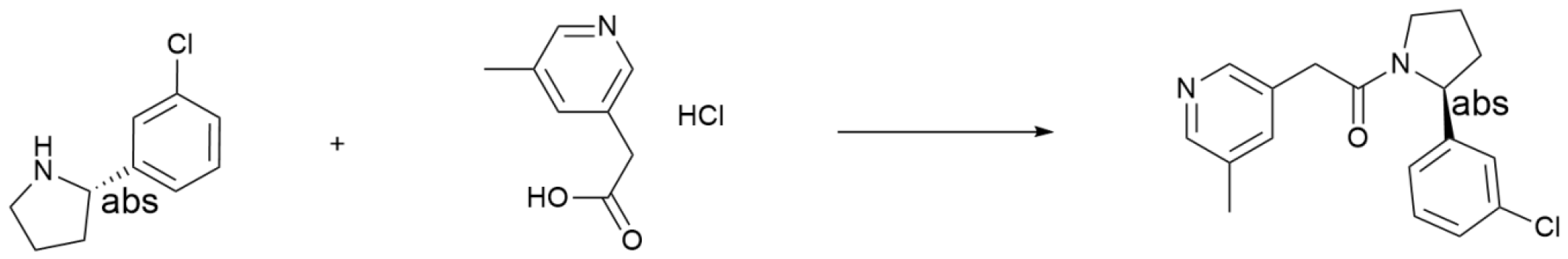
Synthetic route to compound 7. Reagents and conditions: HATU, DIPEA, DMF, RT

**Scheme 2.**
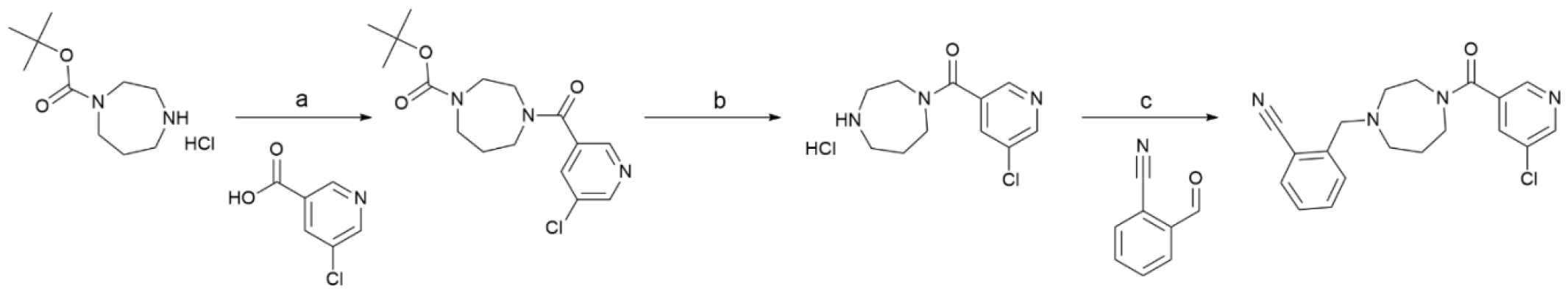
Synthetic route to compound 8. Reagents and conditions: (a) HATU, DIPEA, DMF, RT; (b) HCl, MeOH, RT; (c) NaBH(OAc)3, DIPEA, DCM, RT

**Scheme 3.**
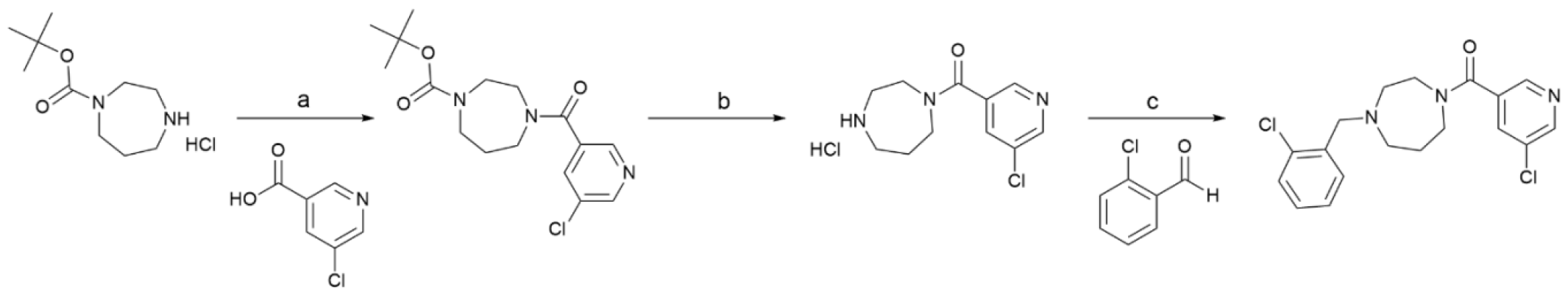
Synthetic route to compound 11. Reagents and conditions: (a) HATU, DIPEA, DMF, RT; (b) HCl, MeOH, RT; (c) NaBH(OAc)3, DIPEA, DCM, RT

**Scheme 4.**
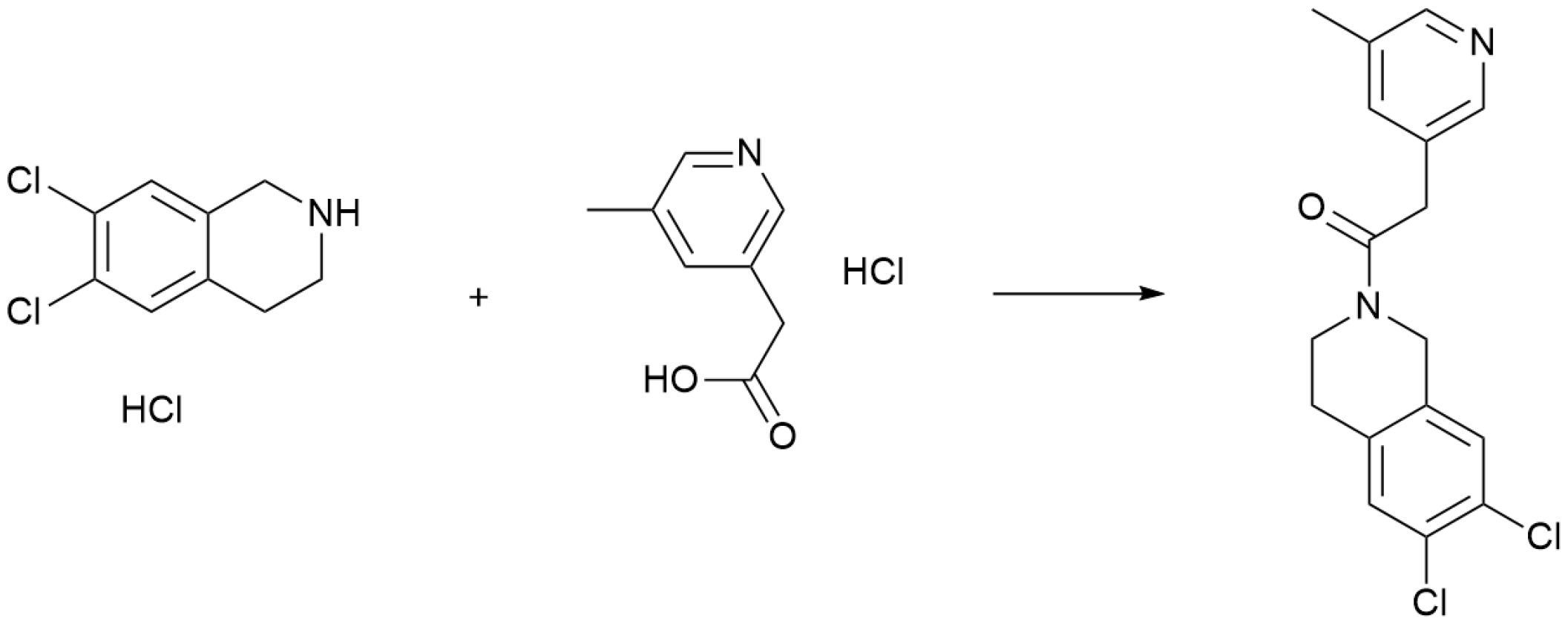
Synthetic route to compound 12. Reagents and conditions: HATU, DIPEA, DMF, RT

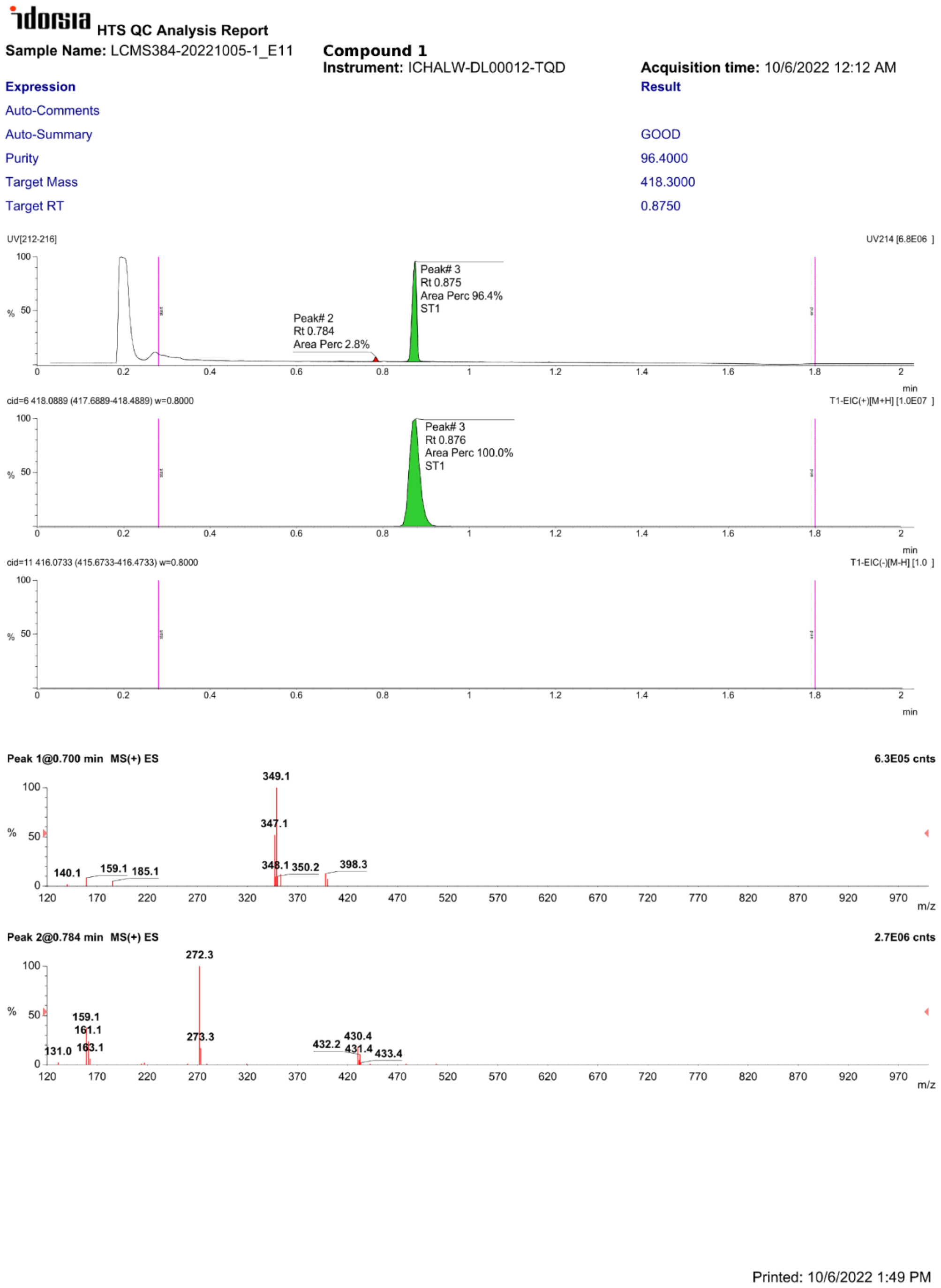

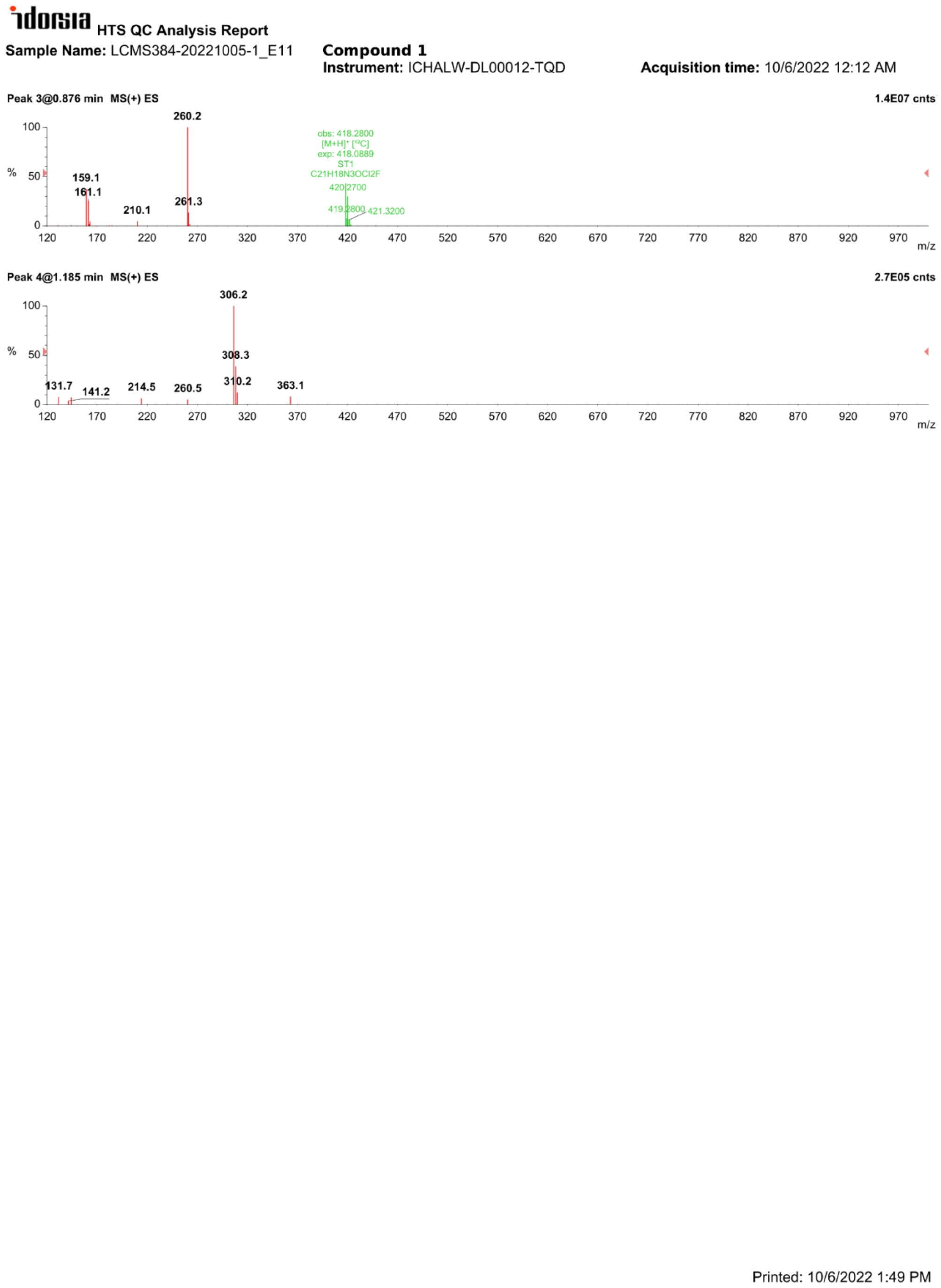

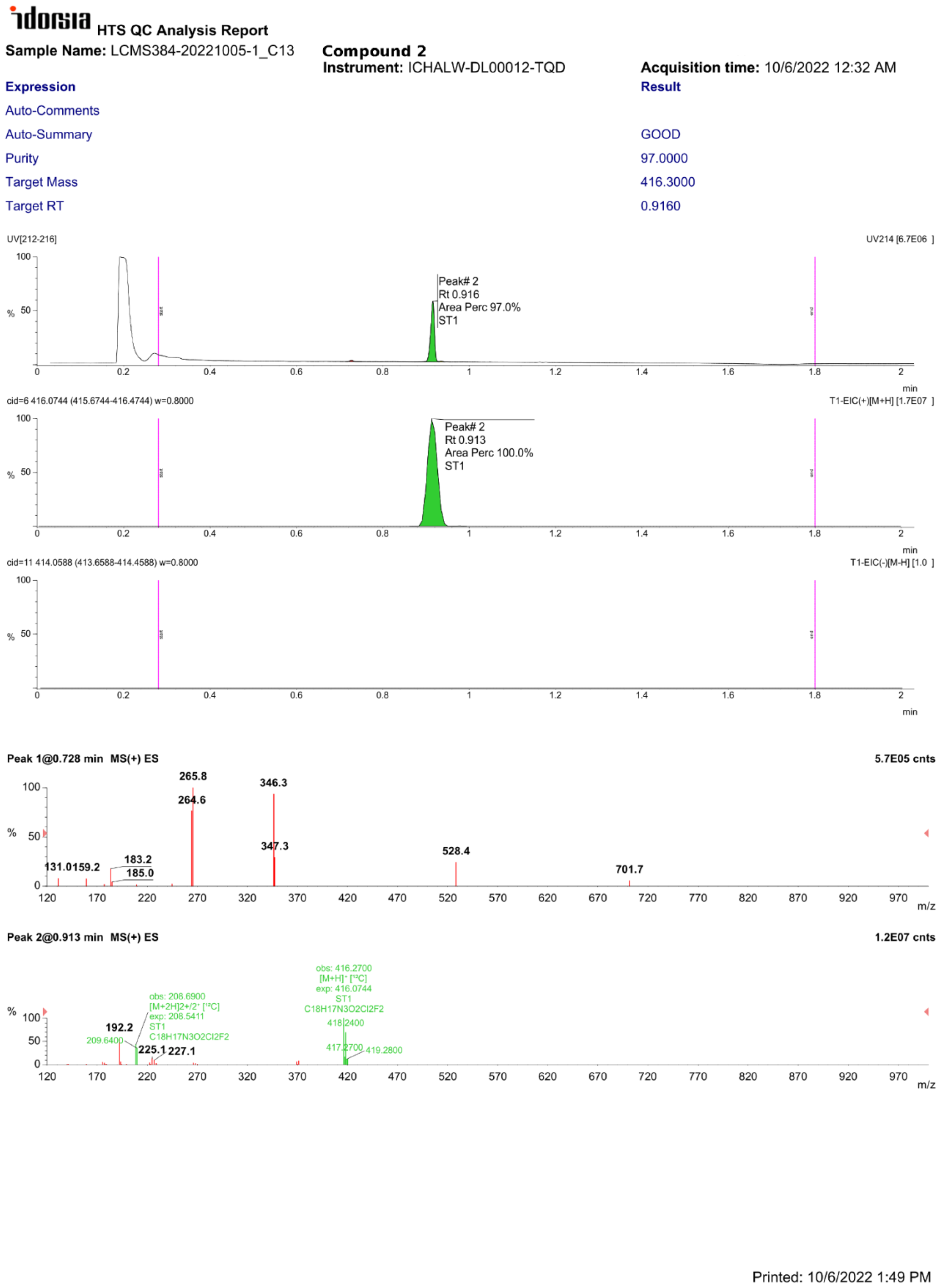

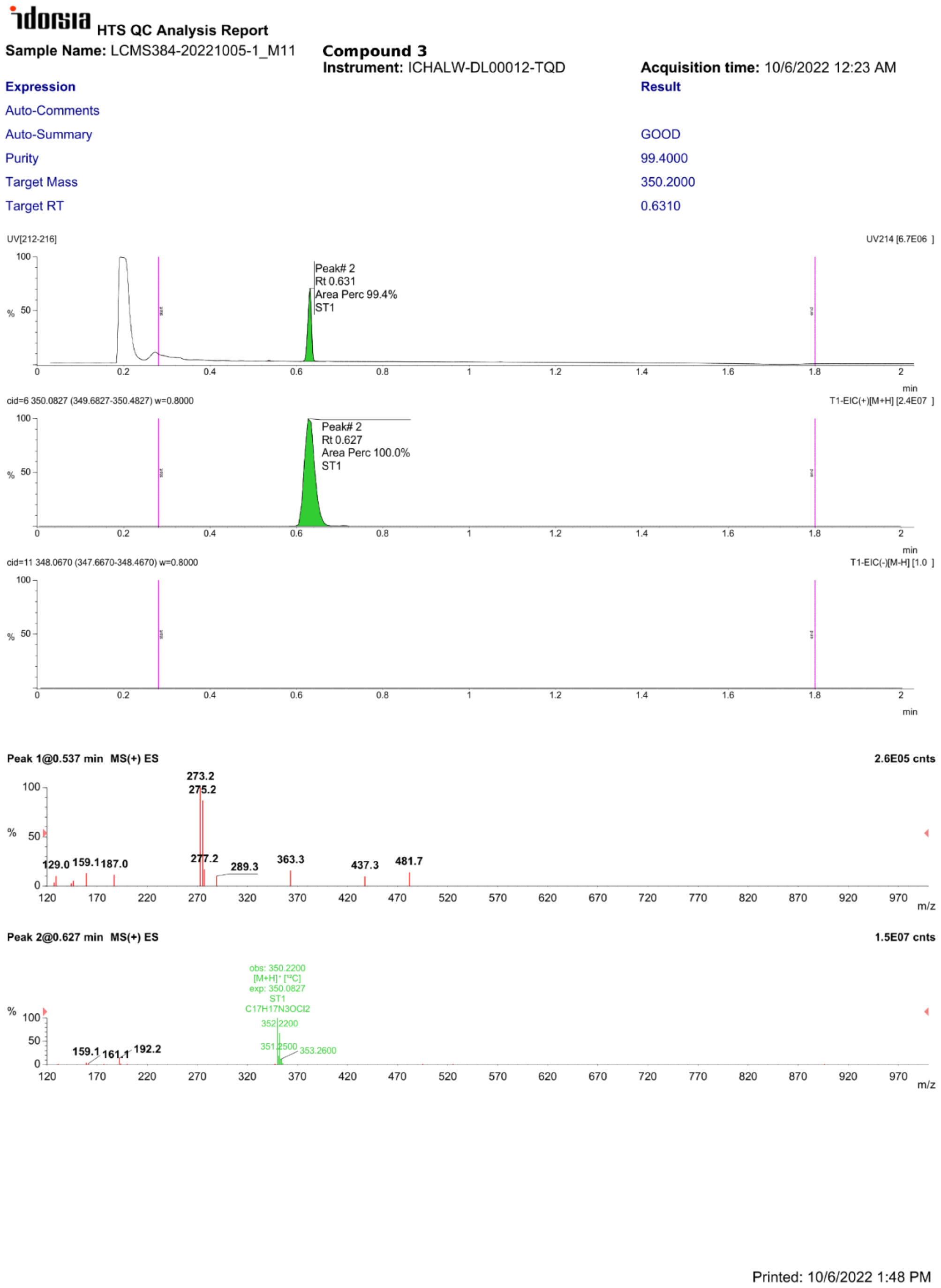

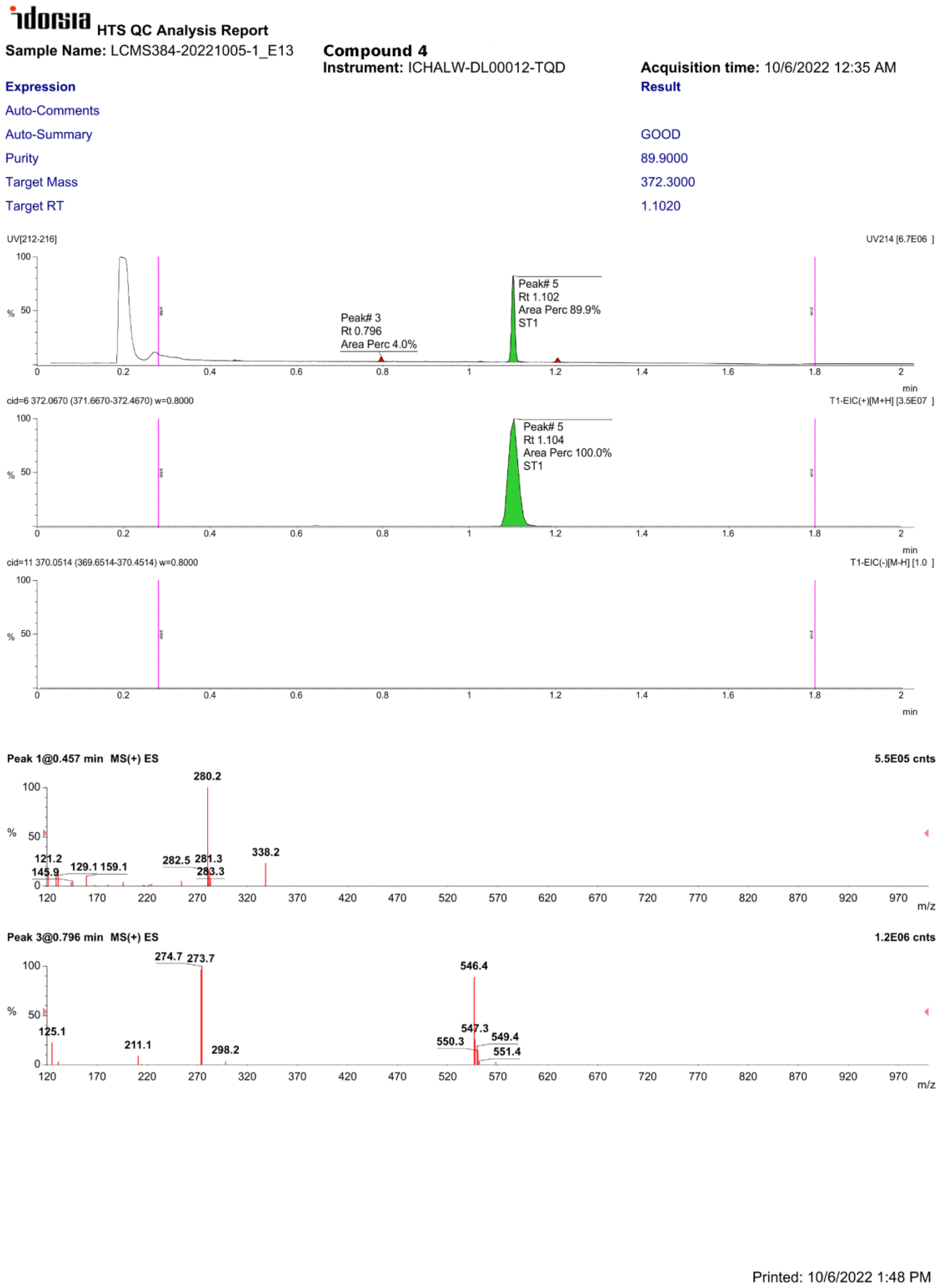

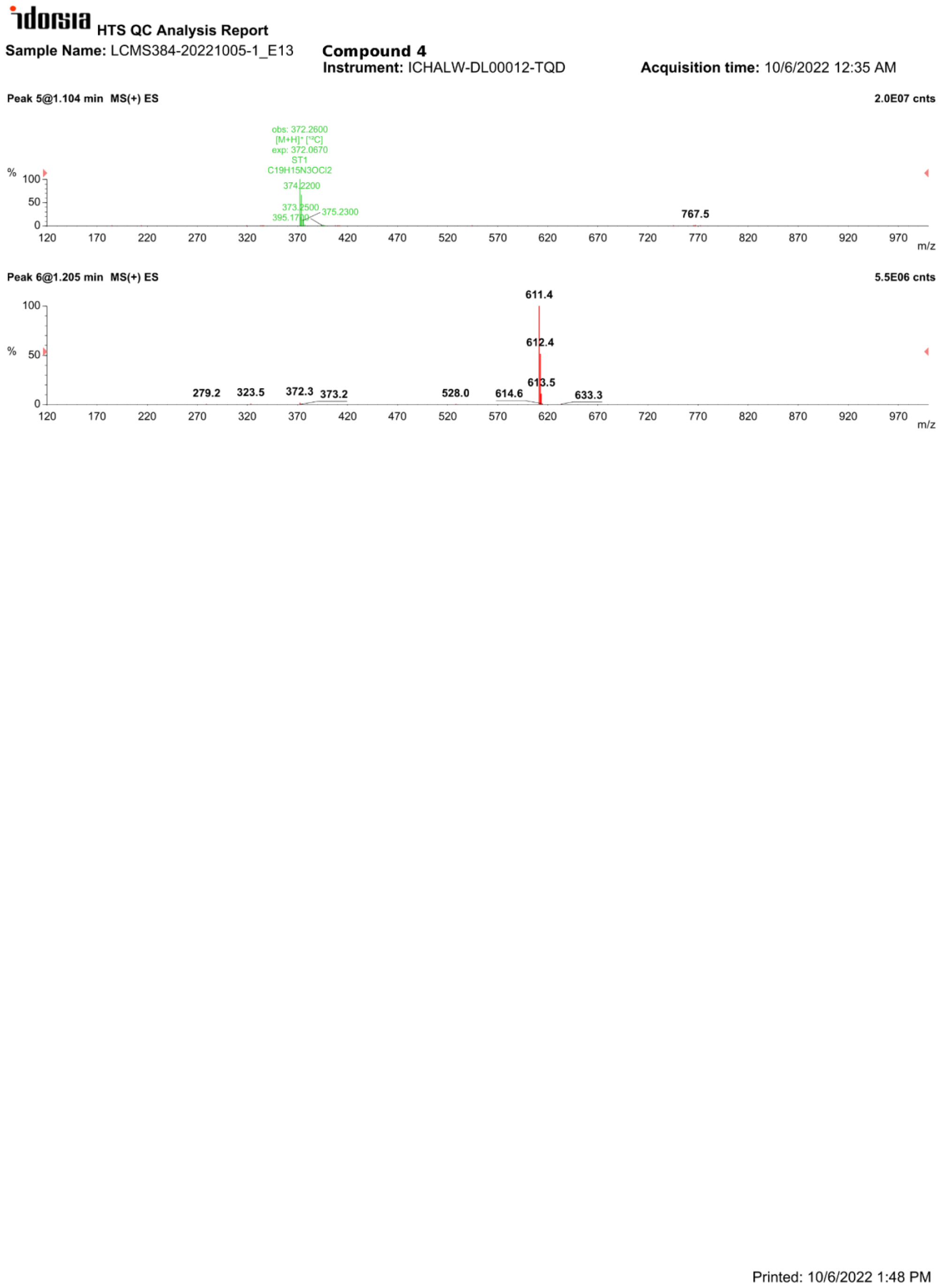

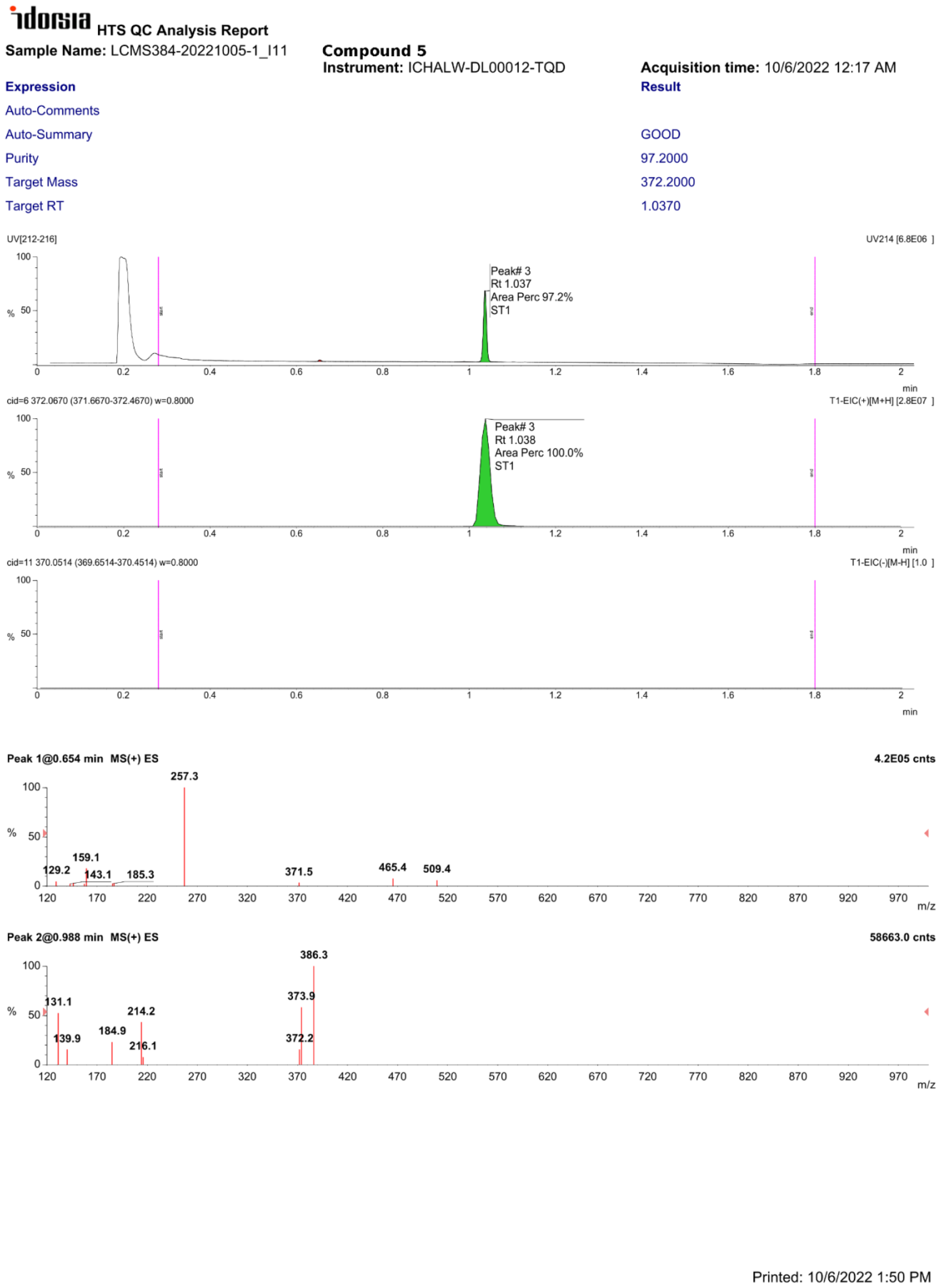

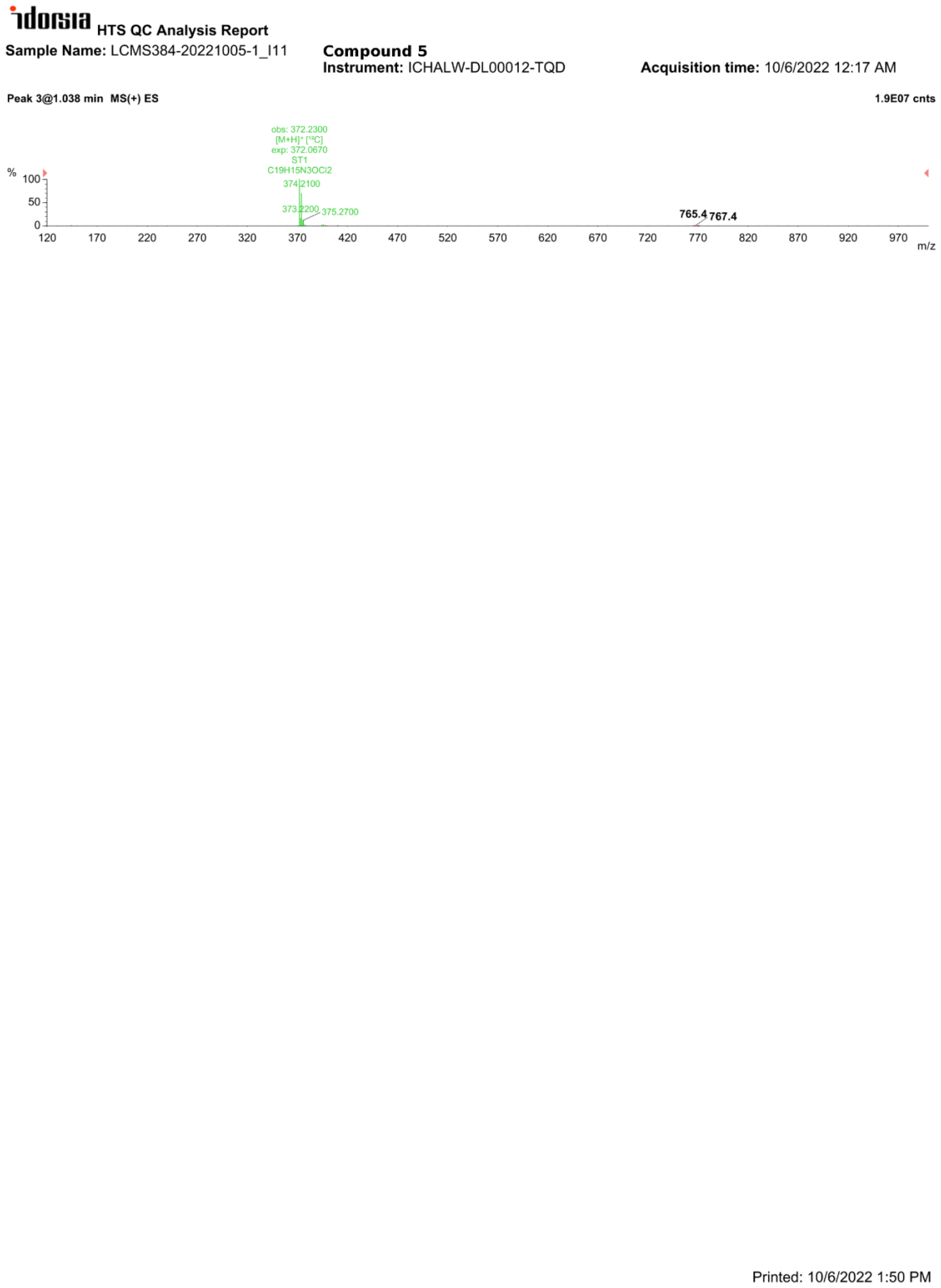

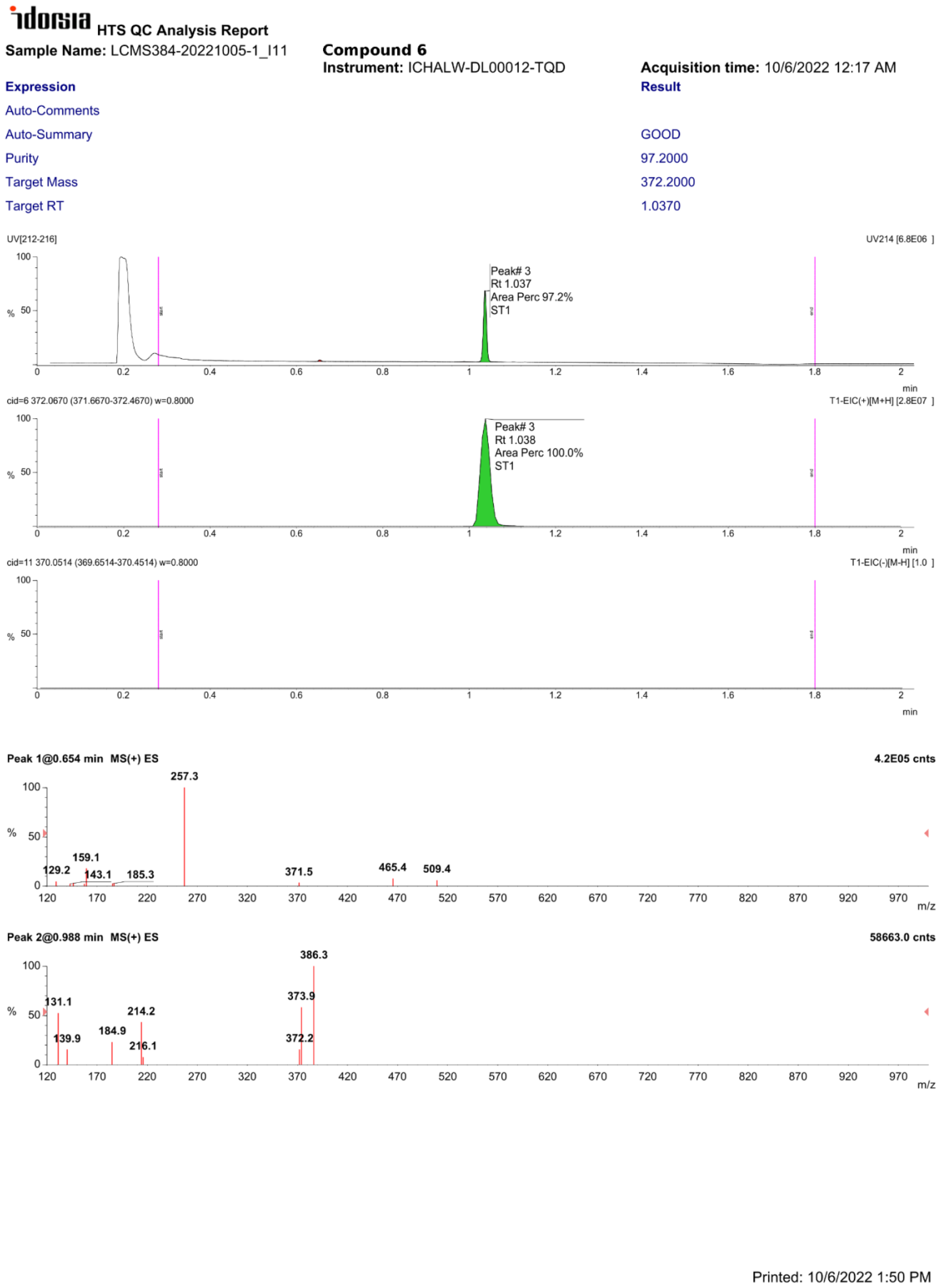

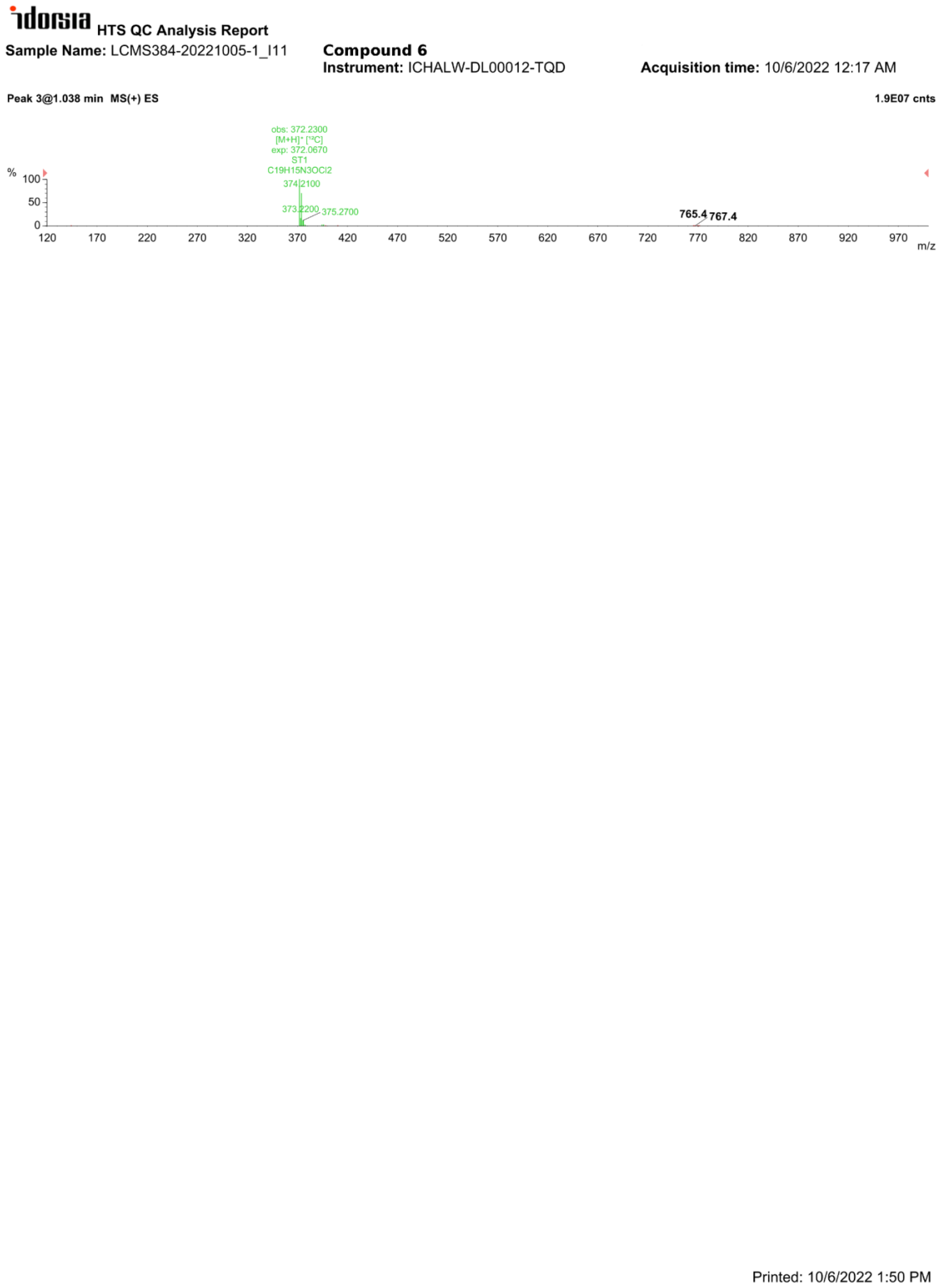

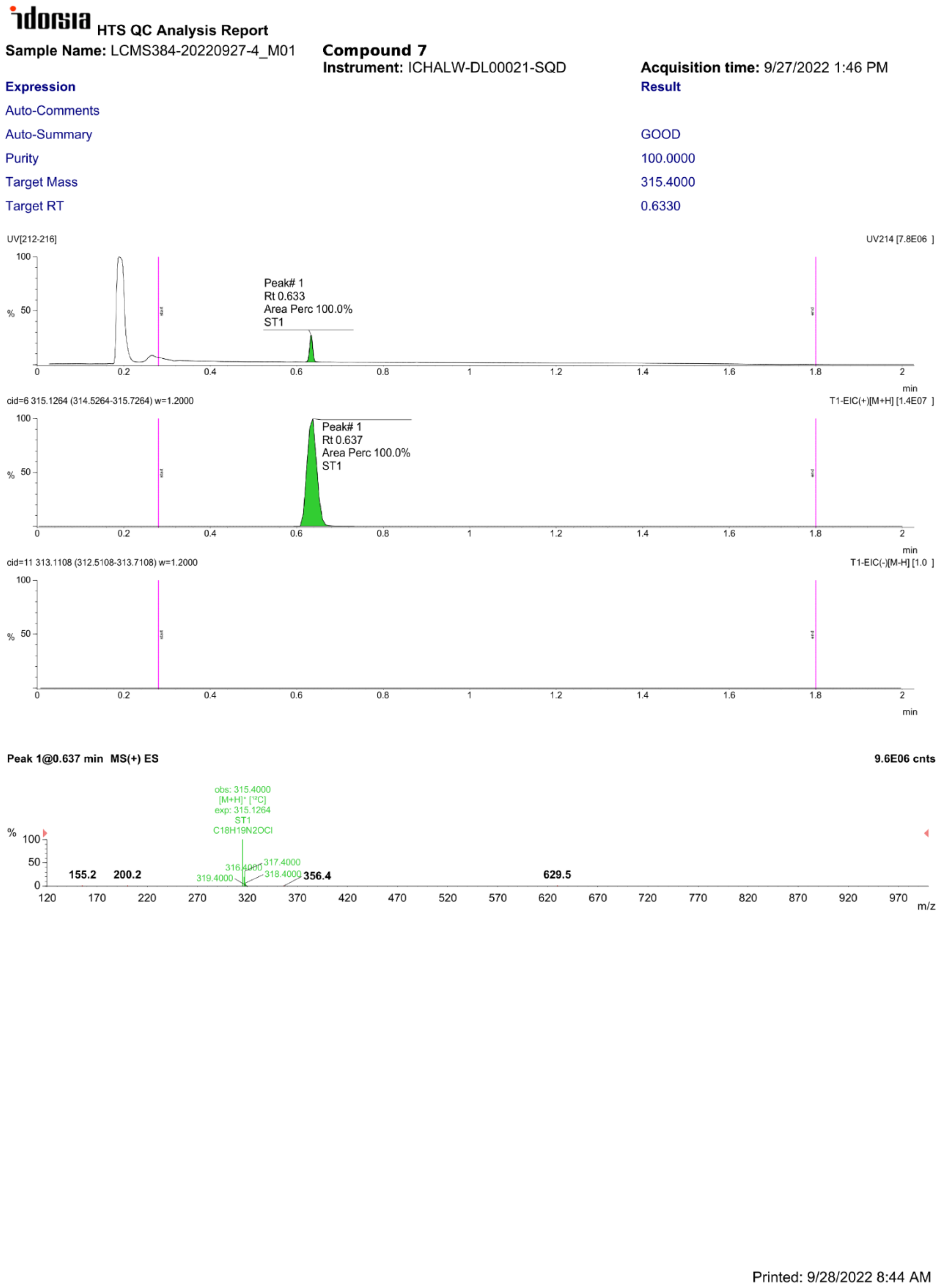

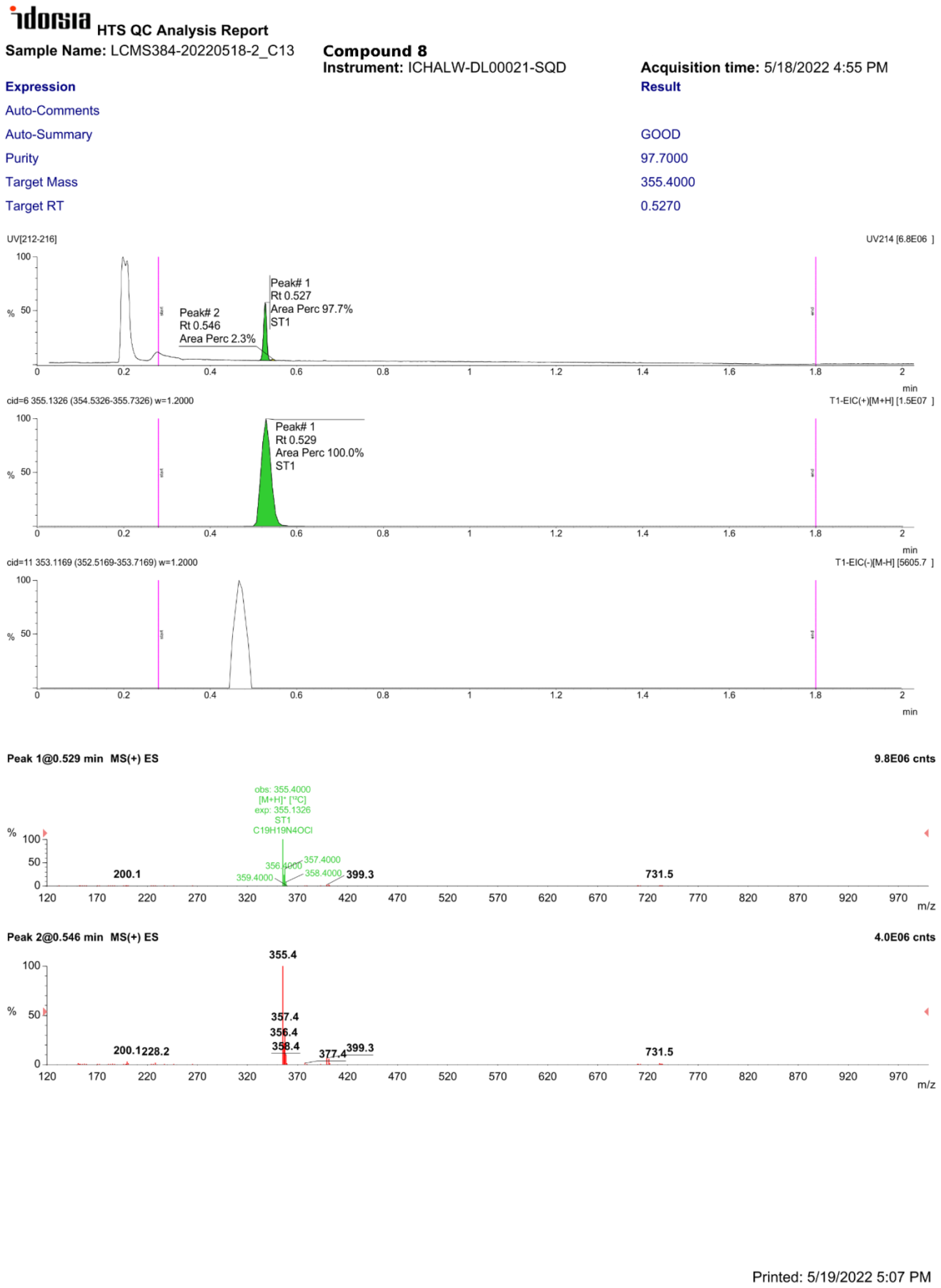

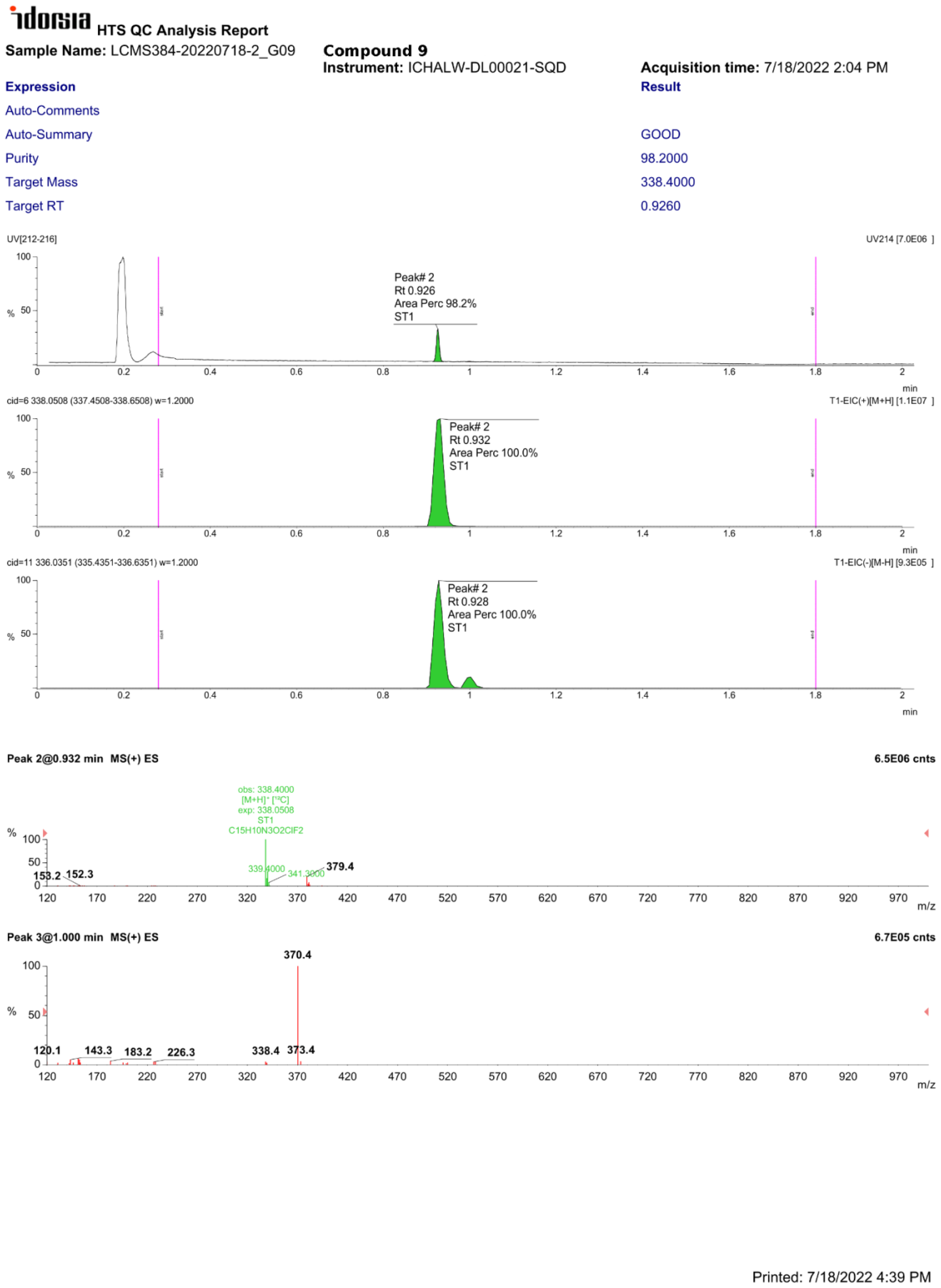

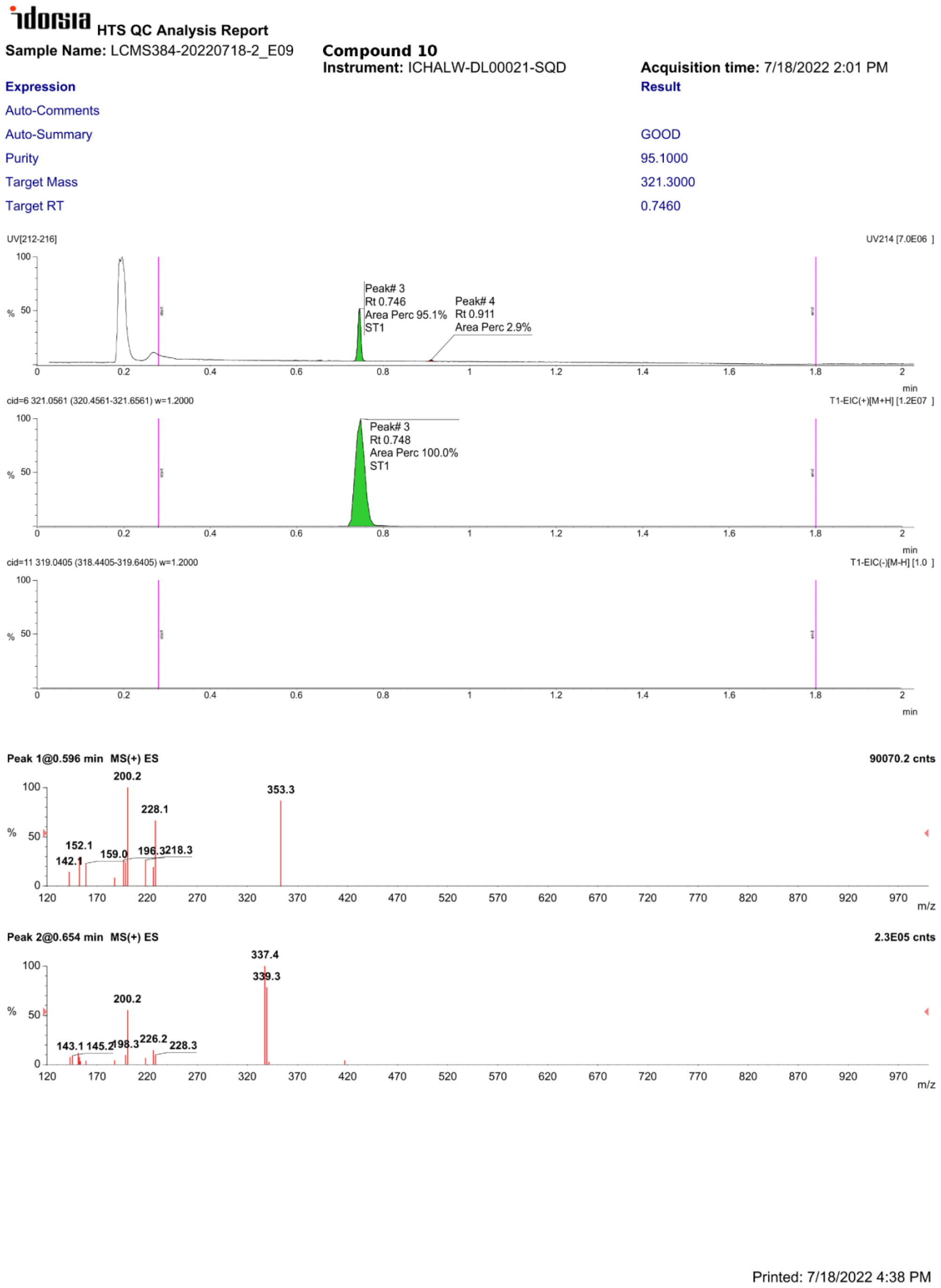

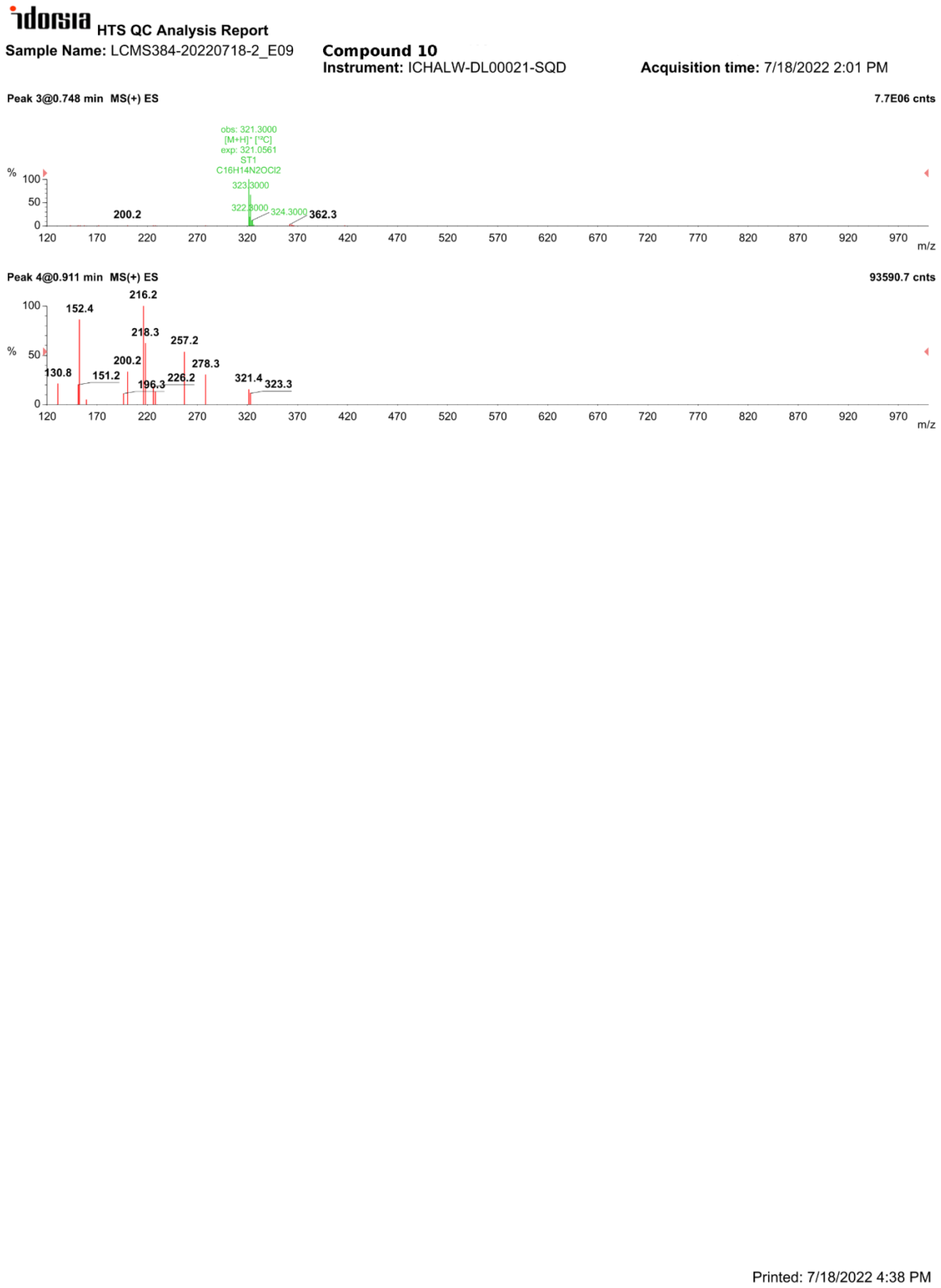

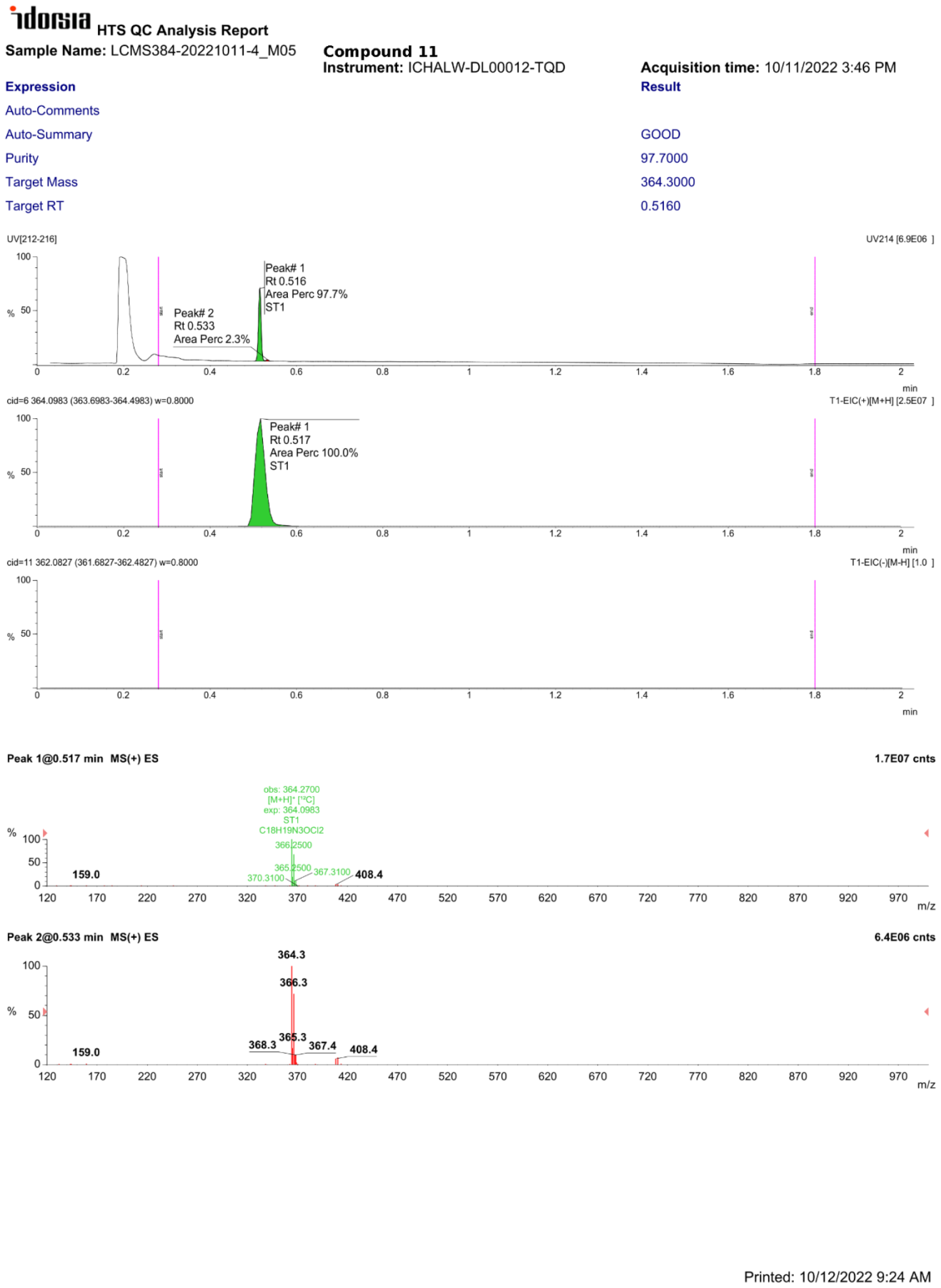

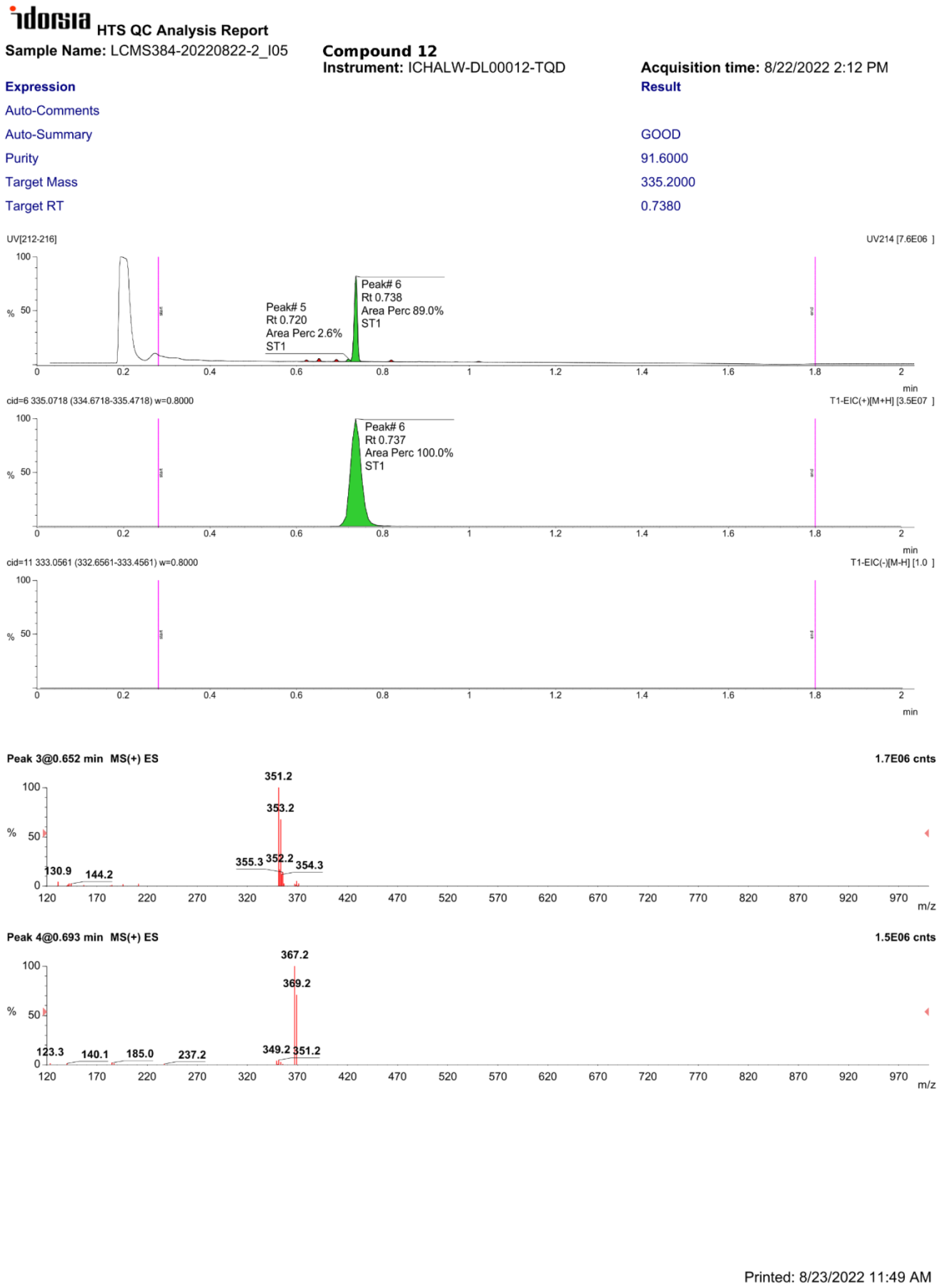

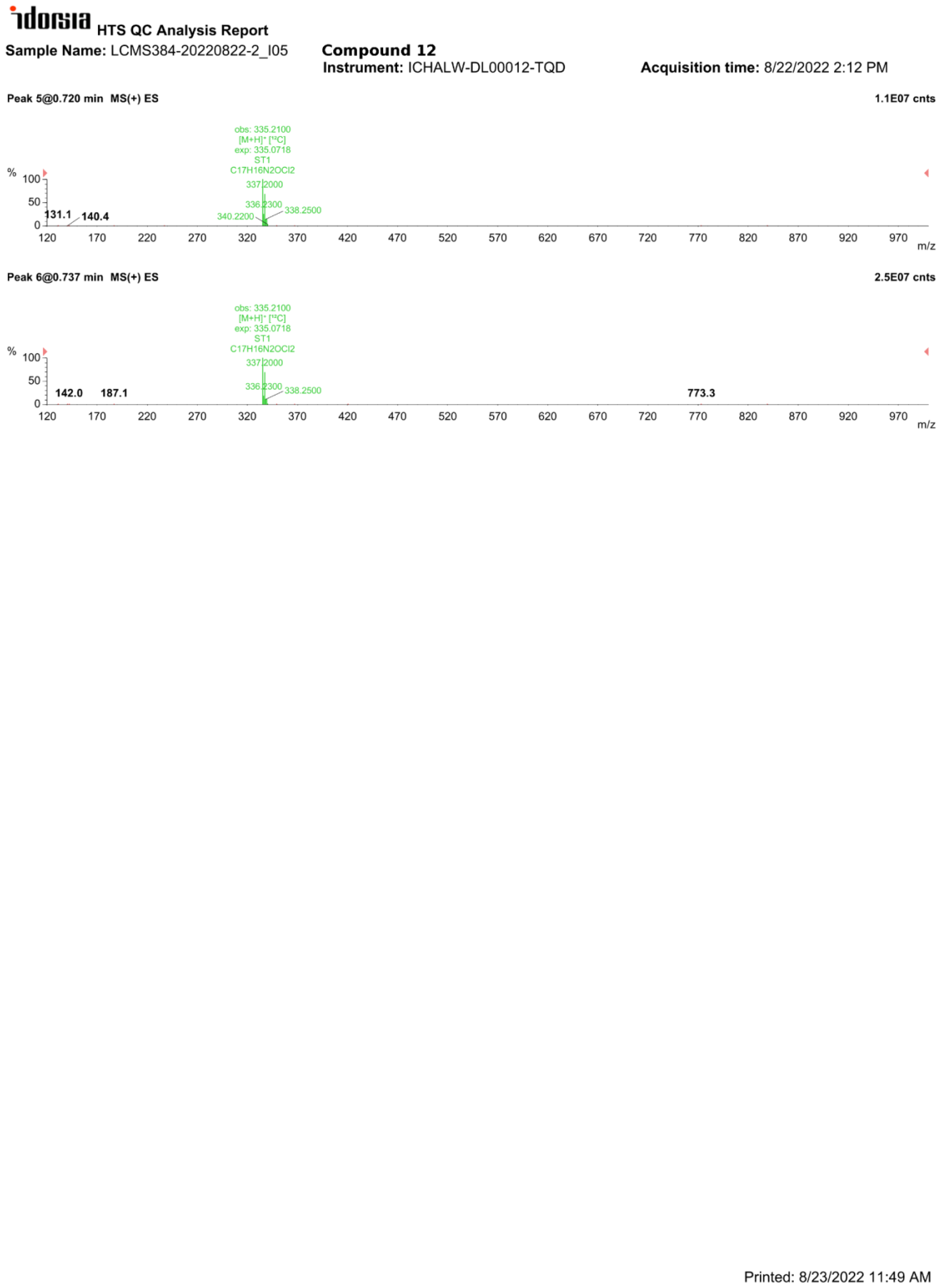

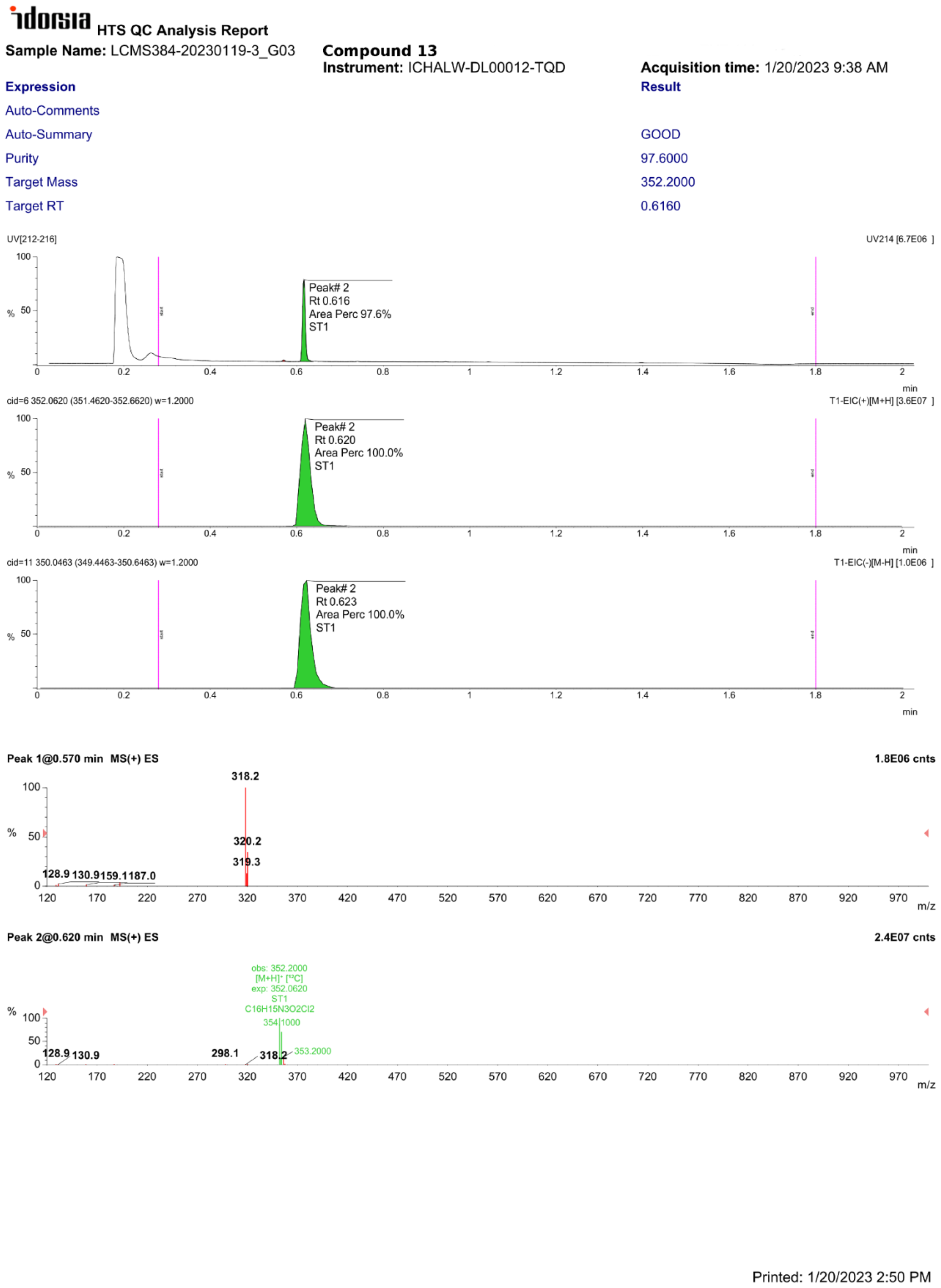

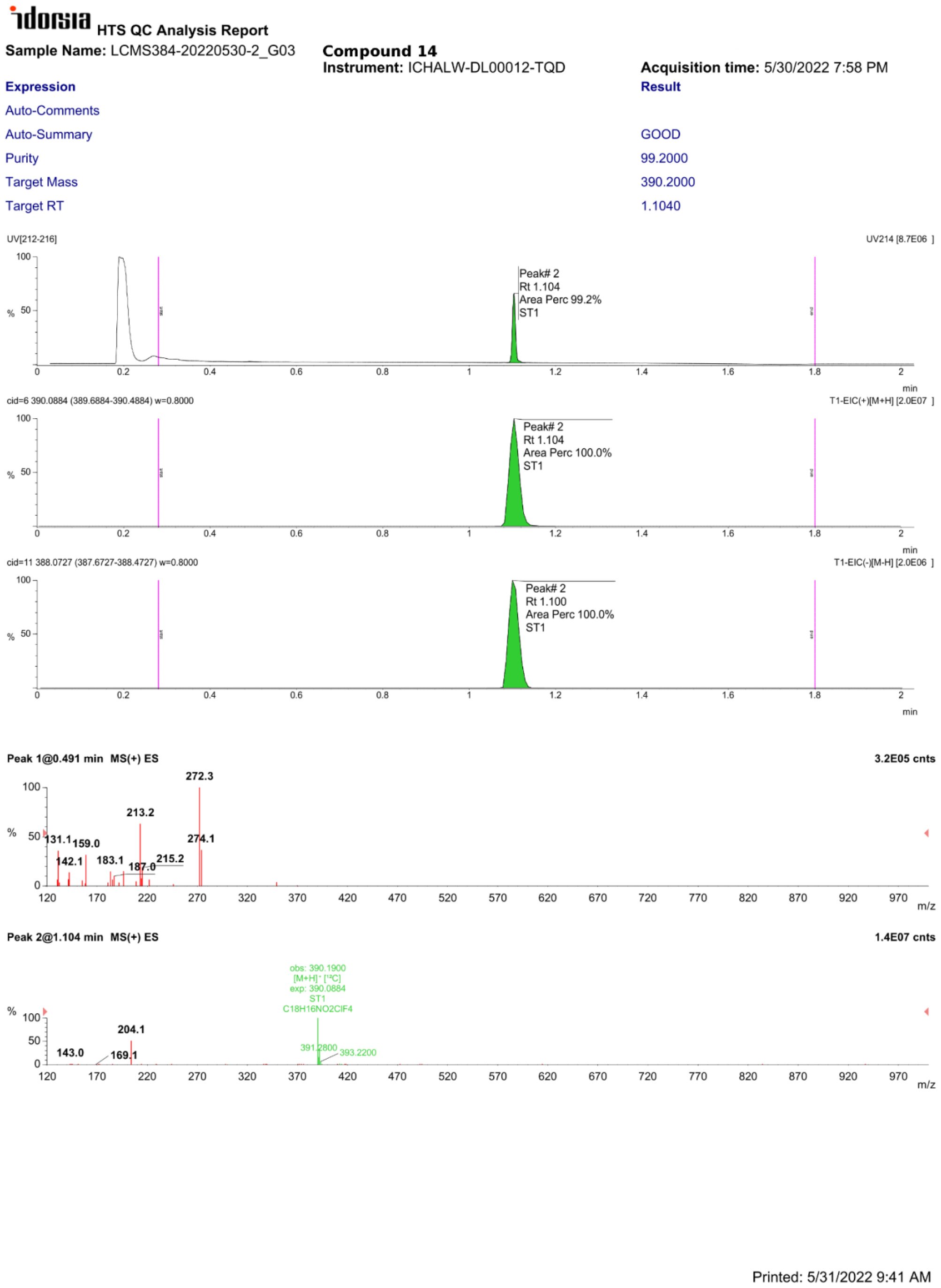

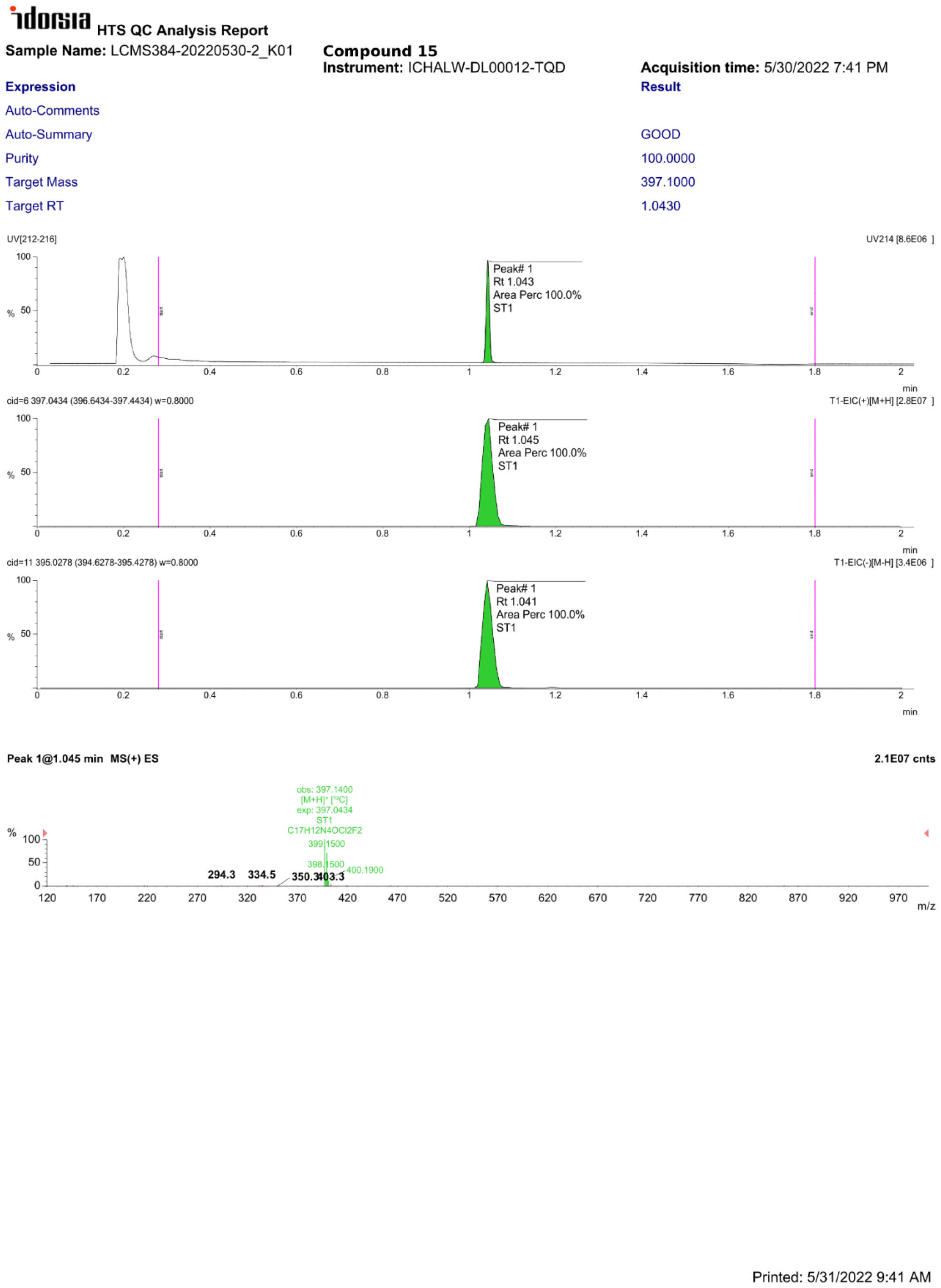

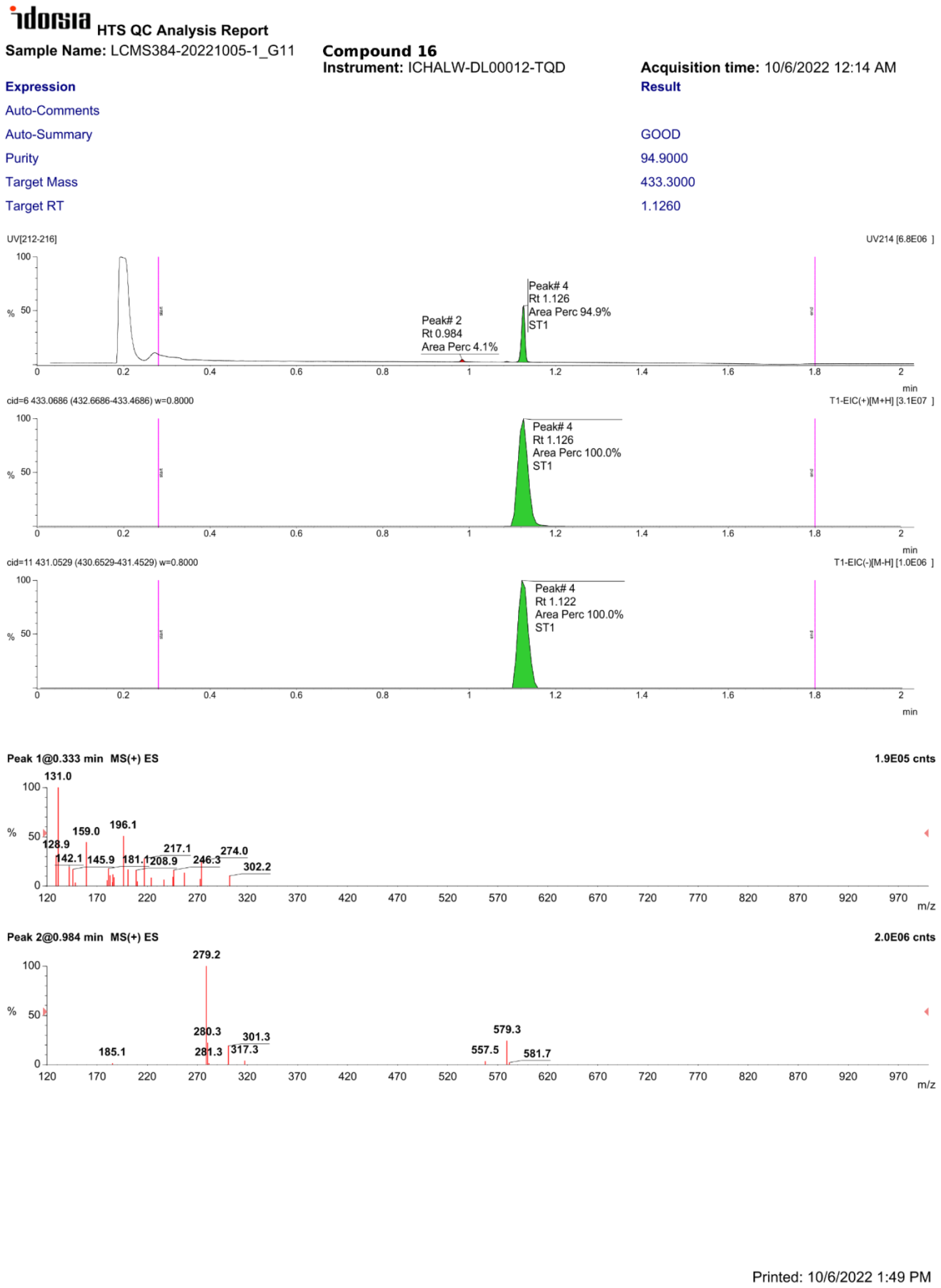

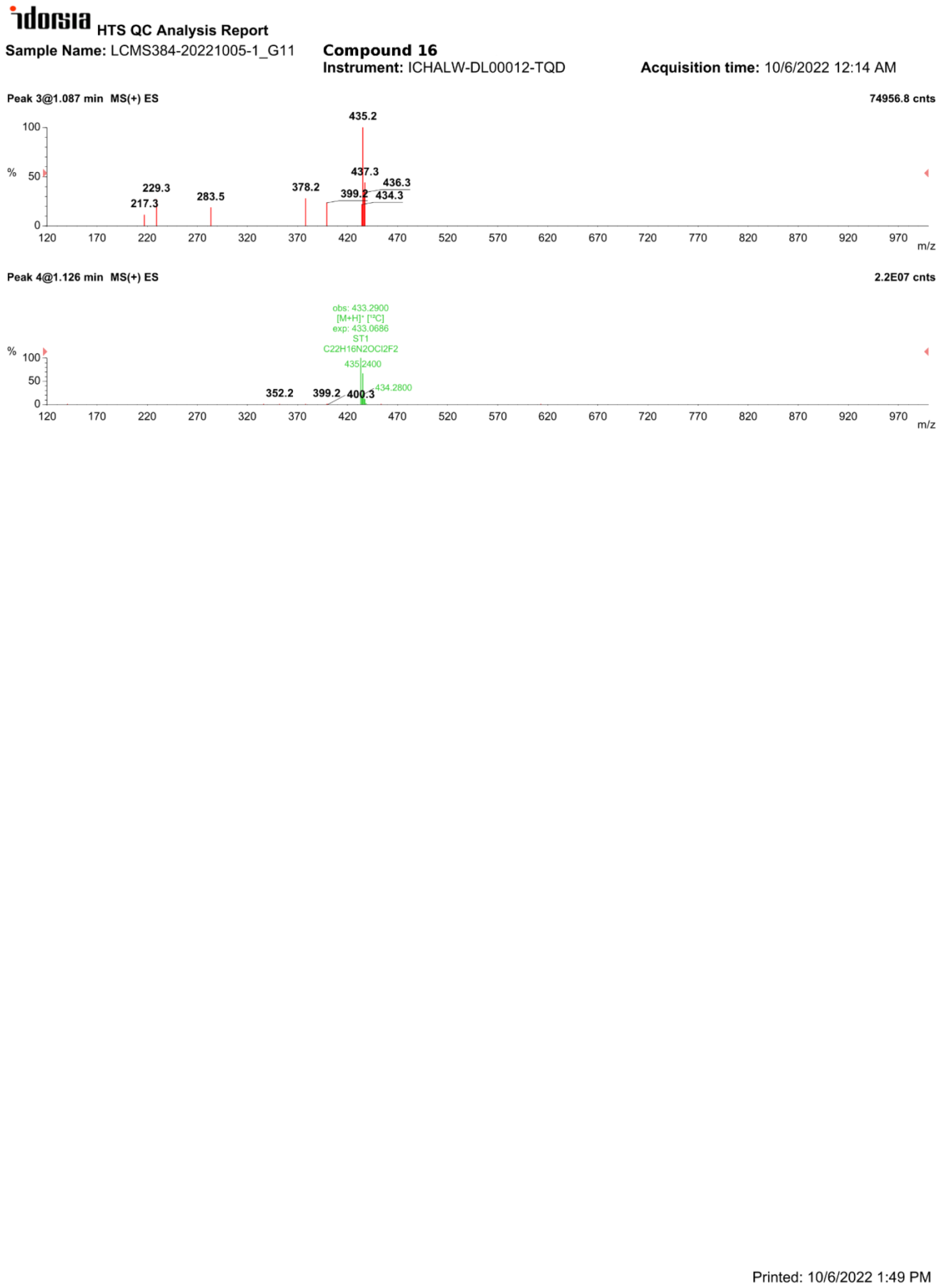

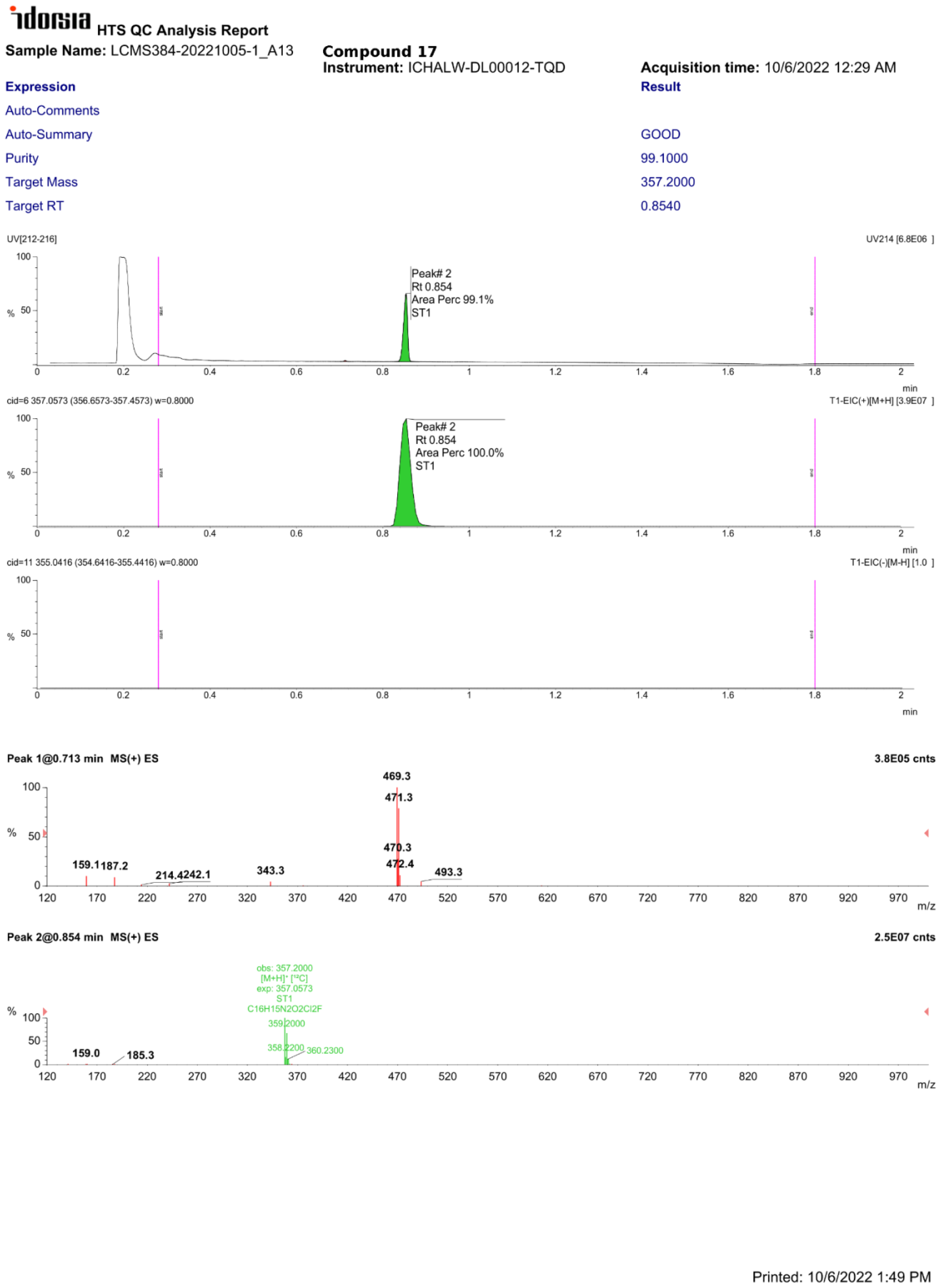

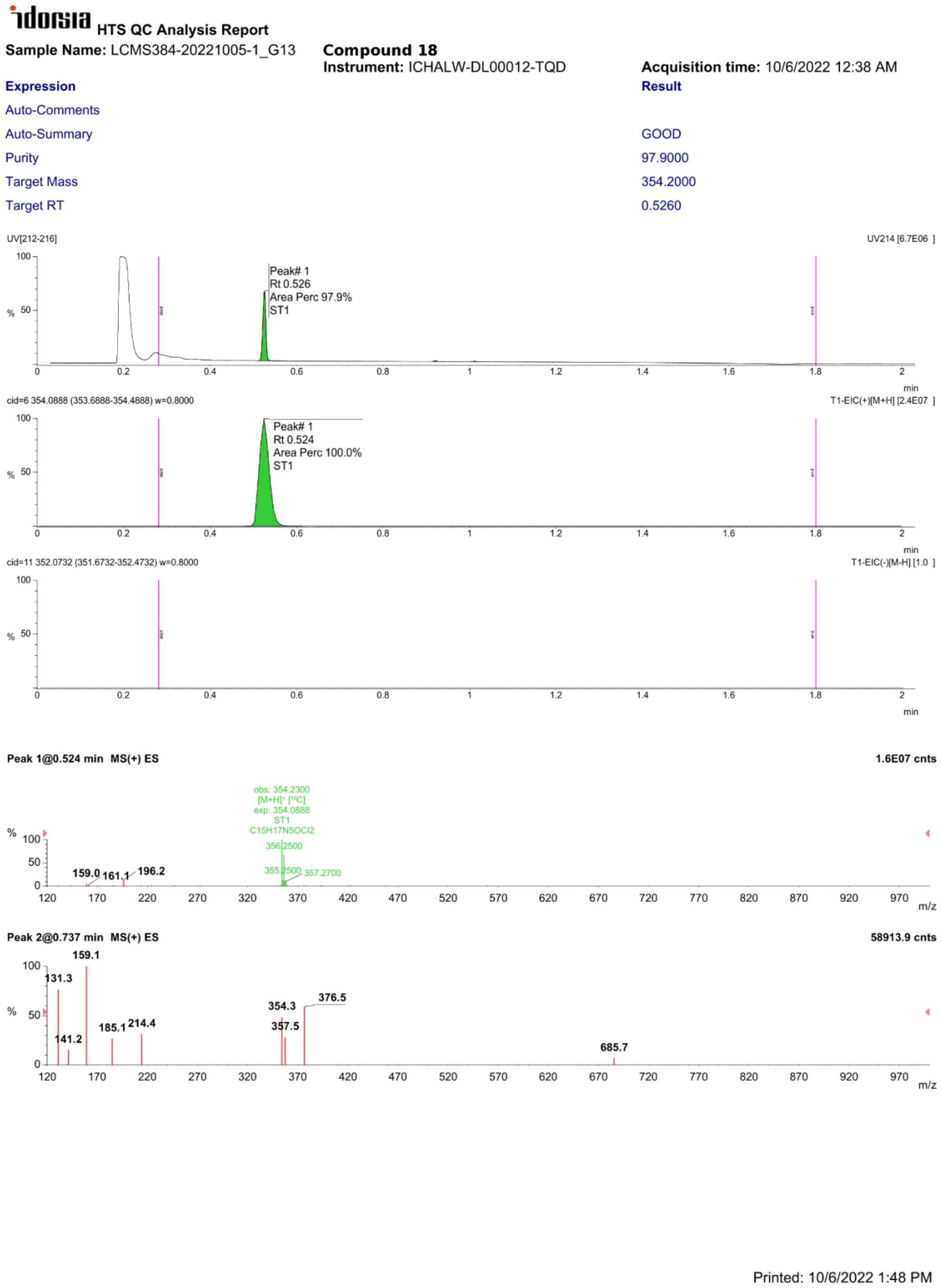

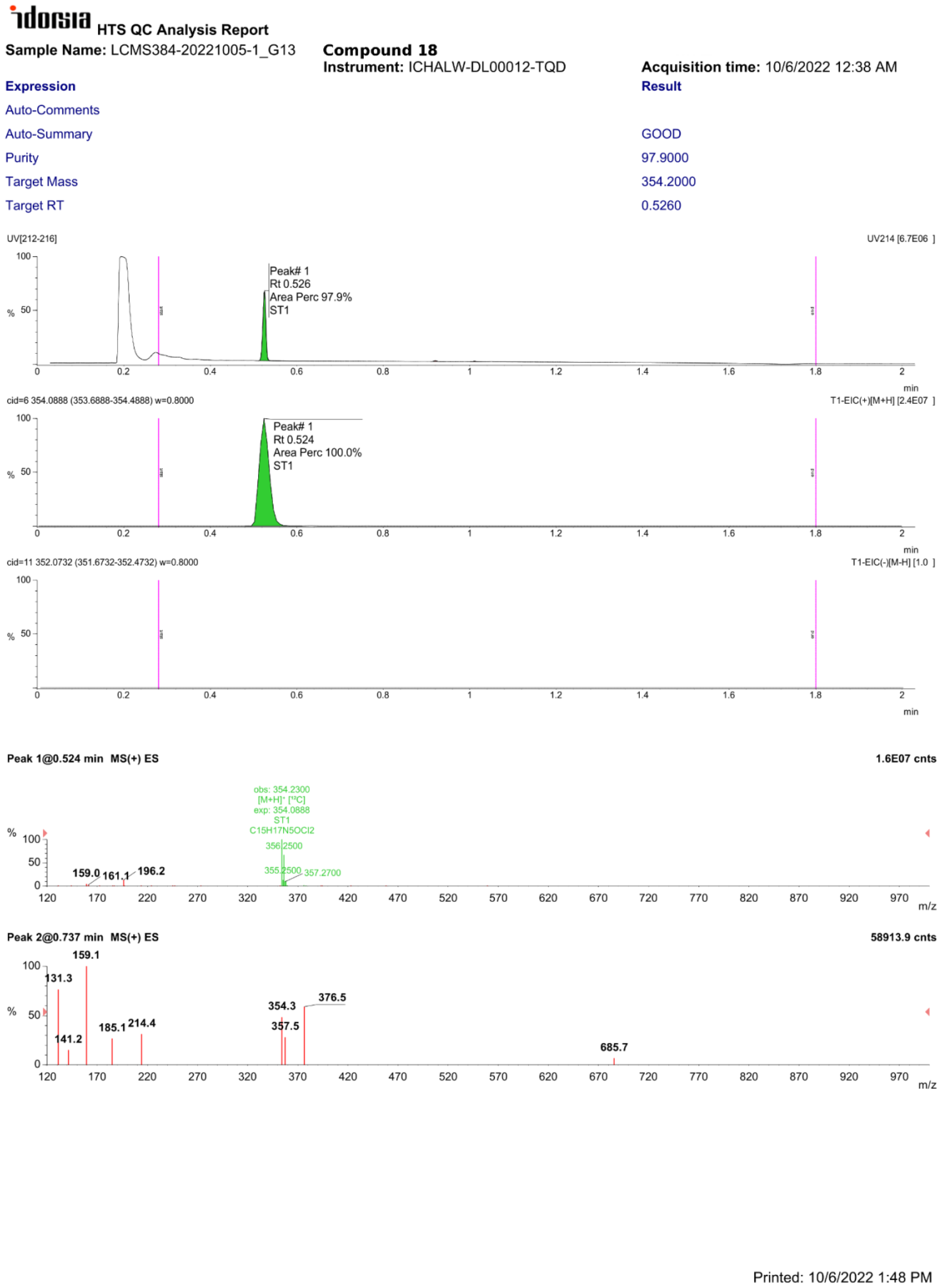

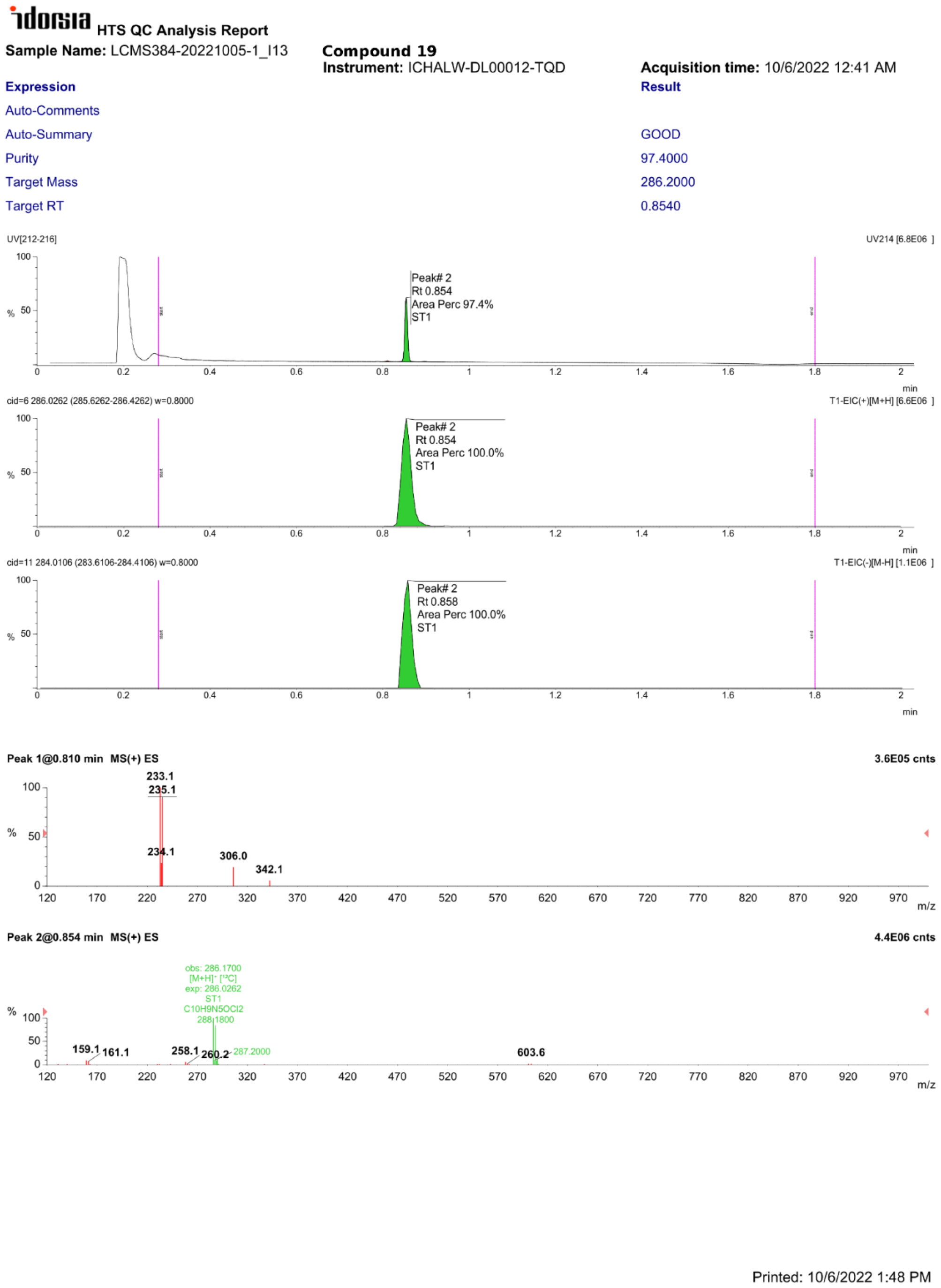

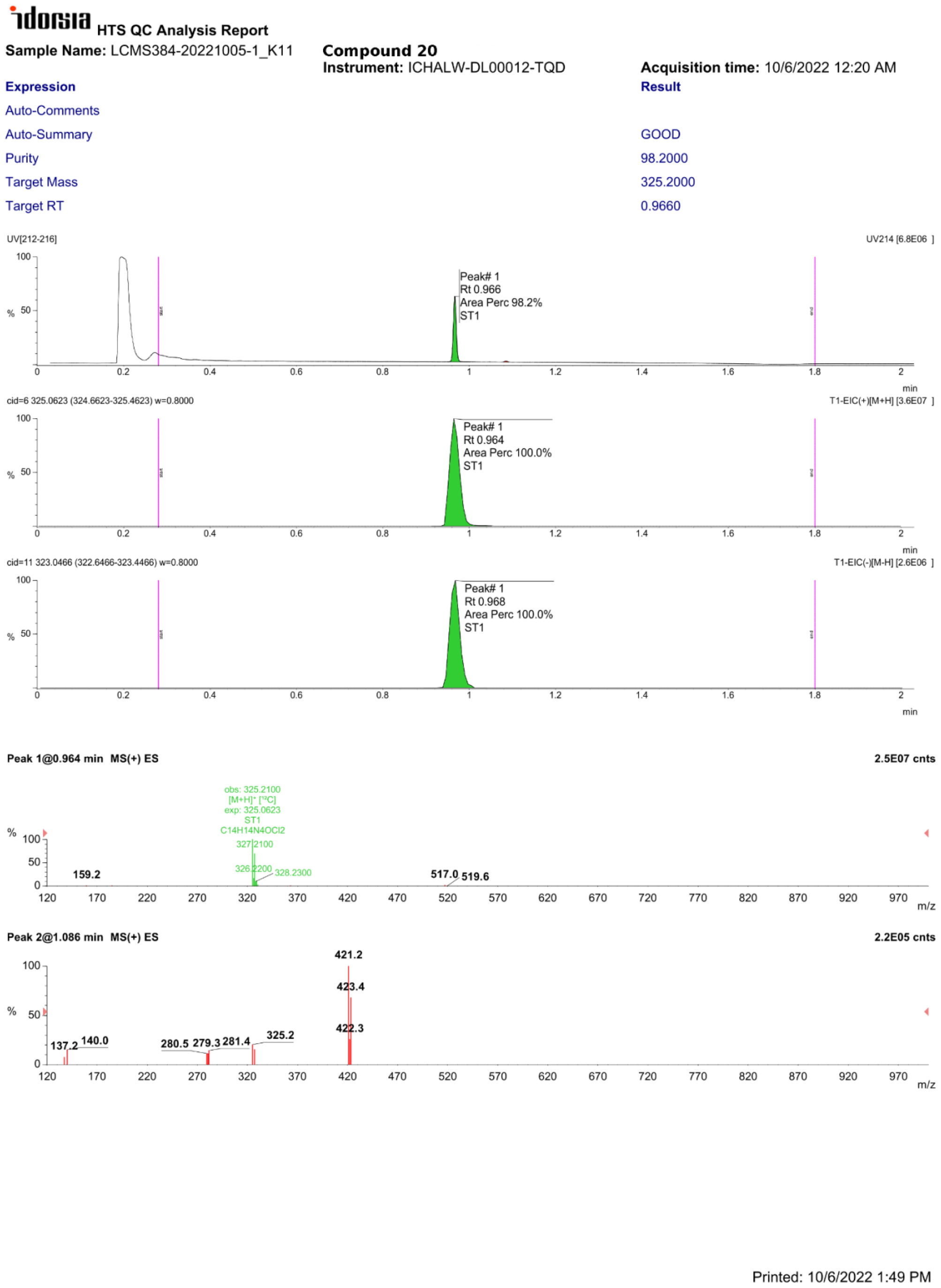

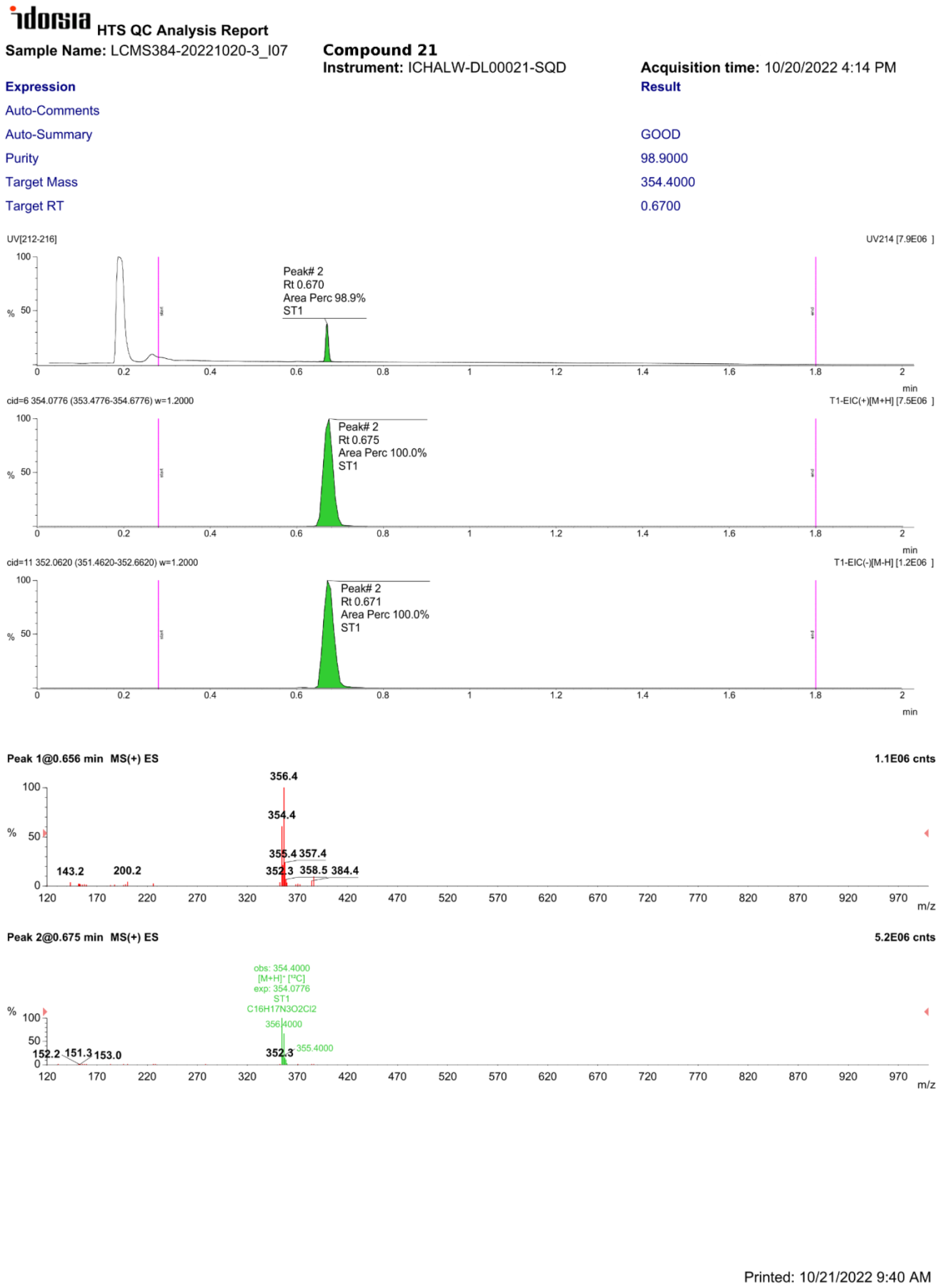

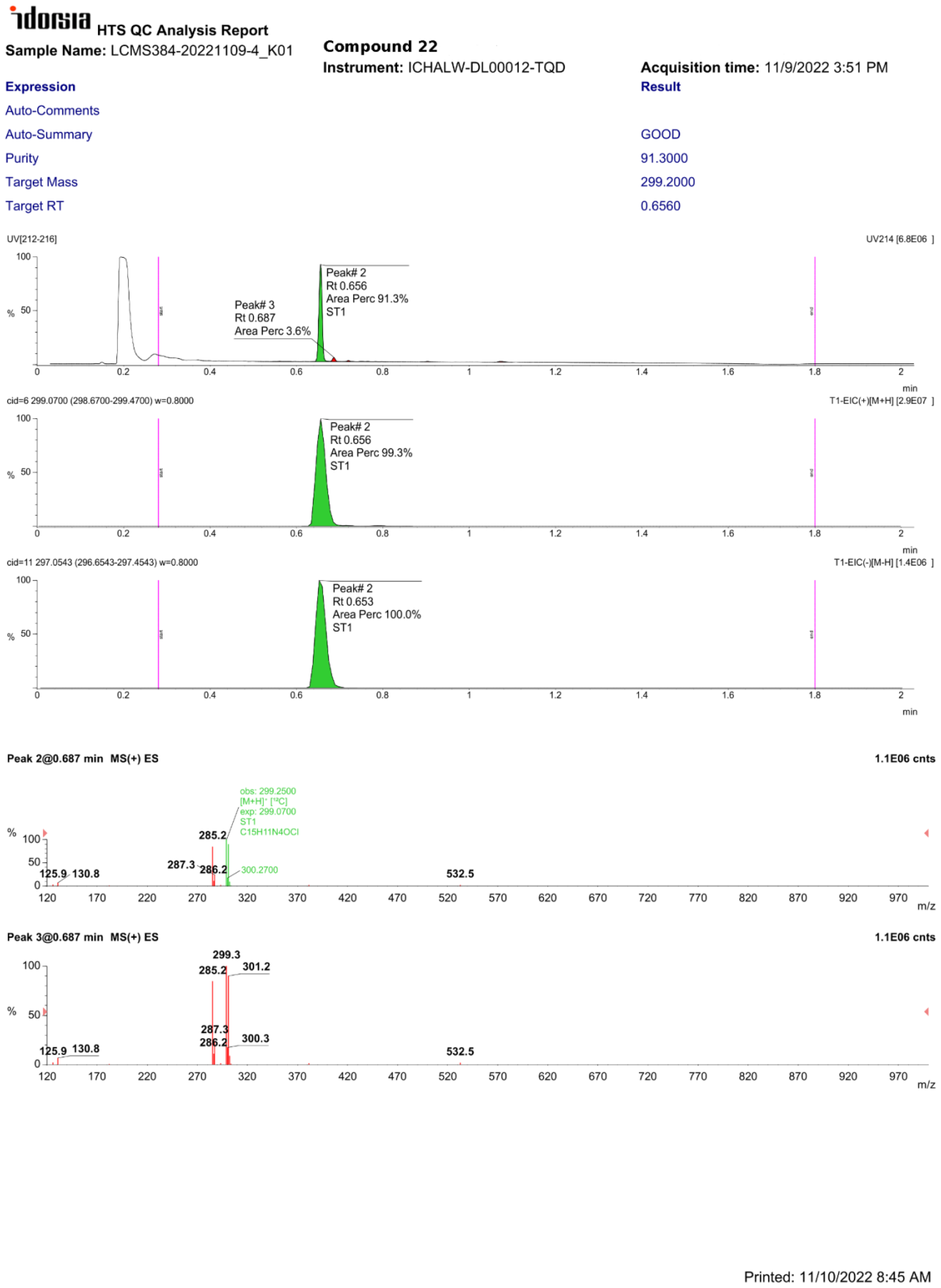

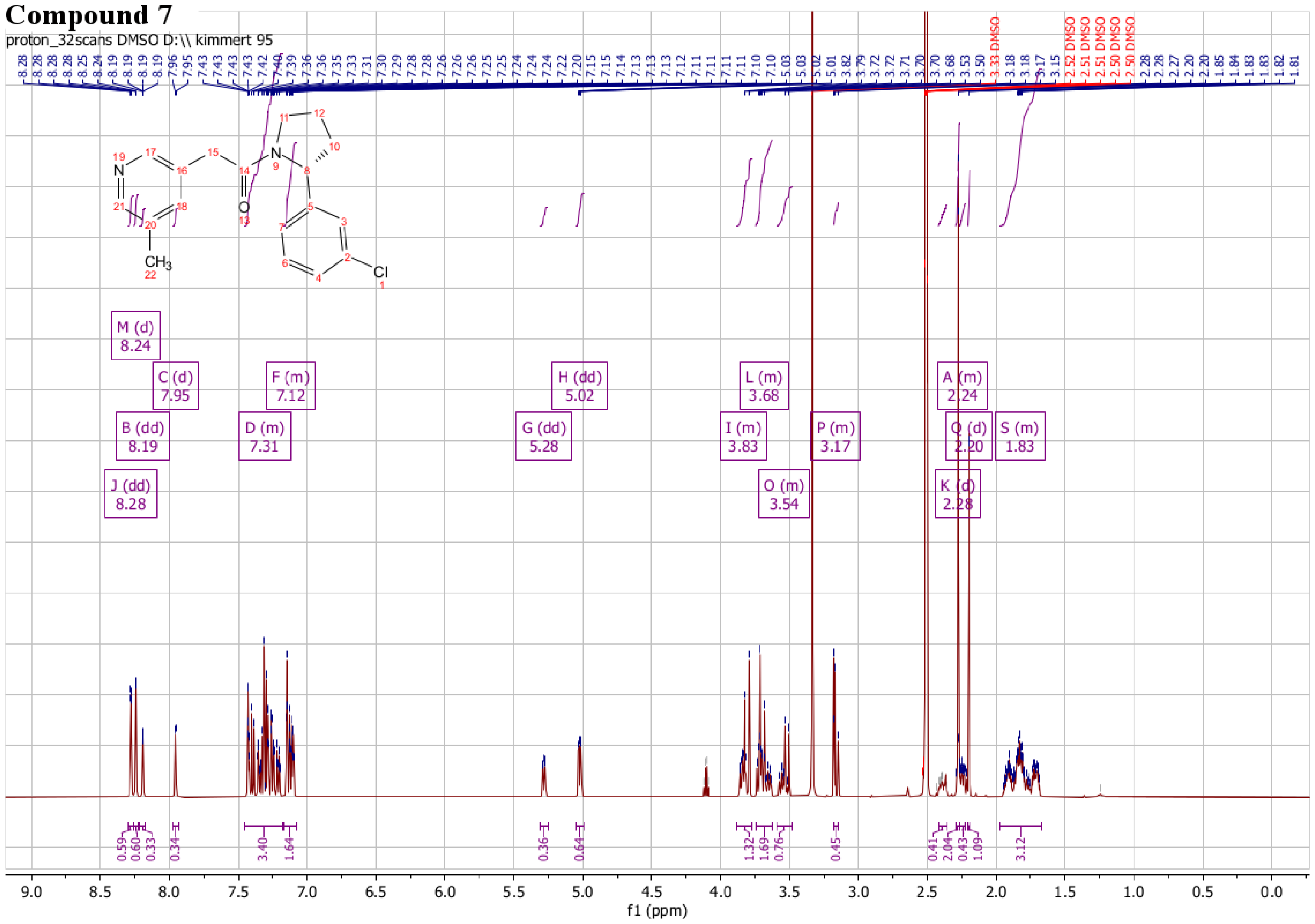

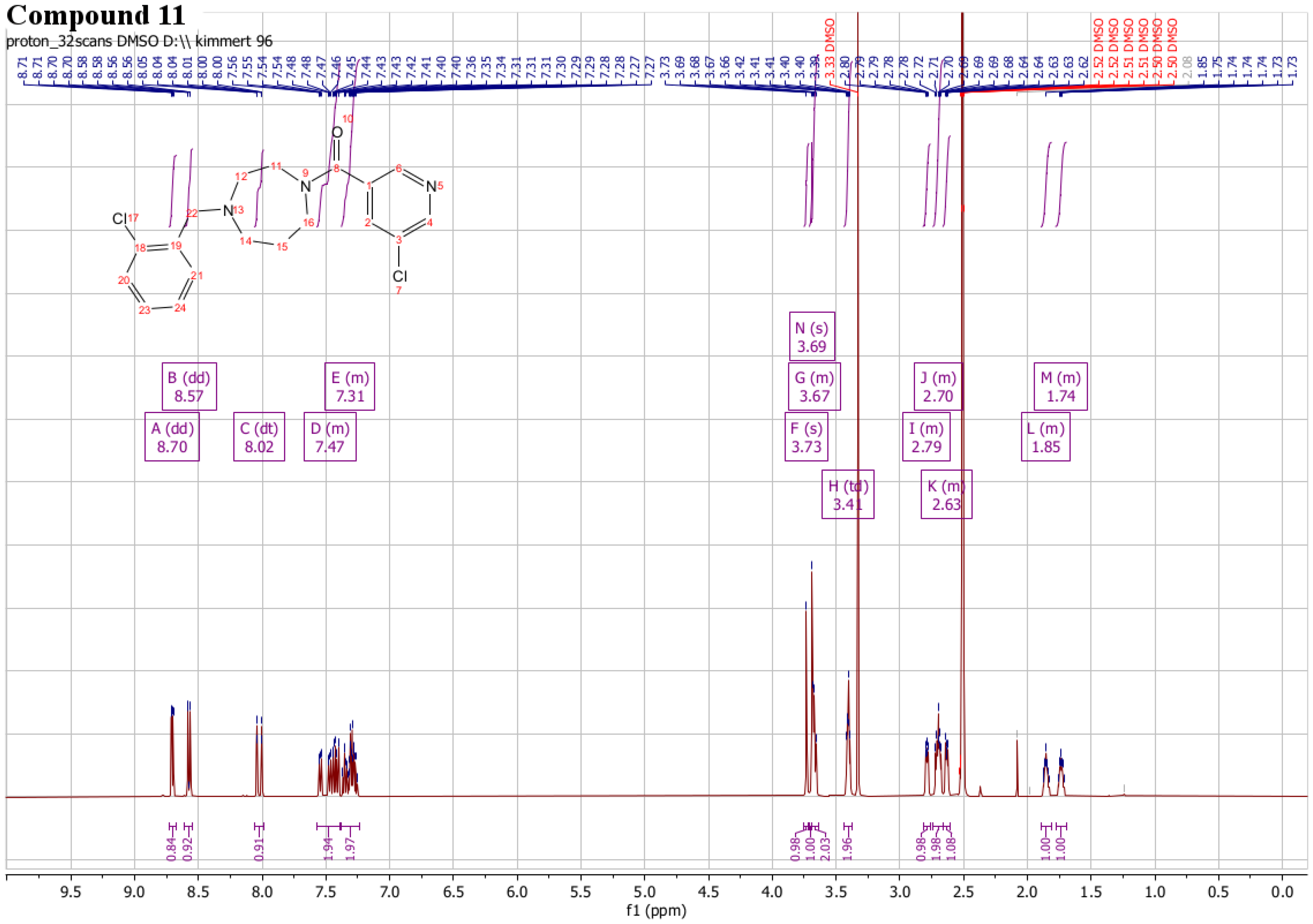

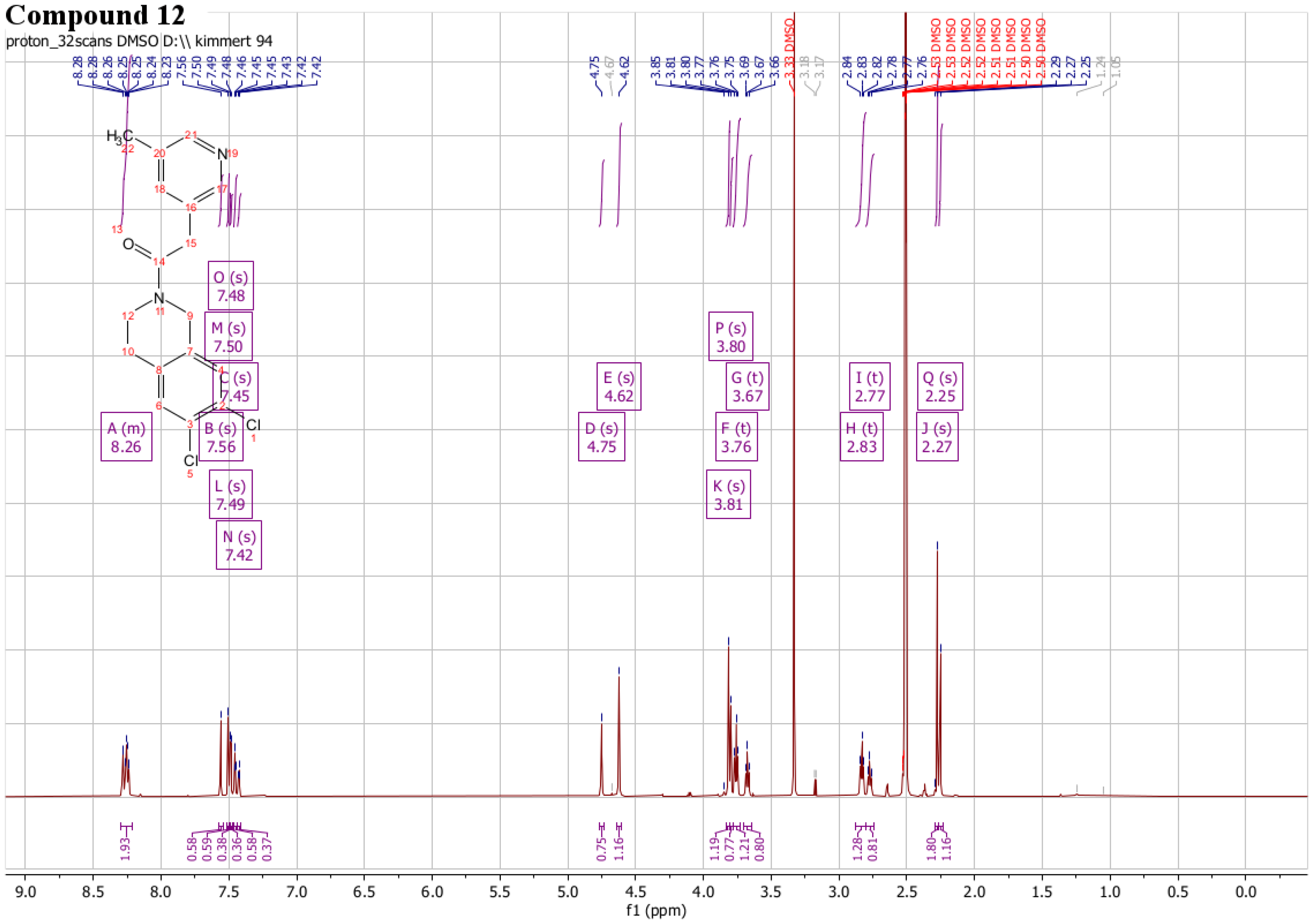

